# Enthalpic and Entropic Contributions to Interleaflet Coupling Drive Domain Registration and Anti-registration in Biological Membrane

**DOI:** 10.1101/2021.09.29.462263

**Authors:** Akshara Sharma, Aniruddha Seal, Sahithya S. Iyer, Anand Srivastava

**Affiliations:** Department of Physics, Indian Institute of Science-Bangalore, C. V. Raman Road, Bangalore, Karnataka 560012, India; School of Chemical Sciences, National Institute of Science Education and Research, Bhubaneswar, Khurda, Odisha 752050, India; Molecular Biophysics Unit, Indian Institute of Science-Bangalore, C. V. Raman Road, Bangalore, Karnataka 560012, India

## Abstract

Biological membrane is a complex self-assembly of lipids, sterols and proteins organized as a fluid bilayer of two closely stacked lipid leaflets. Differential molecular interactions among its diverse constituents give rise to heterogeneities in the membrane lateral organization. Under certain conditions, heterogeneities in the two leaflets can be spatially synchronised and exist as registered domains across the bilayer. Several contrasting theories behind mechanisms that induce registration of nanoscale domains have been suggested[1–3]. Following a recent study[4] showing the effect of position of lipid tail unsaturation on domain registration behavior, we decided to develop an analytical theory to elucidate the driving forces that create and maintain domain registry across leaflets. Towards this, we formulated a Hamiltonian for a stacked lattice system where site variables capture the lipid molecular properties such as the position of unsaturation and various other interactions that could drive phase separation and interleaflet coupling. We solve the Hamiltonian using Monte Carlo simulations and create a complete phase diagram that reports the presence or absence of registered domains as a function of various Hamiltonian parameters. We find that the interleaflet coupling should be described as a competing enthalpic contribution due to interaction of lipid tail termini, primarily due to saturated-saturated interactions, and an interleaflet entropic contribution from overlap of unsaturated tail termini. A higher position of unsaturation is seen to provide weaker interleaflet coupling. Thermodynamically stable nanodomains could also be observed for certain points in the parameter space in our bilayer model, which were further verified by carrying out extended Monte Carlo simulations. These persistent non-coalescing registered nanodomains close to the lower end of the accepted nanodomain size range also point towards a possible “nanoscale” emulsion description of lateral heterogeneities in biological membrane leaflets.

## I. INTRODUCTION

The biological cell membrane contains a host of lipids and other biomolecules such as proteins and glycans, which interact dynamically to facilitate biological processes like signal transduction and membrane protein oligomerization. Lipids in model ternary and quaternary membrane systems can segregate into liquid ordered (*L_o_*) and liquid disordered (*L_d_*) phases. Experimentally observed phase separated domains from *in vitro* studies range from micrometers to nanometers in their size[5–12]. Apart from studies on model in *vitro* systems, segregation of lipids into *L_o_* and *L_d_* phases has also been observed in *in vivo* membranes[13–16]. The functional importance of this phase separation is exemplified during signal transduction across cells, where the relay of signal from the outer to inner leaflet and vice-versa is highly dependent on the communication between the two leaflets. Experimental studies have shown the importance of lipids in synchronising the reception and transmission of messages across the bilayer[17–19]. T-cell and B-cell receptor mediated immune response is a classical example of systems where co-localization of outer and inner leaflet ordered domains, i.e ”registration” of ordered lipids, plays an important role in lipid mediated signal transduction[20–22]. Understanding the origin and molecular driving forces giving rise to ”registration” or ”anti-registration” of the ordered domains in the two leaflets can provide important insights into the related physiologically critical processes.

Localization features such as ”registration”, ”anti-registration”, and the other intermediate states are ultimately subjective and context dependent. The schematic in Fig.1 provides a qualitative understanding of what we mean when we use the terms ”registered”, ”partially registered”, ”unregistered”, ”partially anti-registered”, and ”anti-registered” in this article.

**FIG. 1:**
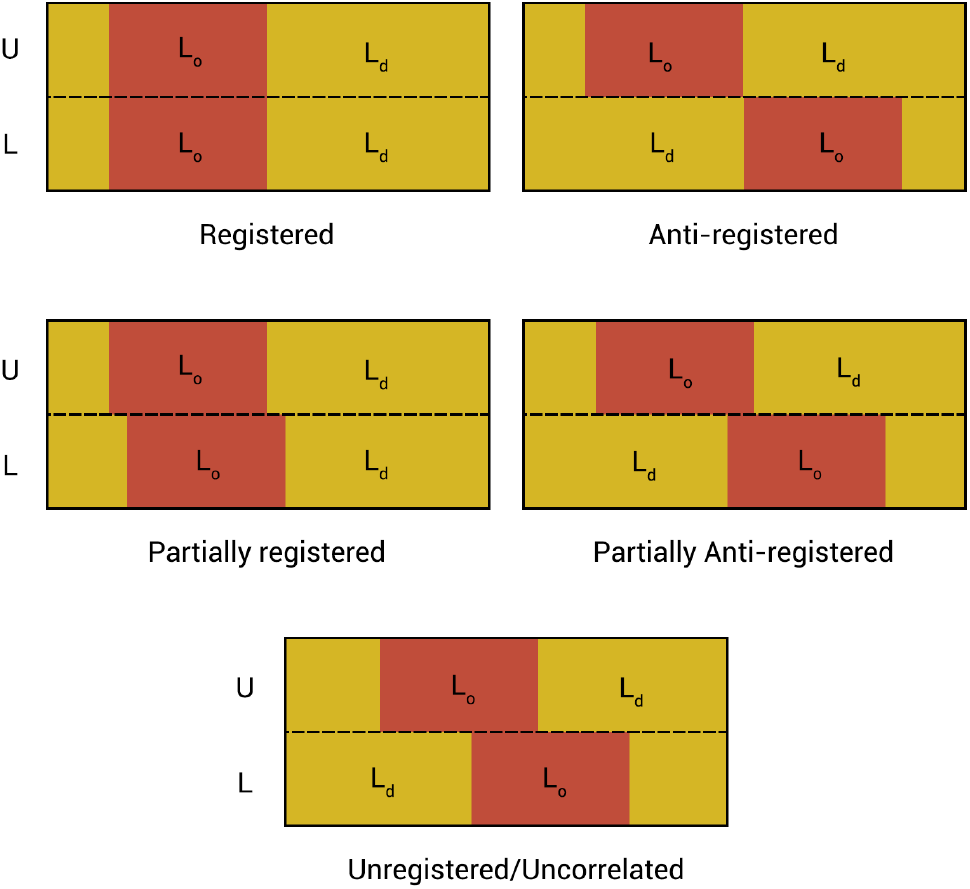
A schematic showing the qualitative meanings for the different states of registration discussed in this article. The membrane sections are represented as rectangles, with red regions corresponding to *L_o_* domains and yellow regions representing *L_d_* domains, with the labels U and L representing the upper and lower leaflet respectively.

Existing studies on the topic contain a variety of theories, some of them conflicting with each other, on the nature of the driving forces for domain registration. Exchange of cholesterol between ordered domains in the outer leaflet and the inner leaflet in asymmetric membranes has been hypothesised to form and maintain registered ordered domains in the inner leaflet in some theoretical models[23–25]. A mismatch energy, contributed when ordered and disordered domains overlap between leaflets, has also been studied as a possible driving force for domain registration[1]. This mismatch energy can be understood in terms of the energy penalty required when *L_o_* and *L_d_* domains interact between leaflets. In contrast, however, several studies claim membrane undulations and domain bending stiffness to cause domain registration without any direct interleaflet coupling[3, 26]. There is also work done suggesting domain boundary line tension as a driving force for registration and this model also does not require explicit coupling of the lipid bilayer leaflets[27, 28]. These studies argue that the difference in splay rigidities between the *L_o_* and *L_d_* domains in both leaflets causes preferential distribution of stiffer domains in regions of the membrane with lower curvature fluctuations. This, along with the energy gain from a decrease of line tension at the domain boundaries, is suggested as a possible explanation towards a driving force for registration of both large and small domains. There have been theoretical and computational studies on another candidate for a possible driving force, hydrophobic mismatch, arising due to a difference in tail lengths leading to differential membrane thickness between *L_o_* and *L_d_* domains[2, 16, 29–32]. Both domain formation kinetics and registration dynamics have been studied using hydrophobic mismatch as a driving force, and it has been theoretically shown to produce phase coexistence metastable states that capture both registered and antiregistered possibilities upon phase separation.

While there is agreement that there isn’t one single driving force for this phenomenon, the complexity of the problem comes from the number of hypotheses available that seem to explain domain registration or antiregistration. Zhang and Lin[4] made a new addition to this list of candidates by showing a significant effect of the position of unsaturation along the lipid tail on domain registration tendencies in otherwise identical systems. The authors carried out MARTINI[33] coarsegrained molecular dynamics (CGMD) simulations of two systems that were a mixture of DPPC, Cholesterol and an unsaturated lipid with different positions of unsaturation, which they called D23 and D34. Fig.2 shows the chemical structures of the lipids.

**FIG. 2:**
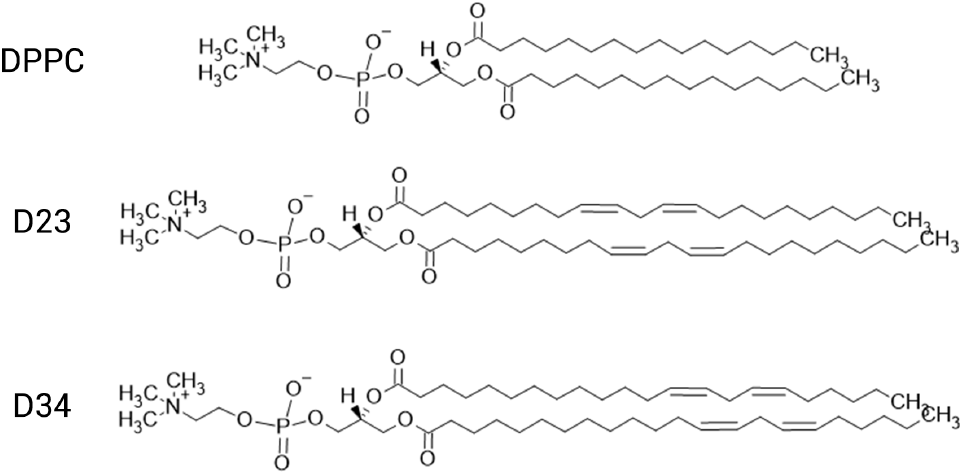
The chemical structures of the DPPC, D23 and D34 lipids. Note that the D23 and D34 lipids differ only in their position of unsaturation and do not occur naturally.

They observed that the system where the unsaturated lipid had lower position of unsaturation showed registration and the other system showed anti-registration. They also suggested that the interleaflet coupling was through an attractive interaction in the interleaflet region between the tail termini of the lipids.

We found this to be a very intriguing face of the problem, and decided to investigate the physical cause of the position of unsaturation affecting domain registration characteristics, which we believed could provide important insights into the overall registration mechanism. Besides testing the deductions made in the original work that solely credited the enthalpic interaction in the interleaflet region for driving registration, we also wanted to explore the role of the competing entropic factors in the interleaflet region. Our primary hypothesis is as follows: A higher position of unsaturation would lead to reduced configurational entropy of the lipid tails in the core of the membrane leading to a reduction in interleaflet interaction. On the other hand, a lower position of unsaturation forces the tail terminus to explore the interleaflet region (instead of bending towards the polar headgroups), which would lead to not only a better enthalpic but also an enhanced entropic contribution to the interleaflet coupling. The entropic aspect of interleaflet coupling is an important and often overlooked aspect that we bring to light in this work.

To test this, we formulate a Hamiltonian with tunable parameters that captures the hypothesised enthalpic and entropic contributions as a function of position of unsaturation along the length of the lipid tail for a lattice system representing the bilayer as a stack of two square lattices. We used this Hamiltonian and conducted a parameter study using Monte Carlo simulations for each point in parameter space for all the systems under investigation. Using these simulations, we try to capture the most dominant factor affecting the registration characteristics of the systems, as well as the effect of the change in position of unsaturation for a system described by our Hamiltonian. The schematic of our workflow is provided in sec.I of the Supporting Information (S.I) as Fig:S1.

We describe our lattice model of the membrane bilayer in the ”Model” section below and provide simulation details, specifications of the studied systems, as well as details about our analysis tools in the ”Methods” section. Following that, we report our observations and findings in the ”Results” section and we put all our results in the context of existing literature and provide a detailed perspective on the implications of our observations in the ”Discussion” section. We finally summarize our work in the ”Conclusions” section with some perspectives on position of unsaturation in lipids as a driving force for domain registration.

## II. MODEL

Following our hypothesis, we decided to construct a Hamiltonian for a system with two coupled membrane leaflets that captured entropic behaviour of the lipid tails as well as the positions of unsaturations in the lipid tails. Each membrane leaflet is modeled as a square lattice, populated by site variables representing different lipids.

The site variables for each species of lipid are Deuterium Order Parameter values, commonly referred to as *S_CD_*, averaged over the length of the lipid tail, and given by

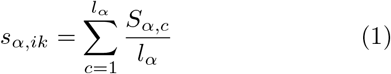

where i is a site on a leaflet, k is the leaflet index, which takes values 0 and 1 representing the lower and upper leaflet respectively. *l_α_* represents the length of the lipid tail of species *α* in terms of number of carbon atoms, and *S_α,c_* represents the *S_CD_* of the *c^th^* carbon atom in the tail of a lipid of species *α*, given by

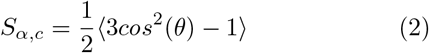

Here, the angle *θ* is the angle between the C-D bond and a reference axis, and the angular brackets represent an ensemble and time average. We obtain the site variables for different lipids through averages taken from all-atom simulations described in section III, and they capture the tail disorder for the respective species of lipids along with the effects of the lipids’ environment. We use these site variables to define Ising-like enthalpic interactions separately for the lateral interactions within a leaflet, as well as interleaflet interactions which are meant to represent the enthalpic interactions arising from the slight overlap of the tail termini in the interleaflet region.

Let us take *ϵ_αα′_* to represent the interaction energy within the leaflet between lipids of species *α* and *α′* occupying orthogonally adjacent sites. Then, we have

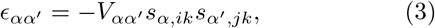

where i and j are adjacent lattice sites, and the parameter *V_αα′_* is a species dependent interaction strength constant. From this, we can write the lateral enthalpic interaction energy contribution over both leaflets, by summing over all pairs within a leaflet for both leaflets, as

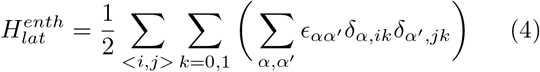

where the *δ*s are kronecker deltas such that *δ_α,ik_* is 1 if site i in leaflet k is occupied by a lipid of species *α*.

This choice of lateral enthalpic interaction is made due to the ease of achieving phase separation. While phase separation isn’t our main phenomenon of interest, it is important for phase separation to occur to some degree for us to look at domain registration/anti-registration behaviour.

Using similar notation as in the equations above, we also define an interleaflet enthalpic interaction given by

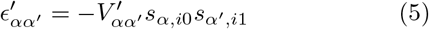

where the interaction occurs between the same site i on the leaflets 0 and 1, and 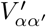 is a species dependent interaction strength parameter. Therefore, the interleaflet enthalpic energy contribution for our model is given by

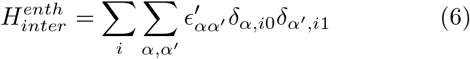

We now move on to the entropic contributions that we intended to capture in our model.

We include an entropy of mixing within each leaflet in a Flory-Huggins like form. For a site i, the contribution to entropy of mixing is given by

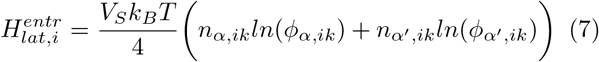

where *n_α,ik_* is the count of lipids of species *α* in the local neighbourhood of site i in leaflet k, with the neighbourhood shown in Fig.3(a), and *ϕ_α,ik_* is the fraction of lipids of species *α* in the same local region. *V_s_* is a strength constant that can be used to tune the magnitude of this contribution. Summing over all sites and both leaflets, we get the total lateral entropy of mixing to be

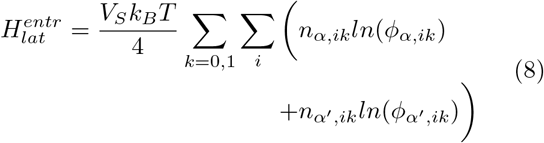

**FIG. 3:**
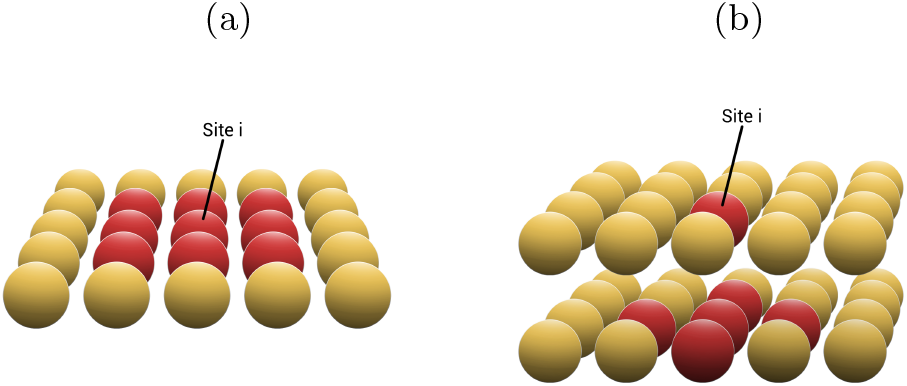
Cartoon representation of a local region of the model, where the red sites are the locality over which the parameters *n_α_* and *ϕ_α_* are calculated for a lattice site i. (a) represents the sites for the entropic term of *H_lateral_*, and (b) represents the sites for the entropic of *H_interleaflet_*

The final and most crucial part of our Hamiltonian is the interleaflet entropic contribution. This is meant to model the entropy of mixing due to the tail termini slightly overlapping in the interleaflet region, which we believe would be affected by the differences in position of unsaturation. To write such an energy contribtuion, we use a similar form as above, but scaled by a factor depending on the position of unsaturation. So, for a site i in leaflet k, the contribution is given by

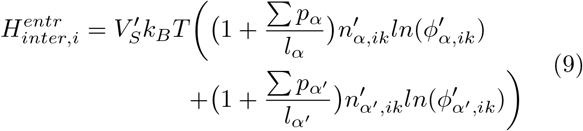

where *p_α_* is the position of an unsaturation along a lipid tail of species *α*, where the carbon atoms are numbered starting from 0 at the ester bond, with the position of unsaturation given by the number of the carbon atom closer to the head group out of the two carbons comprising the unsaturated bond. Also, *p_α_* is summed over the different unsaturations that might be present in a tail. In this case, 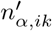 and 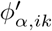 have the same meanings as before, but calculated over a different local neighbourhood, which is shown in Fig.3(b). 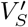 is a strength constant that can be used to tune the magnitude of the contribution. Like before, summing over all sites and both leaflets gives us the total interleaflet entropic contribution as

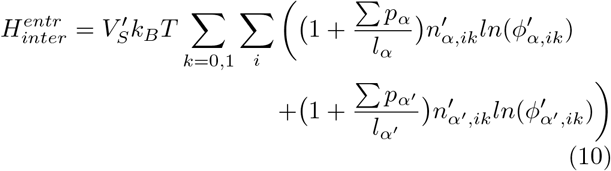

Now, if we put these terms together, we can write the Hamiltonian to be made up of a lateral and an interleaflet part given by

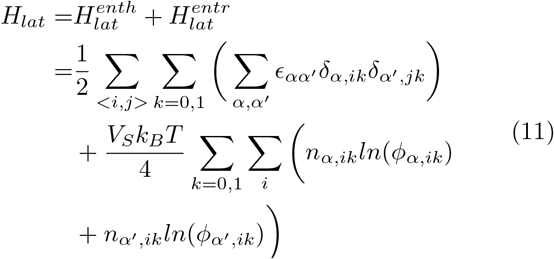

and

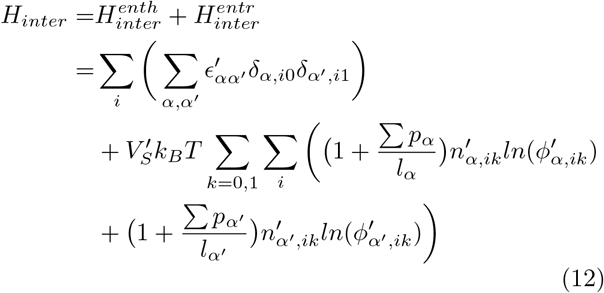

with the total Hamiltonian being given by

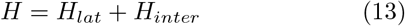

For a system of two lipids A and B, this gives us a Hamiltonian with 9 parameters that are tunable during simulations, shown together for convenience in Table.I below.

**TABLE I:** Description of all the tunable independent parameters of the Hamiltonian for a 2-component system with lipid species A and B.

| Parameter | Description |
| --- | --- |
| T | Temperature |
| $V_S$ | Lateral entropic term strength constant |
| $V'_S$ | Interleaflet entropic term strength constant |
| $V_{AA}$ | Lateral enthalpic strength constant for A-A interaction |
| $V_{BB}$ | Lateral enthalpic strength constant for B-B interaction |
| $V_{AB}$ | Lateral enthalpic strength constant for A-B interaction |
| $V'_{AA}$ | Interleaflet enthalpic strength constant for A-A interaction |
| $V'_{BB}$ | Interleaflet enthalpic strength constant for B-B interaction |
| $V'_{AB}$ | Interleaflet enthalpic strength constant for A-B interaction |

We use the Hamiltonian as defined above to calculate the energy for our systems in the Monte-Carlo simulations used in this study, which we describe in the following section.

## III. METHODS

We begin this section by describing how we obtained the site variables and describing the systems we studied, followed by how they were used to generate data using Monte-Carlo simulations. We also provide details of the analysis tools used to obtain quantitative results on phase separation and domain registration tendencies of the systems from the generated data.

### A. All-atom simulations of artificial lipids

We train the values of the site variable (see eqn.1) for each of the lipids used in the Monte-Carlo simulations from all atom trajectories of DPPC-D23-Chol and DPPC-D34-Chol systems. The study by Zhang and Lin[4] used CGMD simulations where D23 and D34 are artificial lipids. To capture the lipid molecular properties faithfully, we reconstructed an atomic resolution system and carried out AAMD simulations. In our work, we have used a prescription that allows us to build all-atom structures for artificial lipids that have a similar chemical makeup to lipids with well-known forcefield parameters. We provide a schematic of the workflow in Fig.4 and we describe the steps in the following text. We prepared the all-atom descriptions of the artificial lipids, namely D23 and D34 lipids using the Ligand Reader and Modeler Module in CHARMM-GUI[34, 35]. Since we aren’t using partial charges from Quantum-Chemical calculations, we need to check the assigned partial charges with a closely matching parameterized lipid molecule which, in this case, were the partial charges on the sp^2^ and sp^3^ hybridised lipid tail carbons in the lipid DUPC. We then build the multi-component lipid bilayer using the MemGen webserver[36] based on the percentage composition of the required bilayer and the number of lipids per leaflet in our system. We have organized a tutorial with examples for the workflow described above and made it publicly available at https://github.com/codesrivastavalab/Membrane-Nanodomain-Registration/tree/main/Artificial%20Lipid. Also, please note that common modelling errors and issues during the generation of the all-atom bilayer systems might require some troubleshooting, which we discuss in sec.2 of the S.I.

**FIG. 4:**
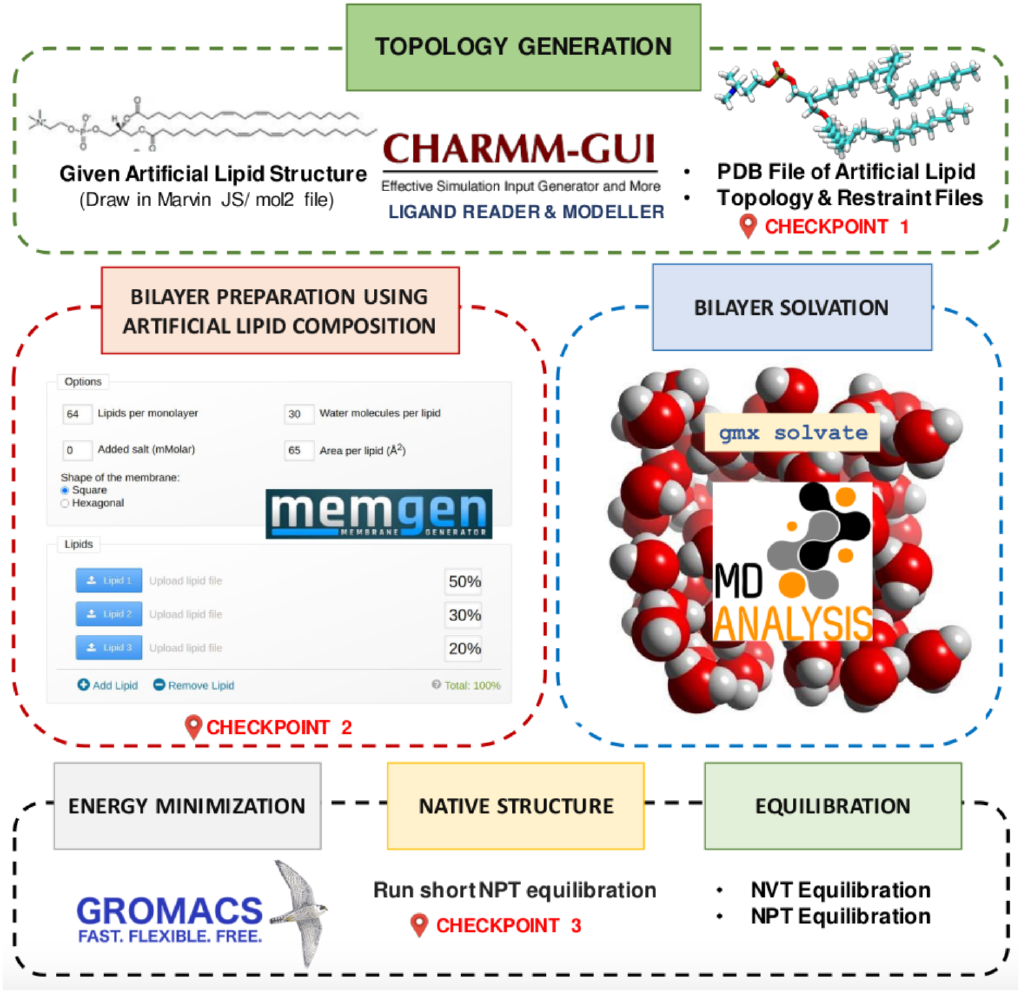
Schematic of the workflow for generating artificial lipid force field parameters for lipids with different position of unsaturation and generation of the bilayer topology. The schematic uses logos of CHARMM-GUI[34, 35], MemGen[36], MDAnalysis[37, 38], and GROMACS[39, 40] softwares along with some molecule representations generated using VMD[41].

For our systems, the average *S_CD_* was calculated for DPPC, D23 and D34 lipids from the all-atom system of 144 DPPC, 80 D23/D34, and 56 Chol molecules in each leaflet, which is close to the ratio used by Zhang and Lin[4]. We used the CHARMM36 force field and first minimized the energy of the system using steepest descent minimization. Then, we carried out two rounds of 125 ps NVT equilibration at 303.15 K and four rounds of 250 ps NPT equilibration. We ran production runs using GROMACS[39, 40] for a period of 1*μs* with the PME[42, 43] method for electrostatics, Nose-Hoover[44] temperature coupling and Parinello-Rahman[45] pressure coupling. The LINCS[46] algorithm was used to restrain hydrogen bonds. The tail order parameters of D23 and D34 lipids were calculated using the gmx order program in GROMACS. The *S_CD_* values obtained are averaged over all the D23/D34 lipids in the D23/D34, DPPC, Chol ternary mixture and across 7.5 ns (*S_CD_* remain the same with increased averaging time). *S_CD_* of *n^th^* carbon is calculated using the position of *n* – 1 and *n* + 1 carbons.

Fig.5 shows the *S_CD_* values for D23 and D34 lipids. The decrease in the *S_CD_* values occurs at the position of double bonds in D23 and D34 lipids. The dip in value of *S_CD_* for the C atoms near the end of the alkyl tails of D34 lipids indicates the lower position of the double bonds as seen from the molecular structure of the lipids. On an average, D34 lipids show more ordering of the alkyl tails in comparison to D23 lipids. The final values of the site variables calculated by averaging the *S_CD_* over all tail carbon atoms for the DPPC-D23 systems were DPPC-0.3074681, D23-0.1692048, and for the DPPC-D34 systems, they were DPPC-0.3565843, D34-0.2400412. As we can see, the average *S_CD_* is higher for the unsaturated lipid with a lower position of unsaturation. Please note that the purpose of the AAMD simulations is only to extract out a more realistic site variable for our Hamiltonian.

**FIG. 5:**
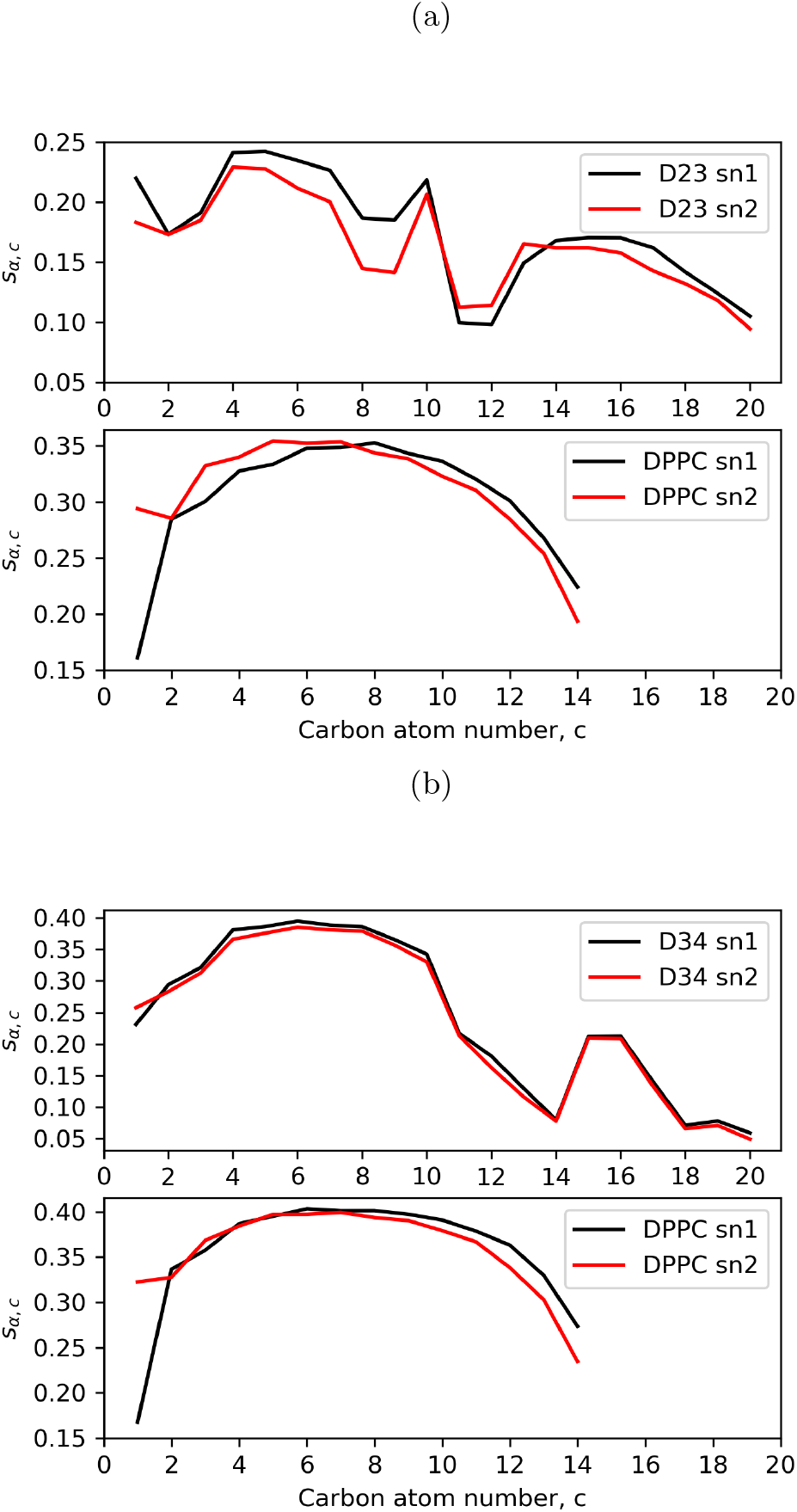
a) shows the *S_CD_* for the tail carbon atoms of the D23 and DPPC lipids as calculated by averaging over all D23 lipids of the D23-DPPC system. b) Shows the same for the D34-DPPC system.

### B. Setup for Monte Carlo simulations

We studied 6 systems, each being a stack of 2 100 × 100 square lattices. The upper and lower leaflet have symmetric populations in each system, with no lattice points left empty. So, each leaflet contains 10,000 lipids of different species depending on the system. The details of the 6 systems are shown in table.II.

**TABLE II:** Composition of the bilayers of the 6 systems with symmetric leaflets.

| Components | System Composition |
| --- | --- |
| DPPC, D23 | $2 \times (5000 \text{ DPPC}, 5000 \text{ D23})$ |
| DPPC, D23 | $2 \times (2500 \text{ DPPC}, 7500 \text{ D23})$ |
| DPPC, D23 | $2 \times (7500 \text{ DPPC}, 2500 \text{ D23})$ |
| DPPC, D34 | $2 \times (5000 \text{ DPPC}, 5000 \text{ D34})$ |
| DPPC, D34 | $2 \times (2500 \text{ DPPC}, 7500 \text{ D34})$ |
| DPPC, D34 | $2 \times (7500 \text{ DPPC}, 2500 \text{ D34})$ |

Each system underwent a Monte-Carlo simulation with traditional Metropolis acceptance[47] for each combination of parameters in the following parameter space in reduced units

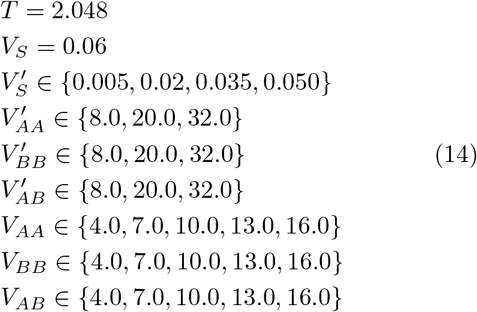

with the restriction that *V_AB_* ≤ *V_BB_* ≤ *V_AA_*, since points outside this would not show phase separation. This gives us 3780 points in parameter space for each of the 6 systems. The temperature in reduced units corresponds to the temperature of the all-atom simulations from which the site variables were calculated. Also, we chose the range of the in-plane enthalpic strength constants such that the interaction energy would lie in the neighbourhood of the physiological DPPC-DPPC interaction energy estimated by P. Almeida in 2009[48]. Further references to any point in parameter space will be written as a chain of indices of parameters in the order shown in eqn.14 above. For example, point 0-0-0-2-1-0-4-4-0 refers to the parameters *T* = 2.048, *V_S_* = 0.06, 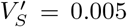, 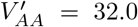, 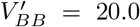, 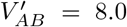, *V_AA_* = 16.0, *V_BB_* = 16.0, *V_AB_* = 4.0.

The systems for each point in parameter space were initialized by randomly choosing from their two components at each lattice site, with a check to ensure we didn’t exceed the specified population for the chosen species. This caused the initial configurations to have regions of one or the other species, but this was lost very quickly in a few MC moves during the simulations, thereby not affecting the converged state of each system for any point in the parameter space. Each initialized system corresponding to a point in the parameter space underwent Monte-Carlo simulations for 10^7^ moves, each move consisting of an exchange attempt of site variables between two randomly chosen sites, first for the upper leaflet followed by an attempt for the lower leaflet separately. The system configurations were output every 5000 moves, giving us 2000 configurations for each point in parameter space for each system. The last 200 configurations were used for analysis. The Monte-Carlo data for each point in parameter space for each of the 6 systems was further analysed using tools as described ahead.

### C. Analysis tools

Owing to the large number of parameter space points for each of the 6 systems, it is impractical to visually classify the phase separaton and domain registration of each point in parameter space, so we wrote simple C routines to do the classification for us without visual aid, whose functioning is described below. The programs used for the analysis described here as well as that for the MC simulations described in the previous section have been made publicly available at https://github.com/codesrivastavalab/Membrane-Nanodomain-Registration/tree/main/Lipid%20Domain%20Registration.

#### 1. Quantification of phase separation using domain size distributions

Nanoscale molecular-level heterogeneity and structures are non-trivial to quantify [49–52] .We used a Depth-First Search (DFS)[53] based algorithm to determine domain size distributions for each leaflet for a given configuration to categorize them as phase-separated(PS), not phase-separated(NPS) or partially phase separated(PPS) based on a consistent set of cutoffs. If > 80% of lipids of each species were in domains of size > 1600 lipids, they were categorized as PS, if > 80% of lipids of either species were in domains of size < 400 lipids, they were considered NPS. Any configurations not in either category were considered to be PPS. A point in parameter space was assigned one of the three states if a majority of configurations from the last 10% of output configurations in it’s Monte-Carlo simulation were of that state. Although the cut-offs are somewhat arbitrary, since they are applied consistently to all of the systems, the results still show reliable qualitative trends with the variation of the tunable parameters.

#### 2. Domain registration quantification using KL Divergence

For domain registration, we used Kullback-Leibler(KL) divergence[54] of the two leaflets as a measure of registration. KL Divergence (denoted by *D_KL_*) was the tool of choice for us because of its simplicity as well as its ability to clearly and quantitatively separate systems based on fixed cut-offs. *D_KL_* for a pair of probability distributions P(x) and Q(x) on a random variable x on the same probability space is given by

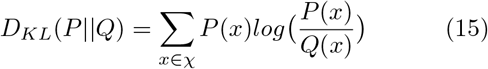

where P(x) and Q(x) are constructed from a 2D leaflet configuration of the upper and lower leaflet respectively from the Monte-Carlo trajectories of a system with lipids A and B as follows

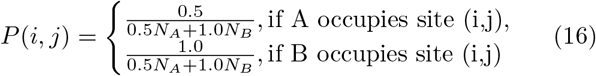

where *N_A_* and *N_B_* are the populations of lipids of species A and B, respectively.

Q(i,j) is identically defined, but in the lower leaflet. The choice of weights 0.5 and 1 is arbitrary and it doesn’t affect our categorization as we rescale the *D_KL_* values of each system to lie between 0 and 1 before continuing with classification of the data. Configurations were categorized as registered (R), partially registered (PR), unregistered (UR), partially anti-registered (PAR), and anti-registered (AR) based on cut-off values of *D_KL_* at 0.1, 0.3, 0.7, and 0.9, which correspond to 10%, 30%, 70%, and 90% of the lipids of lowest population in mismatched configurations between leaflets. Once again, the category of the majority of the analysed configurations from the Monte Carlo simulations of a point in parameter space was assigned to that point.

## IV. RESULTS

In this section, we describe the results of the simulations and analysis that we discussed above. We first look at the results of the MC simulations. We have reported the convergence of the simulations in section III of the SI, where we have shown the energy profiles of randomly selected points from across the parameter space to show sufficient convergence of the MC simulations. The Hamiltonian that we wrote can successfully capture phase separation in leaflets, and can capture different extents of domain registration and anti-registration behaviour. Fig.6 shows some select configurations generated at various points in parameter space in our systems to illustrate this.

**FIG. 6:**
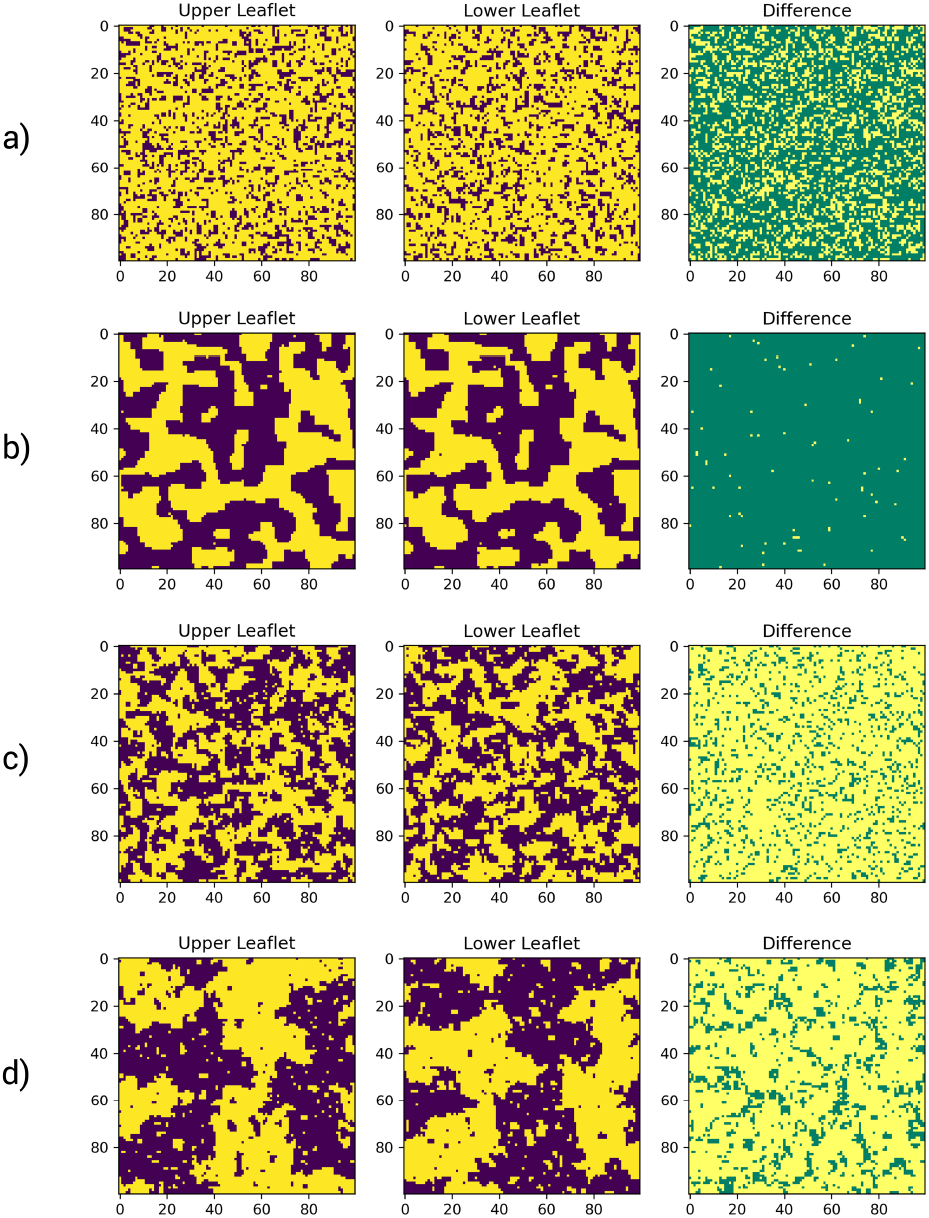
In the shown leaflet configurations, yellow represents DX sites and purple represents DPPC sites. In the difference plot, yellow represents a mismatch and green represents a match between leaflets at that site. a) shows the upper and lower leaflet configurations with their difference for parameters 0-0-0-1-1-1-2-2-0 for the 1:3 DPPC-D34 system, which is not phase separated (NPS, UR), b) shows the same for parameters 0-0-0-2-0-0-4-4-0 for the 1:1 DPPC-D34 system which is phase separated and registered (PS, R), c) for parameters 0-0-0-2-0-2-4-0-0 for the 1:1 DPPC-D34 system which is partially phase separated and partially anti-registered (PPS, PAR), and d) for parameters 0-0-1-2-2-2-4-4-0 for the 1:1 DPPC-D34 system which is phase separated and anti-registered (PS, AR).

After using our analysis programs to classify the points in parameter space for all 6 systems for both registration and phase separation using the last 10% of the configurations from the Monte carlo simulations of each point in parameter space to conduct the analysis and categorization, we now look at the trends shown in phase separation and domain registration with a variation in the different parameters.

### A. The lateral enthalpic parameters have the most significant effect on phase separation

We plotted histograms of the populations of parameter space points in various states of phase separation and domain registration. As can be seen from Fig.7 for the 1:1 DPPC-D34 system, we find that the relative proportion of PS points increases and NPS points decreases as either *V_AA_* or *V_BB_* increase. Conversely, upon increasing *V_AB_*, we find that the relative proportion of PS points decreases and NPS points increase. This effect on phase separation, being purely enthalpic in its origin, is reflected in all the 6 systems studied, which is an unsurprising but reassuring observation to establish the fidelity of our modeling work. It must be noted that the number of points in parameter space are not equal for each value of the lateral enthalpic parameters due to the restriction we have demanded, i.e; *V_AB_* ≤ *V_BB_* ≤ *V_AA_*. What is important, instead, in these histograms with the varying lateral enthalpic strength constants, is the relative proportions of the different states at each value of the parameter. The 1:1 DPPC-D23 system shows the same trends, but with an overall lower propensity towards PS, and higher tendency towards NPS states. The plot has been included in sec. IV of the S.I as Fig:S3. One interesting result is that phase separation is highly suppressed in our 1:3 and 3:1 systems, as illustrated by Fig.8, which shows the same plot for the 1:3 DPPC-D34 system. This is in strong concurrence with results shown in in-silico study by Perlmutter and Sachs[55] where they observed that any deviation from equal amounts of *L_o_* and *L_d_* regions led to a suppression of phase separation, of which our result is an extreme case. This shows that a significant skew from a 1:1 proportion doesn’t favor phase separation, and is in accordance with previous experimental work on vesicular systems with biological lipids as well as results from theoretical models[9, 56, 57]. The remaining three systems show similar trends, and their plots have also been included in sec. IV of the S.I as Fig:S4, Fig:S5, and Fig:S6. From all six plots, one can extract some more general information, i.e, the DPPC-D34 systems have a slightly higher propensity to phase separate than the DPPC-D23 systems. Also, the 1:3 systems show a slightly higher population of PS and PPS points than their 3:1 counterparts, suggesting that the systems with higher populations of unsaturated lipids phase separate slightly better than those with higher populations of saturated lipids in each leaflet, which is also captured in phase diagrams in previous experimental work[56].

**FIG. 7:**
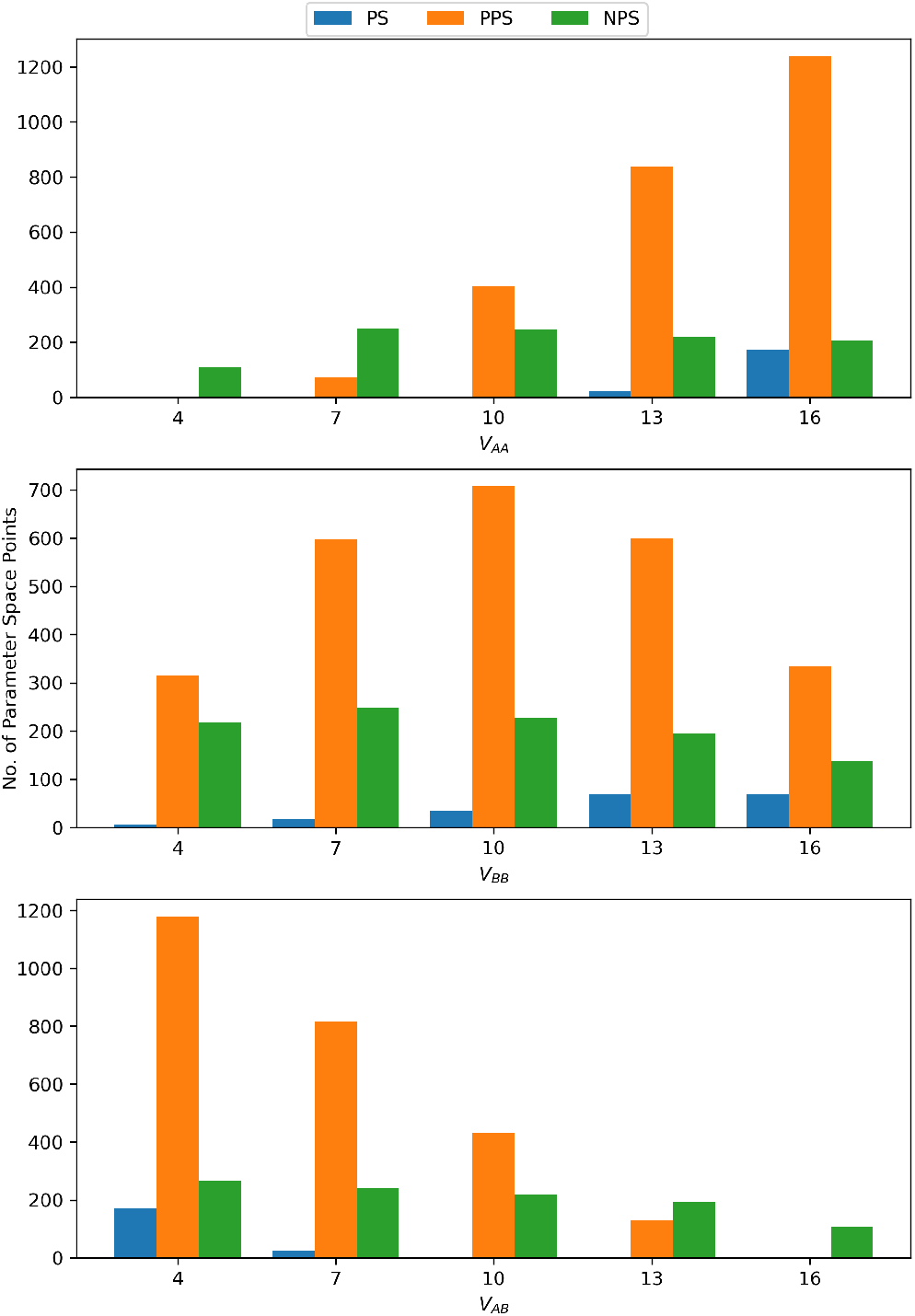
Histograms showing the populations of parameter space points in PS, PPS, and NPS states at different values of the lateral enthalpic interaction strength constants *V_AA_, V_BB_*, and *V_AB_* for the 1:1 DPPC-D34 system.

**FIG. 8:**
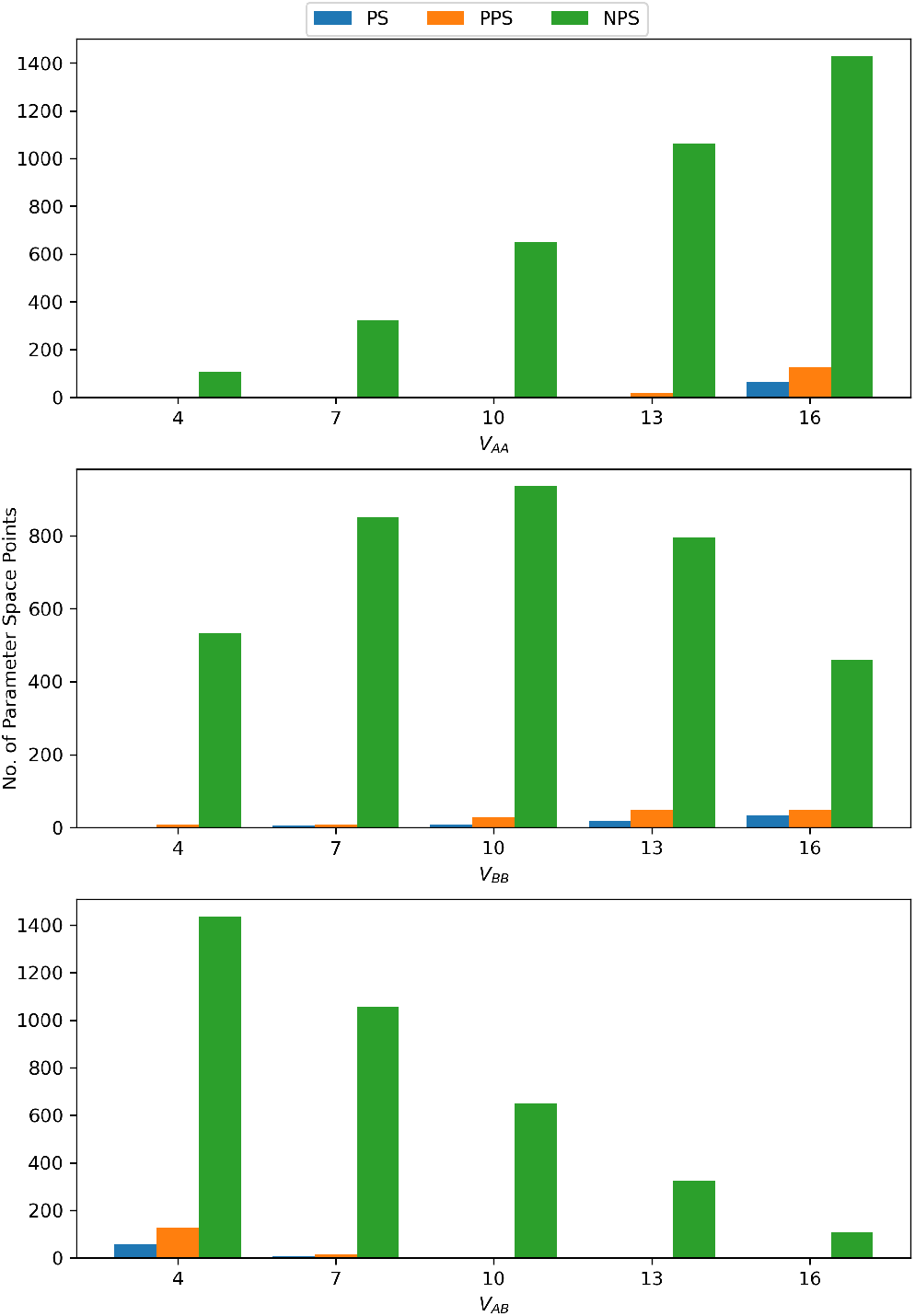
Histograms showing the populations of parameter space points in PS, PPS, and NPS states at different values of the lateral enthalpic interaction strength constants *V_AA_ ,V_BB_*, and *V_AB_* for the 1:3 DPPC-D34 system.

### B. Interleaflet enthalpic parameters have subtle effects on phase separation

By plotting similar histograms to observe the variation of populations with the variations in the interleaflet enthalpic interaction parameters 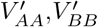, and 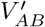, we find that the effects exist, but are very subtle. As can be seen from Fig.9, which shows the plots for the 1:1 DPPC-D34 system, 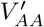 and 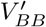 both show a slight decrease in PS points with their respective increase, and a slight increase in NPS points with their increase. 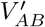, on the other hand, shows the exact opposite trend. Again, this is observed in both DPPC-D23 and DPPC-D34 systems, and the other plots are included in sec.V of the S.I as Fig:S7, Fig:S8, Fig:S9, Fig:S10, and Fig:S11. While the effect is quite subtle, the trends still clearly exist. It is known that the exact effect of the interleaflet enthalpic interactions on phase separation depend acutely on the lengths of the acyl chains of lipids involved[58]. We believe that the current result could be interpreted as an outcome of the composition of our leaflets containing the long acyl chain artificial lipids and relatively shorter DPPC tails, leading to a mismatch in lengths when self interaction between the leaflets is higher, thereby causing a suppression of phase separation. This mismatch has been hypothesised as a mode of domain anti-registration in various theoretical and in-silico studies[2, 16, 29–32]. In our case, it is an interesting result due to the fact that the length of the acyl chains itself is not explicitly represented in our model (*l_α_* only appears when averaging the *S_CD_*), and it could mean that the average *S_CD_* is somehow sufficient to capture the difference in the conformational behaviour of the lipids due to their different chain lengths.

**FIG. 9:**
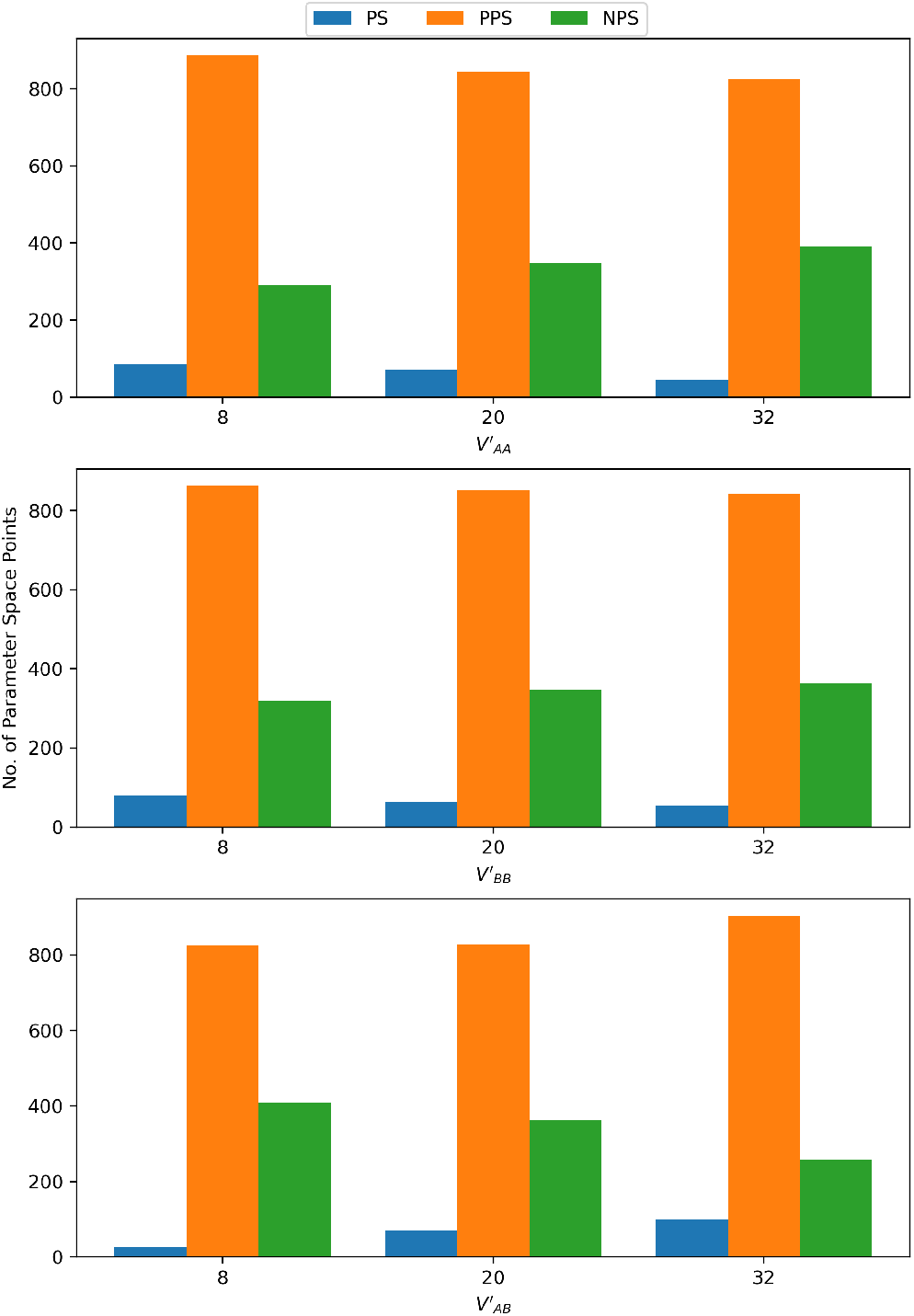
Histograms showing the populations of parameter space points in PS, PPS, and NPS states at different values of the interleaflet enthalpic interaction strength constants 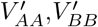, and 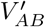 for the 1:1 DPPC-D34 system.

### C. Interleaflet entropic contributions suppress phase separation

The final parameter to look at for effects on phase separation is the interleaflet entropic strength constant. Again, we make a similar histogram as before for all six systems, of which the one for the 1:1 DPPC-D34 system is shown in Fig.10. As can be observed, an increase in the magnitude of the interleaflet entropic contribution causes a decrease in PS and PPS points, accompanied by a significant increase in NPS points. While there haven’t been any targeted studies on this aspect of interleaflet interactions, it could be a plausible explanation for the results obtained in recent experimental studies that find suppression of phase separation in leaflets capable of forming *L_o_* domains depending on the composition of the leaflets[]. Again, the other systems also show similar trends, and their plots are included in sec. VI of the S.I as Fig:S12, Fig:S13, Fig:S14, Fig:S15, and Fig:S16 to avoid crowding of plots here.

**FIG. 10:**
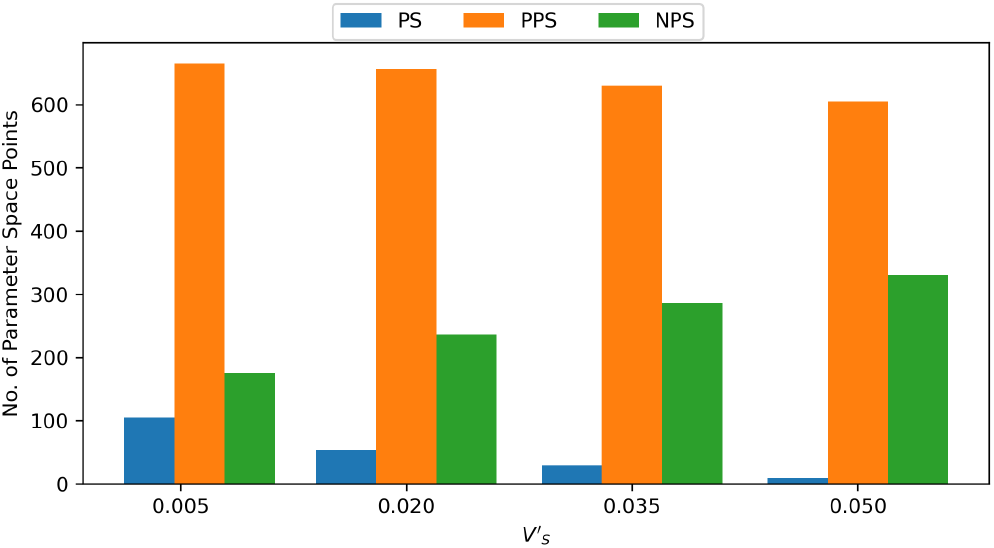
Histograms showing the populations of parameter space points in PS, PPS, and NPS states at different values of the interleaflet entropic strength contant 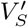 for the 1:1 DPPC-D34 system.

We will now proceed to look at the plots for domain registration and their variation with the parameters of the Hamiltonian in our systems.

### D. Lateral enthalpic parameters do not affect domain registration/anti-registration

By plotting histograms of the populations of R, PR, UR, PAR, and AR points against the different values of the lateral enthalpic parameters, we found that there was no significant change in the relative proportions of the R and AR points, as shown from the plot for the 1:1 DPPC-D34 system in Fig.11. Once again, we emphasize that the relative populations of the different states are more important in these plots due to the total number of points being different for each parameter value, given the constraint, i.e; *V_AB_* ≤ *V_BB_* ≤ *V_AA_* that is demanded of the lateral enthalpic interaction parameters. It must be noted that there is a general increase in both R and AR populations and a decrease in UR populations with higher values of *V_AA_* and *V_BB_*, and lower values of *V_AB_*. We believe that this is simply because these parts of the parameter space correspond to regions of high propensity for phase separation as seen earlier, and therefore, show a higher tendency to fall into either registered or anti-registered categories. However, the lateral enthalpic strength parameters do not favour R or AR in a significant manner. The plots for the remaining five systems are included in sec.VII of the S.I as Fig:S17, Fig:S18, Fig:S19, Fig:S20, and Fig:S21. Due to the very subtle changes in domain registration behaviour, we believe that the lateral enthalpic interactions of lipids cannot provide any observable effects towards domain registration in realistic or real membrane systems.

**FIG. 11:**
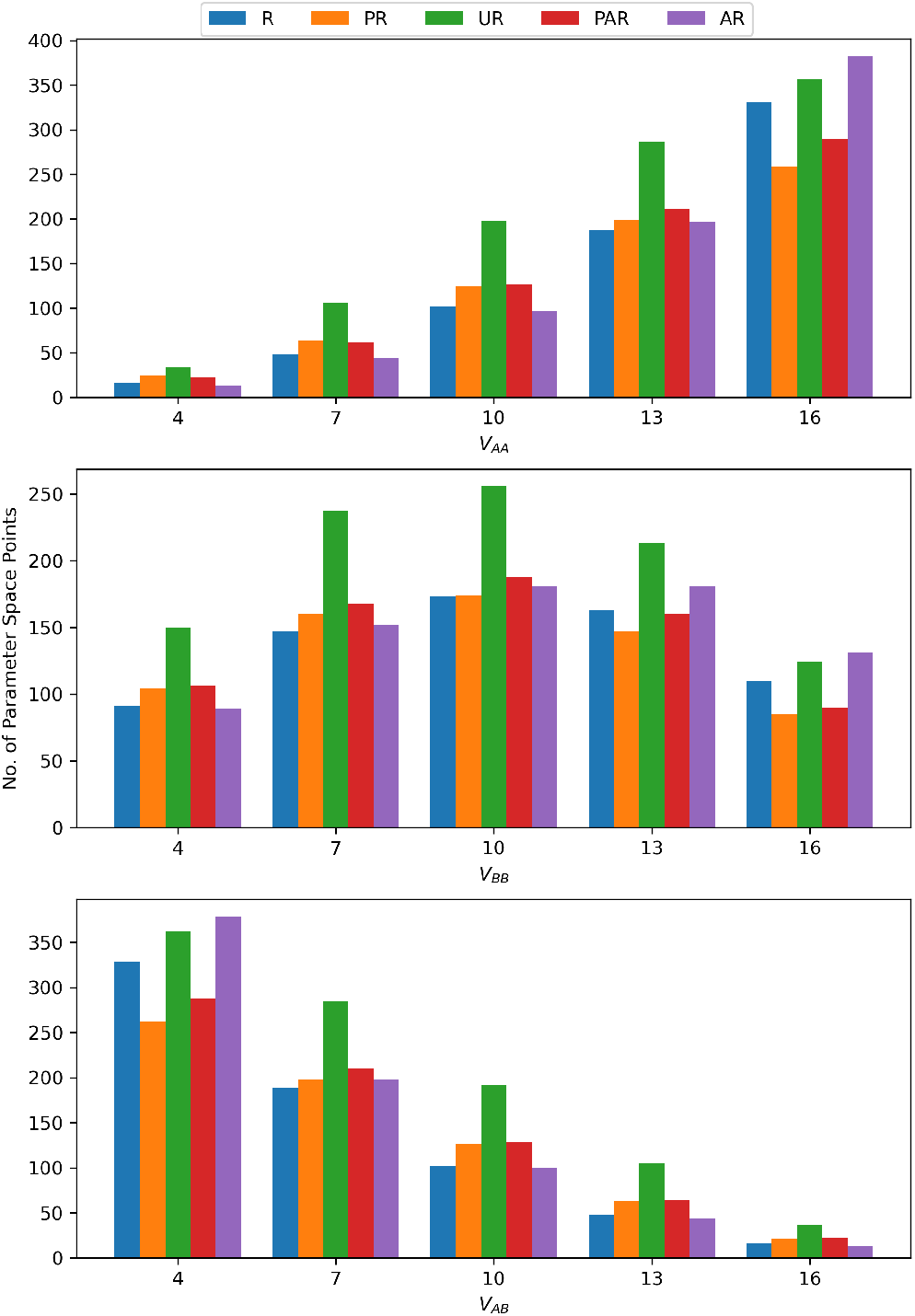
Histograms showing the populations of parameter space points in R, PR, UR, PAR, and AR states at different values of the lateral enthalpic interaction strength constants *V_AA_, V_BB_*, and *V_AB_* for the 1:1 DPPC-D34 system.

### E. Domain registration is promoted by interleaflet enthalpic interaction of saturated lipids

We made similar plots for the interleaflet enthalpic interaction strength parameters 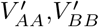, and 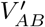. Fig.12 shows the plot for the 1:1 DPPC-D34 system, which shows a very clear trend of increasing R and decreasing AR points with an increase in 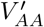 or 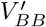. However, this trend is far more drastic for 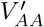 as compared to 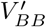, suggesting that the primary driving force for domain registration is the interleaflet enthalpic interaction of the saturated lipids in the system. Interestingly, looking at the plot for the 1:1 DPPC-D23 system in Fig.13, we see that there are far more PAR points than PR points, with overall low numbers of R and PR points. And while this plot shows the same trends as in Fig.12, the overall PAR and AR populations are significantly higher, which supports the observations from the work by Zhang and Lin[4]. While previous studies have found that a general interleaflet interaction[59] is sufficient to show domain registration, we believe our model provides a more refined look at the mechanism behind this phenomenon, which can be tested experimentally in vesicular systems. This is also a plausible explanation or contributing factor to the phenomenon of stabilisation of ordered domains in an opposing leaflet due to ordered domains in one which has been theorised[23, 24] as well as observed in experimental studies for specific lipid types and compositions[60–62]. We can also see the general properties that the DPPC-D34 systems have a larger fraction of points in the R or AR categories, whereas the DPPC-D23 systems have most of their points in the PR, UR and PAR categories as can be observed upon comparing Fig.12 and Fig.13. This would suggest that the energy landscape of the DPPC-D34 system in parameter space has a sharper transition from favouring registration to favouring anti-registration, whereas the DPPC-D23 system has a smoother energy landscape in parameter space. These general trends are also replicated in the plots of all 6 systems, which are shown in the S.I as Fig:S22, Fig:S23, Fig:S24, and Fig:S25.

**FIG. 12:**
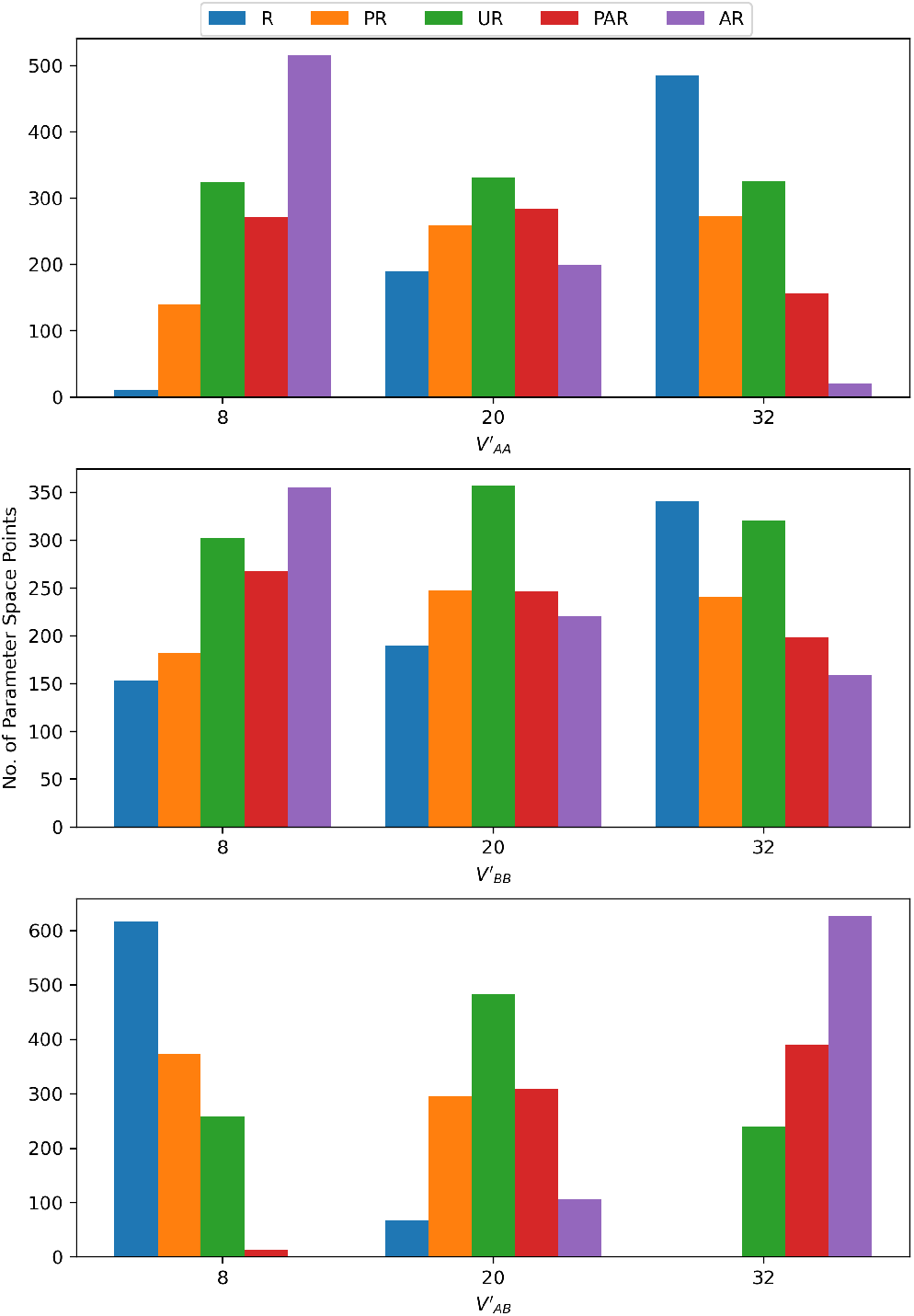
Histograms showing the populations of parameter space points in R, PR, UR, PAR, and AR states at different values of the interleaflet enthalpic interaction strength constants 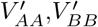, and 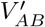 for the 1:1 DPPC-D34 system.

**FIG. 13:**
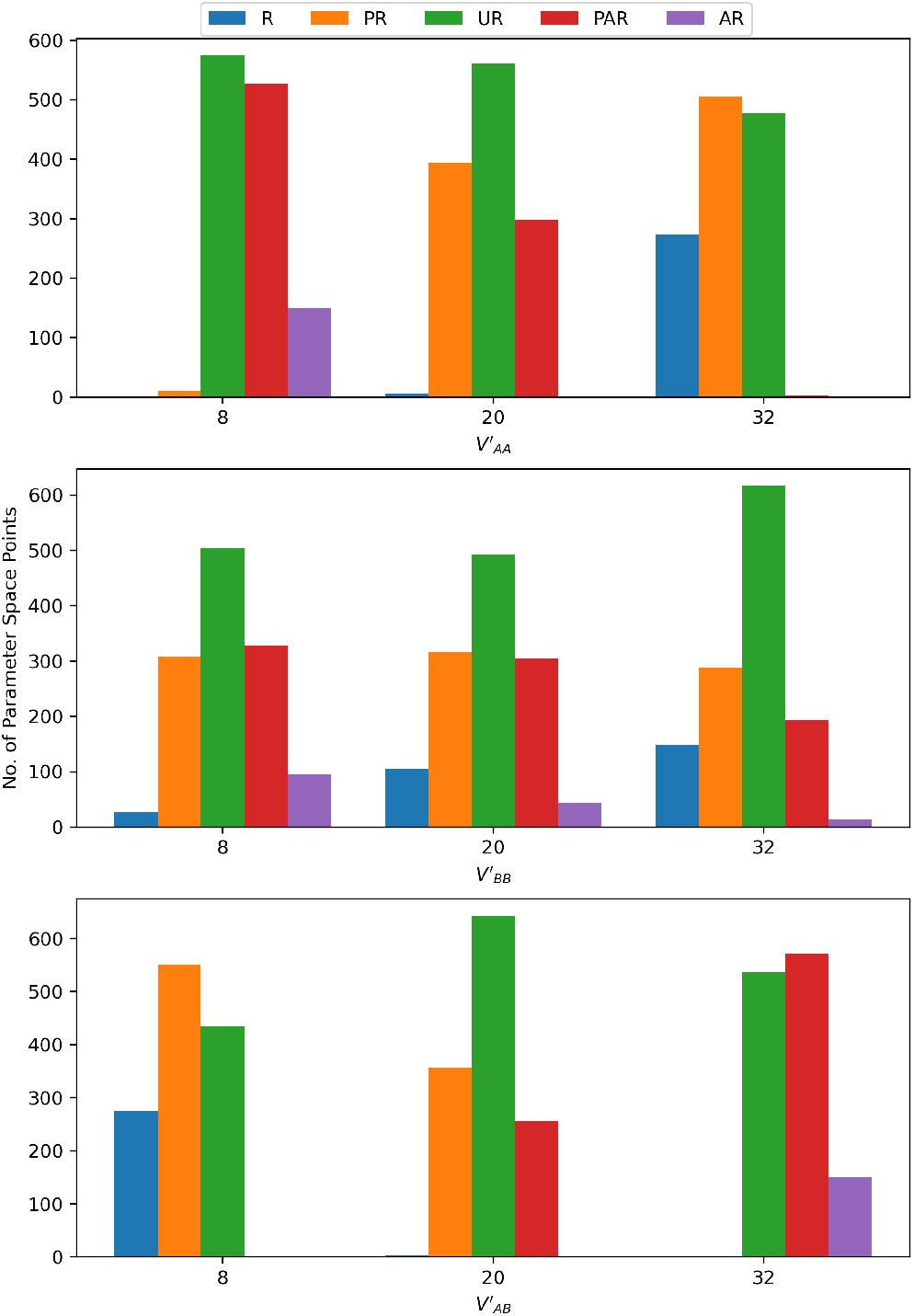
Histograms showing the populations of parameter space points in R, PR, UR, PAR, and AR states at different values of the interleaflet enthalpic interaction strength constants 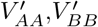, and 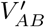 for the 1:1 DPPC-D23 system.

### F. Anti-registration is promoted by interleaflet entropy

Finally, we look at the effect of the interleaflet entropic parameter on domain registration characteristics. We made similar plots as above for the 6 systems, of which the one for the 1:1 DPPC-D34 system is shown in Fig.14. As can be seen from the figure, the number of R and PR points decrease as 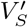 increases, along with a slight increase in PAR and AR points. UR points also show an increase in population as expected. This suggests that the interleaflet entropic interaction disfavors domain registration, and promotes anti-registration. While this is the key result of our work, the effects of the position of unsaturation haven’t been studied experimentally or theoretically before, with the paper by Zhang and Lin[4] being the first in-silico study of the effect of position of unsaturation. We believe that this is a more nuanced look into the effects of position of unsaturation of the lipid tails on domain registration behaviour, taking into account the effects due to overlap of the tail termini and their entropic contribution to the system energy.

**FIG. 14:**
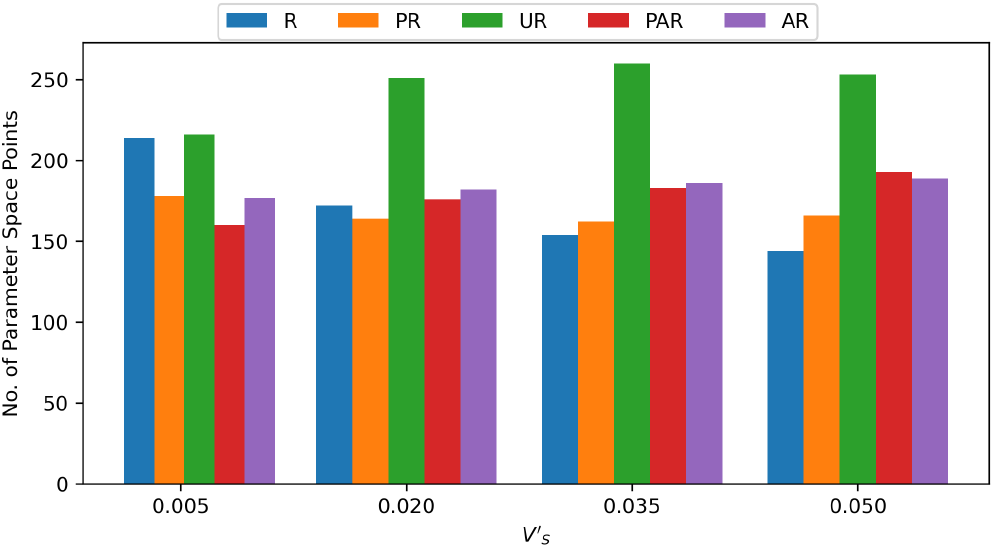
Histograms showing the populations of parameter space points in R, PR, UR, PAR, and AR states at different values of the interleaflet entropic strength contant 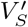 for the 1:1 DPPC-D34 system.

Aside from the above results, by comparing the plots of the 1:1 and 1:3 systems in general, we also find that skewing the ratio of DPPC-D23 or DPPC-D34 away from 1:1 leads to a suppression of domain registration and a significant increase in anti-registration in both 1:3 and 3:1 ratio systems. Also, the plots for the 1:3 and 3:1 systems for both D23 and D34 lipids are quite similar in displaying this trend, suggesting that the increase in anti-registration in this case has to do mostly with the population ratio and not the specific lipids involved. The extent of the change, however, depends on the lipids involved, as we see a more significant dominance of antiregistration in the DPPC-D34 systems with the 1:3 and 3:1 population ratios in their leaflets. There is, however, a slightly higher number of registered points in the 1:3 systems over the 3:1 systems, as can be seen upon inspecting the plots for the 1:3 DPPC-D23 system and the 3:1 DPPC-D23 system, as well as the 1:3 DPPC-D34 system and the 3:1 DPPC-D34 system shown in Fig:S27, Fig:S28, Fig:S29, and Fig:S30 respectively in sec.IX the S.I.

### G. Thermodynamically stable nanodomains

We also found some intriguing results while observing the converged states of some of the points in the parameter space for our systems with skewed population. There were some converged states that showed configurations consisting almost entirely of separate domains that did not grow very large in size, but did not scatter either as the Monte-Carlo simulation progressed. We ran extended MC simulations of 10^9^ moves to ensure that this wasn’t due to a convergence issue, and also saw that these domains persisted throughout the extended simulation. Fig.15 shows a configuration near the end of the extended simulation for the point 0-0-0-2-2-1-4-0-0 in the 1:3 DPPC-D34 system. Some additional points studied are shown in Fig:S31, Fig:S32, and Fig:S33 in the S.I. Also, we have shared a movie file (MovieS1.mp4) of a system that lies in the parameter space exhibiting persistent nanodomains as a part of the S.I. The full set of points that we found are collectively described in the supplementary material.

**FIG. 15:**
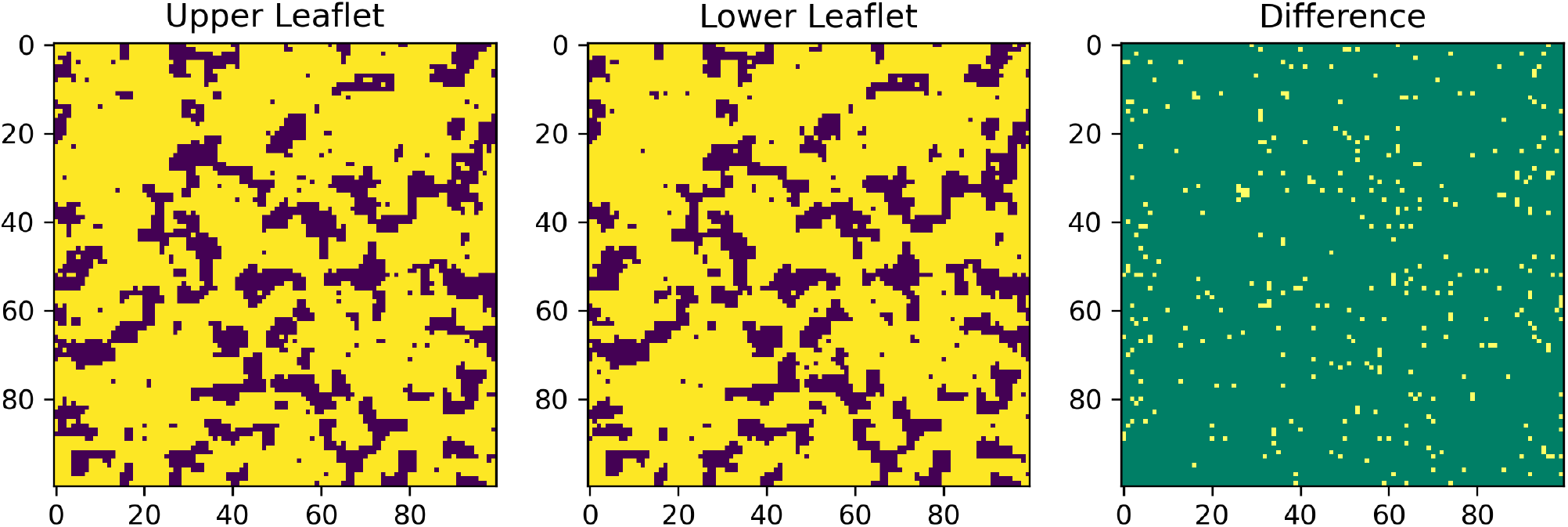
Configuration near the end of the 10^9^ step extended simulation showing the observed nanodomains for the 1:3 DPPC-D34 system at parameter space point 0-0-0-2-2-1-4-0-0. The colours in the plot have the same meaning as in Fig.6.

We also plotted a cumulative domain size distribution over the last 2000 output configurations to more clearly see the spread of sizes of these domains. Fig.16 shows the domain size distribution plot where we can see that there are abundant instances of domains with sizes in the range of around 100 - 500 lipids, which we know from visual inspection of a handful of the configurations aren’t very elongated. The plot is truncated on the left at size 100 because of the large cumulative populations of groups of smaller sizes that exist on the left side of the plot due to its 1/n envelope, where n is the total DPPC population. This plot being for the 1:3 DPPC-D34 system, n is 2500.

**FIG. 16:**
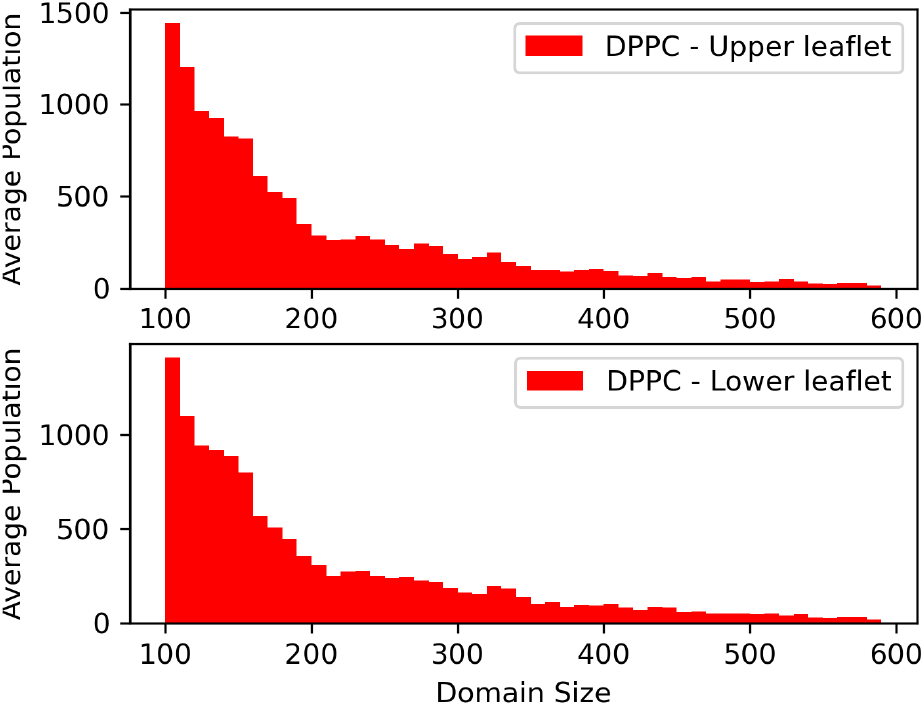
Cumulative domain size distribution plots for DPPC in the upper leaflet and lower leaflet. A significant fraction of the DPPC population lies in domains of sizes ranging from 100-500.

Using typical area per lipid values for DPPC, we can roughly estimate the sizes of these domains, and they fall within the official definition of size of nanodomains of 4-200 nm[63]. With a rough estimate of area per lipid for DPPC to be 64 Å^2^, these domains fall roughly within 9-20 nm in diameter.

The interesting aspect of this observation is that our Hamiltonian only incorporates simple enthalpic and entropic contributions, which suggests that the interplay of the entropic contribution to push towards smaller aggregations as well as the interleaflet enthalpic interaction between registered/anti-registered domains might be sufficient to preserve and evolve small domains, and this could possibly be an aspect that can be further studied in explaining the membrane as a microemulsion of Lo and Ld domains as has been previously studied theoretically[25, 64, 65] along with recent experimental evidence supporting this interpretation of the biological membrane structure[66]. It must be noted that the points where we observe this tend to have stronger interleaflet self interaction, as well as stronger interaction between DPPCs within a leaflet, with weaker D34-D34 and DPPC-D34 interactions. The interaction parameters within the leaflets seem quite justified, and the relative interleaflet enthalpic parameters could form a basis for further investigation of this phenomenon.

## V. DISCUSSION

As a primary result from our work, we find that the interleaflet entropic interaction term promotes antiregistration in the DPPC-D23 and DPPC-D34 systems that we have studied. While there has been no previous targeted study on the effects of position of unsaturation and therefore the entropic contribution due to overlap of tail termini in the interleaflet region on the domain registration behaviour, we believe our work provides a clearer possible explanation to the effect and, with some recent experimental advances[67, 68], provides an opportunity for experimental studies on the same.

We also found that anti-registration was strongly promoted in the systems with the 1:3 and 3:1 DPPC-DX ratio in each leaflet, with domain registration seen only for points with very high values of 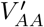 (DPPC-DPPC interleaflet enthalpic interaction strength parameter). In all the six systems, we see that the increase in domain registration due to increase in 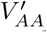 is greater than that due to increase in 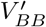 (corresponding to DX-DX interleaflet enthalpic interaction). It is therefore clear that the saturated lipid tail termini in the interleaflet region provide the primary enthalpic driving force for domain registration, whereas the unsaturated lipid tail termini provide an entropic counter that promotes domain antiregistration. This is supported by studies showing that lipid acyl chain dynamics is a key contributor to interleaflet coupling[69] and our results provide a more nuanced explanation for some of the results in previous theoretical[23, 24, 59] and in-vivo studies[60–62] on domain registration in systems with saturated and unsaturated lipids capable of *L_o_* and *L_d_* phase separation.

The enthalpic contribution of the unsaturated lipids is also important, however, in increasing the coupling between leaflets. We see from the increased fraction of R and AR points in the DPPC-D34 systems that the higher D34-D34 interleaflet enthalpic interaction compared to that of D23-D23 allows the system to more easily converge to registered or anti-registered states. This is more difficult in the DPPC-D23 systems, as a weaker coupling between leaflets increases fluctuations in the interleaflet region, thereby showing a more spread out categorization of registration states. This illustrates that the function of the lower position of unsaturation is, therefore, to simply improve coupling between the leaflets both through enthalpic and entropic contributions, and that the system converges to a registered or anti-registered state depending on the relative enthalpic and entropic contributions, which can vary significantly based on the specific lipids in the system.

We think that the behaviour demonstrated in the paper by Zhang and Lin[4] can be explained by our interpretation of stronger interleaflet coupling in the DPPC-D34 system as compared to the DPPC-D23 system due to a lower position of unsaturation in the D34 lipids instead of their original explanation that suggested that the D23 lipids had interleaflet interactions closer to that of the saturated lipid DPPC, leading to lesser enthalpic driving force for registration. We predict that the number and size of packing defects in the interleaflet region would be higher in a system where the unsaturated lipid has a higher position of unsaturation, and that there will be better packing in a system where the unsaturated lipid has a lower position of unsaturation[70, 71]. Since the increase in interleaflet entropic interaction strength 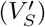 can also be loosely interpreted as a rise in temperature the way our Hamiltonian is defined, our observation that lower values of 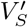 allow greater number of domain registered points in parameter space also demonstrates the better enthalpic coupling seen in systems at lower temperatures.

Also, from the large parameter space plots in sec. XI and sec.XII, we find that phase-separation and antiregistration are strongly correlated when 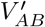 is high, whereas phase separation and registration are correlated when 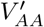 and 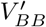 are high and 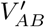 is low. This indicates that the registration and anti-registration tendencies of a domain can influence the phase-separation in the other leaflet. This has also been observed in multiple theoretical studies earlier[23, 24, 64] and further supports the feasibility of our model. Additionally, we observe the presence of thermodynamically stable nanodomains with our curvature-independent Hamiltonian. These observations are in line with the direct, probe-free imaging of model and bioderived membrane using cryogenic electron microscope[72] and fluorescence resonance energy transfer (FRET) and, electron spin resonance (ESR) studies on model quaternary lipid systems[66], which provide evidence towards the existence of nanodomains in cell membranes. The experimental evidences are supported by theoretical studies by DW Allender and M Schick[25, 65] where the authors suggested that inhomogeneity in biological membranes might exist in the form of a microemulsion of Lo and Ld regions. They attribute this to spontaneous curvature of the membrane, bending modulus, surface tension as well as the energy cost of formation of concentration gradients. We believe our observation supports the microemulsion hypothesis and provide a possible direction of further investigation.

This study primarily focused on the position of unsaturation, and not on factors like tail interdigitation that have also been shown to affect interleaflet coupling. We avoided this, as we already had too many parameters in the Hamiltonian to work with. However, for the lipids that we worked with, it shouldn’t be very significant as the lengths of the lipids were roughly similar. The DPPC-D23 and DPPC-D34 systems are very good model systems to isolate the effect of the difference in position of unsaturation due to their similarity in all other aspects.

## VI. CONCLUSIONS

In this study, we have demonstrated that a physics based model accounting for the position of unsaturation in lipid tails can successfully capture domain registration/anti-registration behaviour of lipid membranes, and directly provided verification and a more complete explanation for the results observed by Zhang and Lin[4]. Our work suggests that the lower the position of unsaturation, the better the coupling. Whether the system equilibrates to a registered or anti-registered state or something in between depends on the relative enthalpic and entropic contribution to this interleaflet coupling from the interactions of lipid tail termini in the interleaflet region. The lipids with lower positions of unsaturation improve coupling due to them having a larger number of configurations where their tail termini are in the interleaflet region and not bent away. This is an important leap in the understanding of the problem, as it shows us that it is not simply the enthalpic interaction in the interleaflet region that drives domain registration, and that the entropic contributions are essential in explaining the domain registration behaviour observed in the model systems used in Zhang and Lin’s work[4]. It also supports the idea of a need for a direct coupling between leaflets in order to observe domain registration behaviour, as suggested in many of the earlier theoretical articles on domain registration[1, 23–25]. However, it doesn’t discredit previous work that claim indirect coupling mechanisms suggesting line tension, local curvature and domain stiffness as the primary mechanisms of domain registration[3, 26–28].

We believe that the overall phenomenon of domain registration is a combination of all the facets currently hypothesised in some proportions depending on the exact lipid compositions and their individual structures and tendencies, and our work serves to add some much needed clarification on the facet of position lipid tail unsaturations.

## SUPPORTING MATERIAL

1. supporting_information.pdf : supporting information
2. movieS1.avi : Persistent registered nanodomain movie

## AUTHOR CONTRIBUTIONS

Anand Srivastava conceived the idea and designed the experiments in consultation with Akshara Sharma. Aniruddha Seal and Sahithya Iyer developed the protocol for creating the artificial lipid bilayer systems under considerations and set up and ran the all-atom simulations. Anand Srivastava and Akshara Sharma formulated the Hamiltonian functions and Akshara Sharma carried out the Monte Carlo simulation and all the analyses. Akshara Sharma and Anand Srivastava wrote the paper with help of other authors.

## ACKNOWLEDGMENTS

Anand Srivastava (A.S.) would like to thank Prof. Madan Rao for discussions. Financial support from the Indian Institute of Science-Bangalore and the high-performance computing facility ”Beagle” setup from grants by a partnership between the Department of Biotechnology of India and the Indian Institute of Science (IISc-DBT partnership programme) are greatly acknowledged. A.S. thanks the startup grant provided by the Ministry of Human Resource Development of India and the early career grant from the Department of Science and Technology of India. A.S. also thanks the DST for the National Supercomputing Mission grant. FIST program sponsored by the Department of Science and Technology and UGC, Centre for Advanced Studies and Ministry of Human Resource Development, India is gratefully acknowledged by the authors.

## Supplementary Information

### I. SCHEMATICS OF THE WORKFLOW

Fig.1 shows the overall workflow followed in the project. Starting from the hypothesis, we first carried out the all-atom simulations with the appropriate lipid structures for the D23 and D34 lipids to supply the site-variable to the lattice bilayer Hamiltonian that was developed in parallel. Following this, the parametric study was conducted using this input while varying the other parameters of the Hamiltonian to observe the qualitative trends shown with the variation of each parameter. Finally, we draw conclusions from these trends and interpret the analysed data which is compared to existing results to validate our model.

### II. TIPS AND TROUBLESHOOTING FOR THE AA SYSTEM GENERATION

This section contains some extra steps that might be required, as learnt from our experience during the development of the procedure.

- To make asymmetric bilayers, one may make two bilayers using this tool, each of different compositions as required, and then fuse together a leaflet of each.
- The method can require some troubleshooting in the case of appearance of artificial voids. To avoid this, we must visualize the structure obtained and if we see artificial voids in the structure, we must prepare a smaller patch of the system and then replicate it along X and Y axis to form the final patch. If the voids still persist, the leaflets need to be moved close to each other, until they overlap.
- The final membrane bilayer structure formed after patching usually shows bad contacts among the waters in the system. Hence, we remove these water molecules, after which we re-solvate the bilayer and remove the water molecules which have penetrated below the headgroups[1],[2] and proceed to the next steps with this new structure. One must also make sure to update the topology file after this.
- A short NPT equilibration (5 ns worked for us) might be required to be performed on the energy-minimized structure because the structure of the lipids obtained after the energy minimization is not in their native state. So, running an NVT Equilibration on this system directly can make the system blow up and separate the leaflets on either side of the simulation box.

**FIG. 1:**
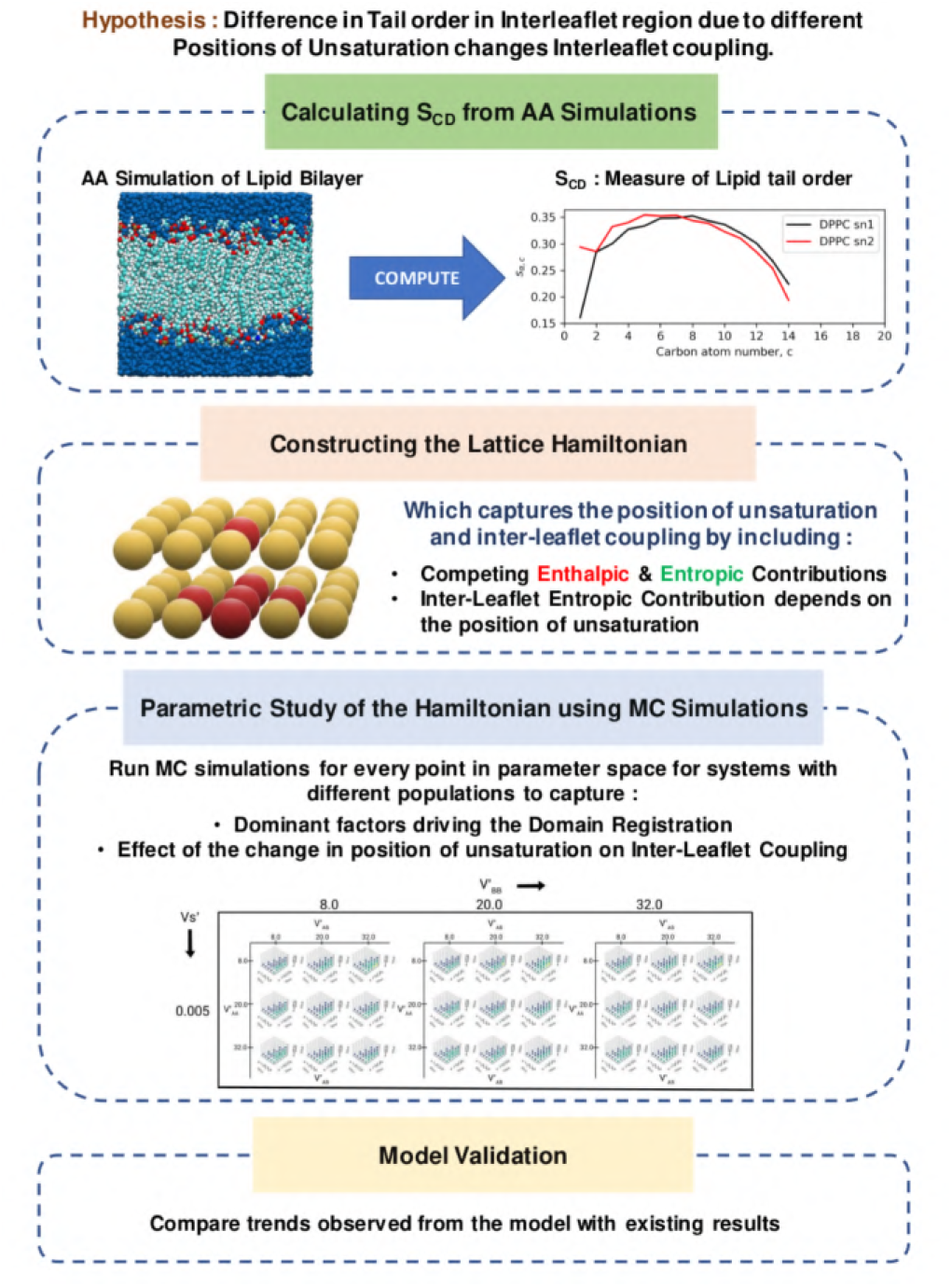
A schematic showing the workflow adopted in this project.

### III. CONVERGENCE OF MC SIMULATIONS

In this section, we briefly discuss the convergence of the MC simulations for the various points in parameter space of the six systems studied. Figure.2 shows the energy plots with respect to number of MC moves of four arbitrarily chosen parameter space points from each system. We can see that the systems have reached a stable energy with an acceptably small variance in energy values. The variation of the energy for all points is within 1% of the final total system energy at that point in parameter space.

**FIG. 2:**
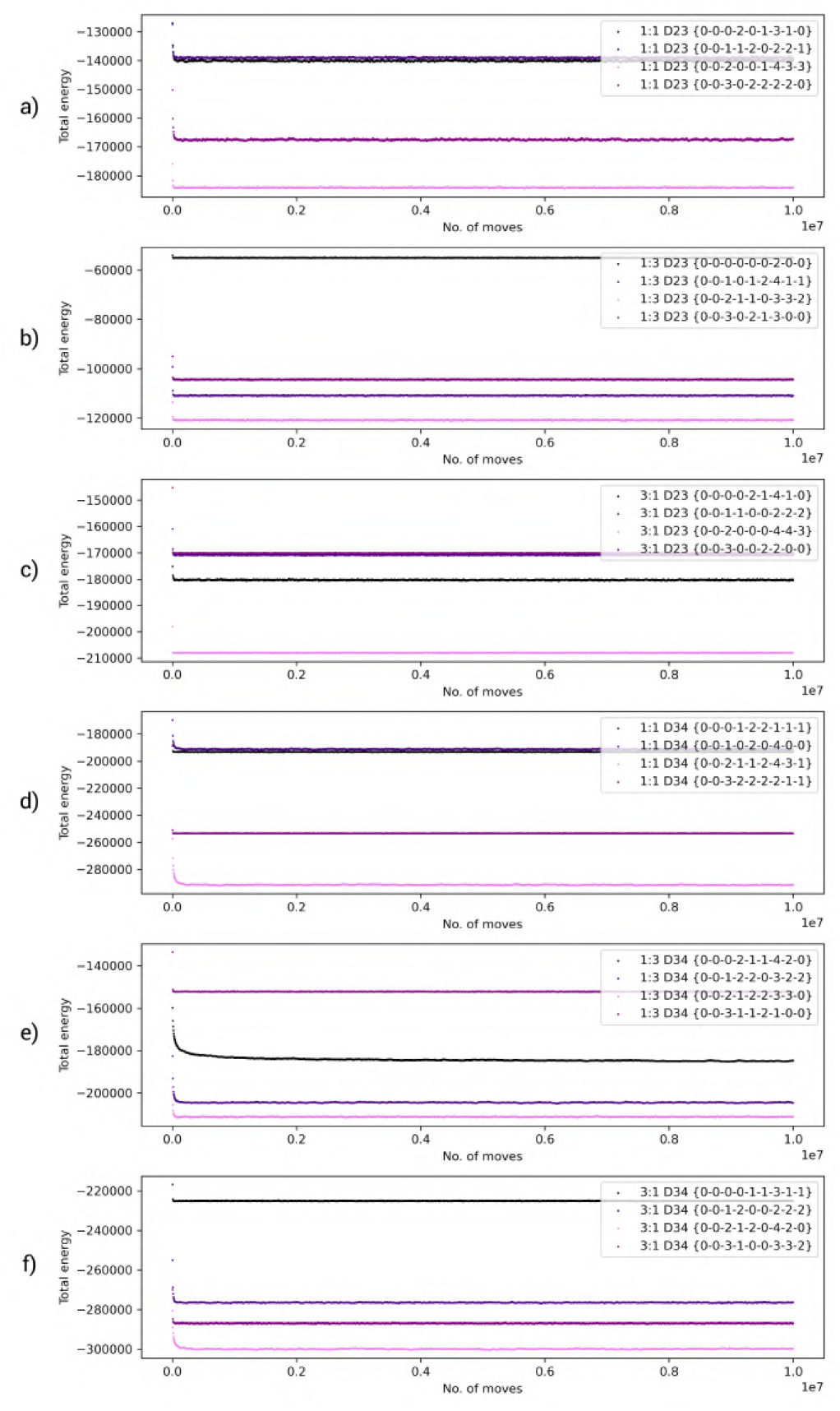
Total system energy vs. number of MC moves plotted for 4 points from each of the 6 systems, a) 1:1 DPPC-D23 systems, b) 1:3 DPPC-D23 systems, c) 3:1 DPPC-D23 systems, d) 1:1 DPPC-D34 systems, e) 1:3 DPPC-D34 systems, f) 3:1 DPPC-D34 systems. As can be seen, the energy converges quickly and remains steady.

Although we haven’t plotted the energy vs moves curve for every point in parameter space, we have verified manually that systems with all the different types of final configurations produced during our study indeed show acceptable variation in energy near the end of their simulations.

### IV. PHASE SEPARATION HISTOGRAMS FOR THE LATERAL ENTHALPIC PARAMETERS

This section shows the histograms showing the populations of PS, PPS, and NPS states at different values of the lateral enthalpic parameters *V_AA_, V_BB_*, and *V_AB_* for the systems excluded from the main manuscript in section IV.A, namely the 1:1 DPPC-D23 system in fig.3, the 1:3 DPPC-D23 system in fig.4, the 3:1 DPPC-D23 system in fig.5, and the 3:1 DPPC-D34 system in fig.6.

**FIG. 3:**
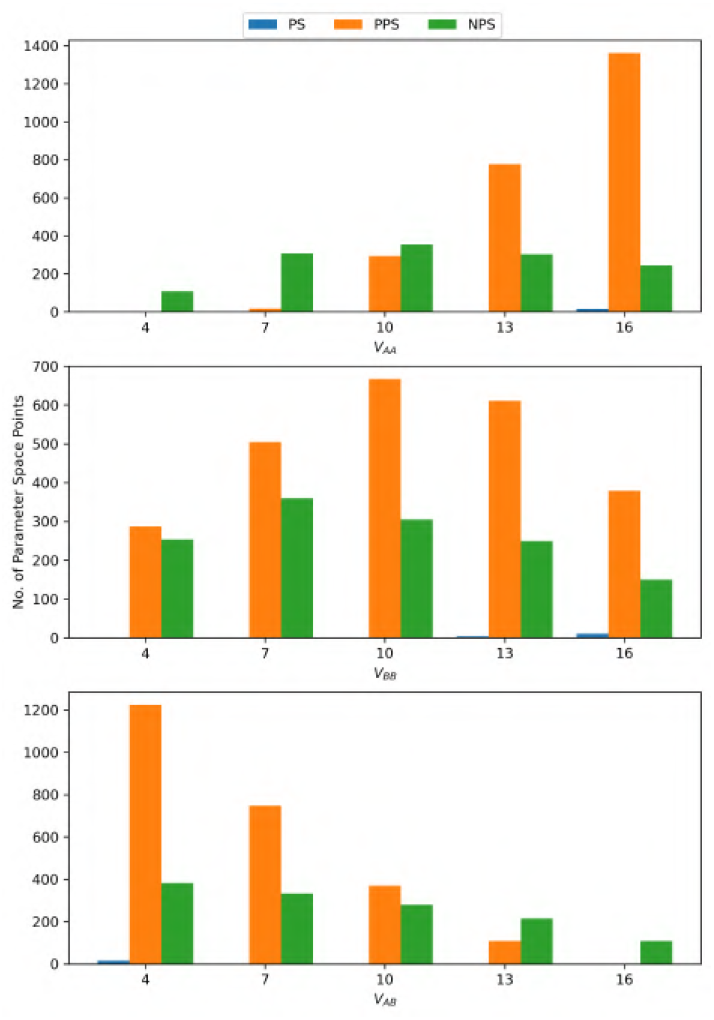
Histograms showing the populations of parameter space points in PS, PPS, and NPS states at different values of the lateral enthalpic interaction strength constants *V_AA_, V_BB_*, and *V_AB_* for the 1:1 DPPC-D23 system.

**FIG. 4:**
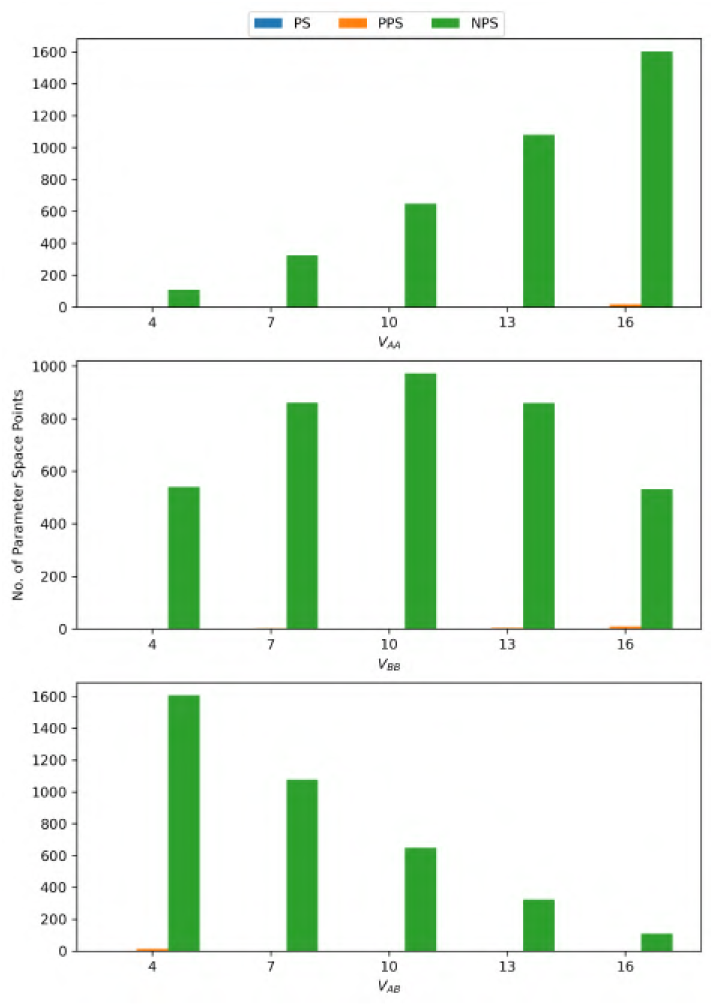
Histograms showing the populations of parameter space points in PS, PPS, and NPS states at different values of the lateral enthalpic interaction strength constants *V_AA_, V_BB_*, and *V_AB_* for the 1:3 DPPC-D23 system.

**FIG. 5:**
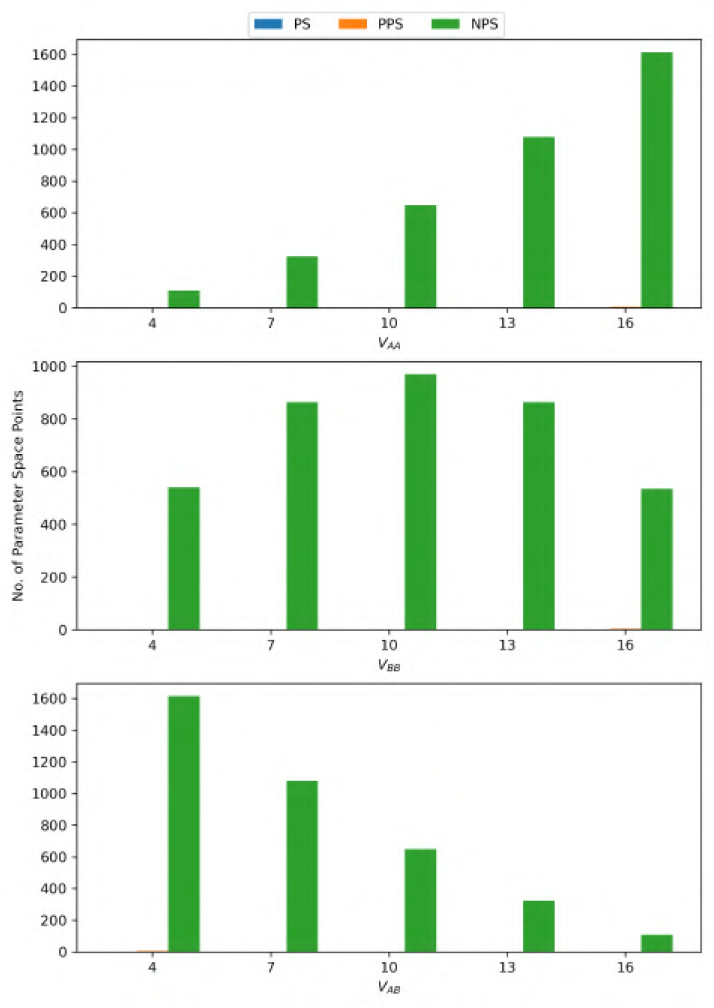
Histograms showing the populations of parameter space points in PS, PPS, and NPS states at different values of the lateral enthalpic interaction strength constants *V_AA_, V_BB_*, and *V_AB_* for the 3:1 DPPC-D23 system.

**FIG. 6:**
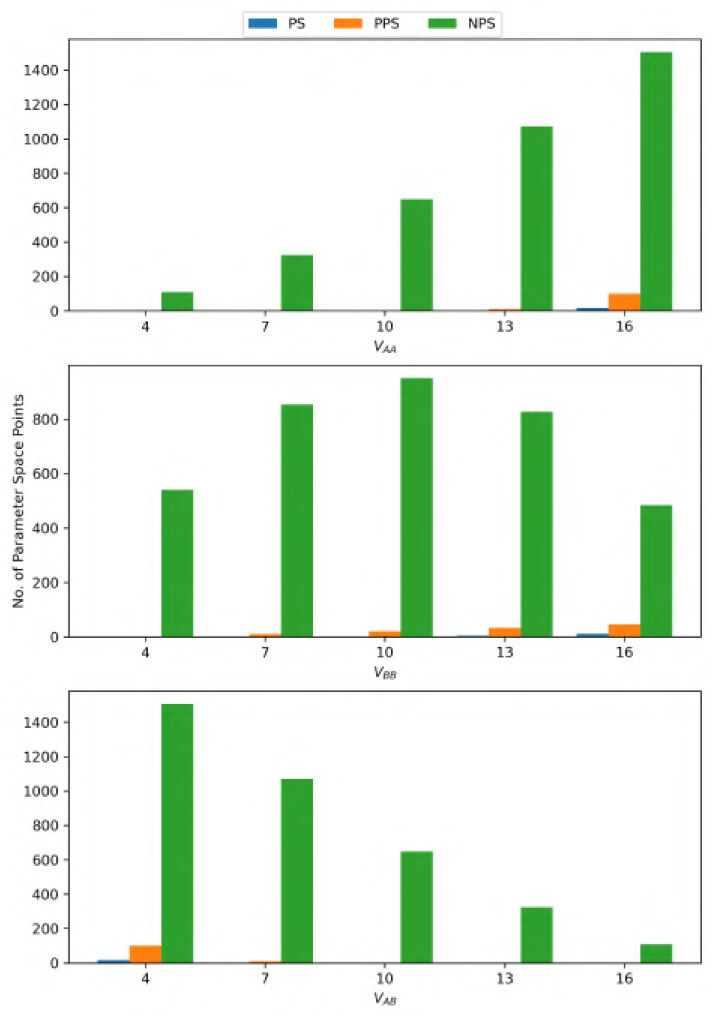
Histograms showing the populations of parameter space points in PS, PPS, and NPS states at different values of the lateral enthalpic interaction strength constants *V_AA_, V_BB_*, and *V_AB_* for the 3:1 DPPC-D34 system.

### V. PHASE SEPARATION HISTOGRAMS FOR THE INTERLEAFLET ENTHALPIC PARAMETERS

This section shows the histograms showing the populations of PS, PPS, and NPS states at different values of the interleaflet enthalpic parameters 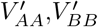, and 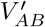 for the systems excluded from the main manuscript in section IV.B, namely the 1:1 DPPC-D23 system in fig.7, the 1:3 DPPC-D23 system in fig.8, the 3:1 DPPC-D23 system in fig.9, the 1:3 DPPC-D34 system in Fig.10, and the 3:1 DPPC-D34 system in Fig. 11.

**FIG. 7:**
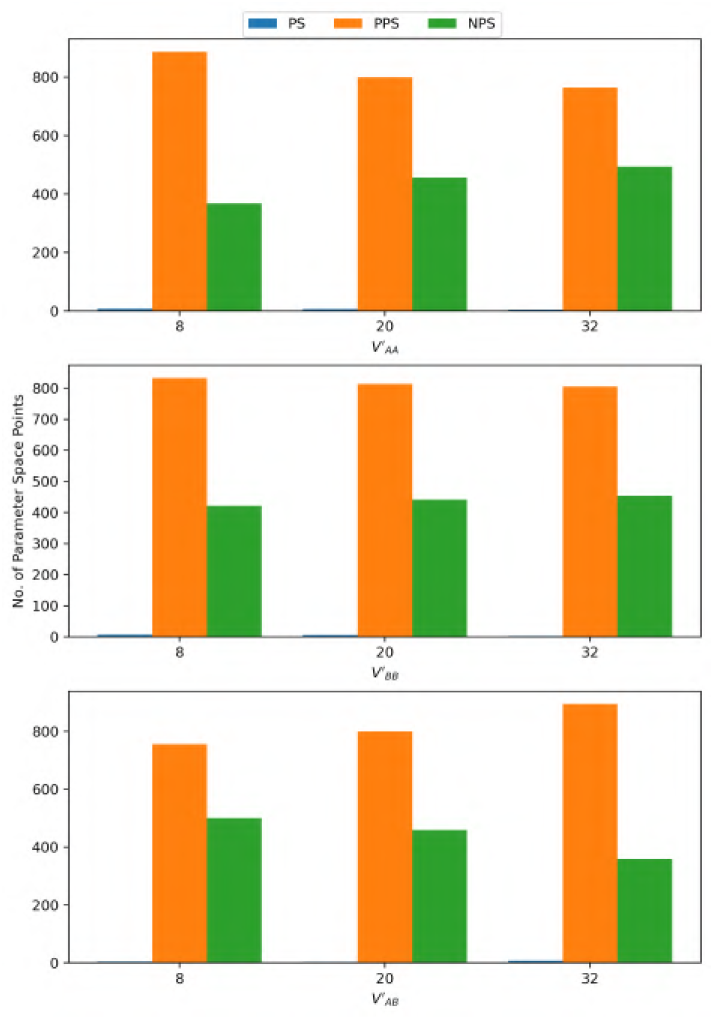
Histograms showing the populations of parameter space points in PS, PPS, and NPS states at different values of the interleaflet enthalpic interaction strength constants 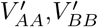, and 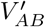 for the 1:1 DPPC-D23 system.

**FIG. 8:**
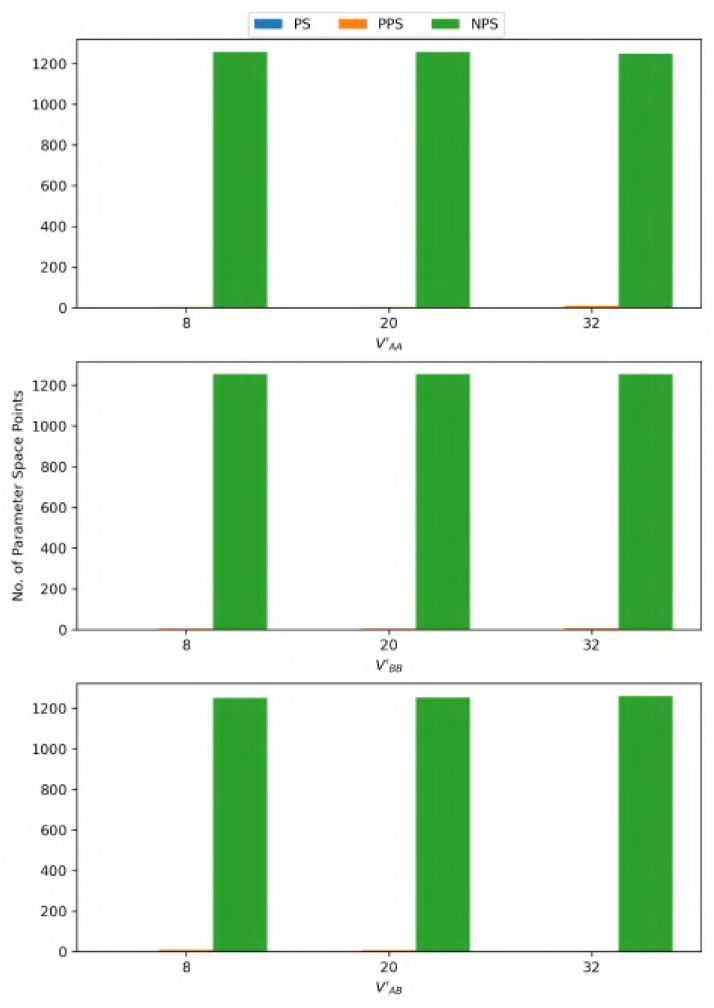
Histograms showing the populations of parameter space points in PS, PPS, and NPS states at different values of the interleaflet enthalpic interaction strength constants 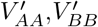, and 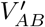 for the 1:3 DPPC-D23 system.

**FIG. 9:**
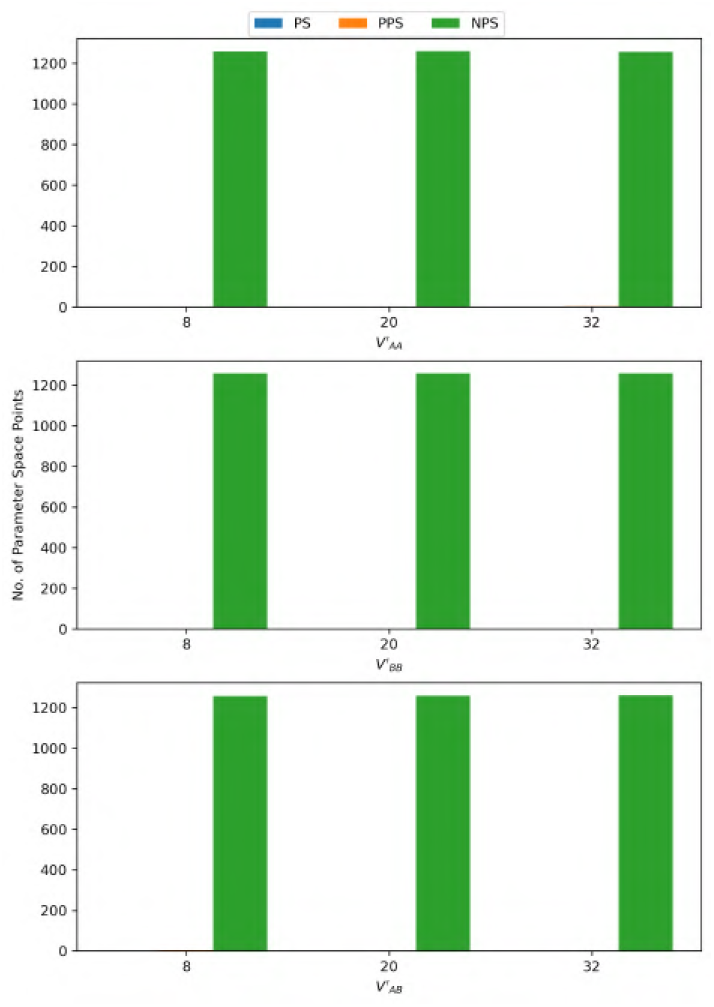
Histograms showing the populations of parameter space points in PS, PPS, and NPS states at different values of the interleaflet enthalpic interaction strength constants 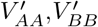, and 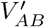 for the 3:1 DPPC-D23 system.

**FIG. 10:**
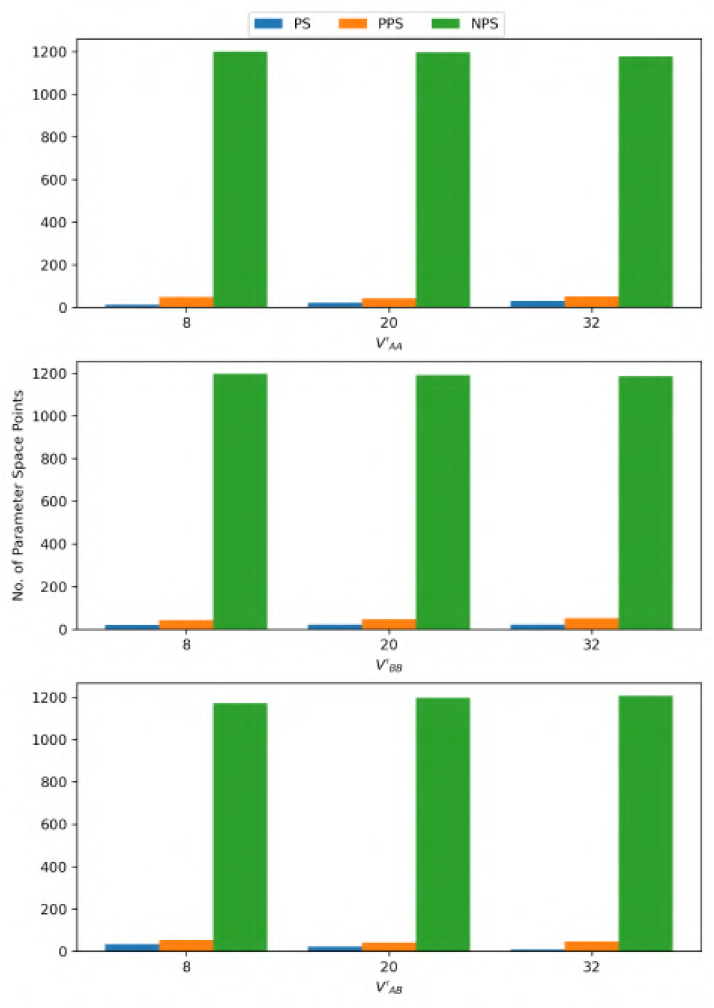
Histograms showing the populations of parameter space points in PS, PPS, and NPS states at different values of the interleaflet enthalpic interaction strength constants 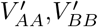, and 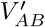 for the 1:3 DPPC-D34 system.

**FIG. 11:**
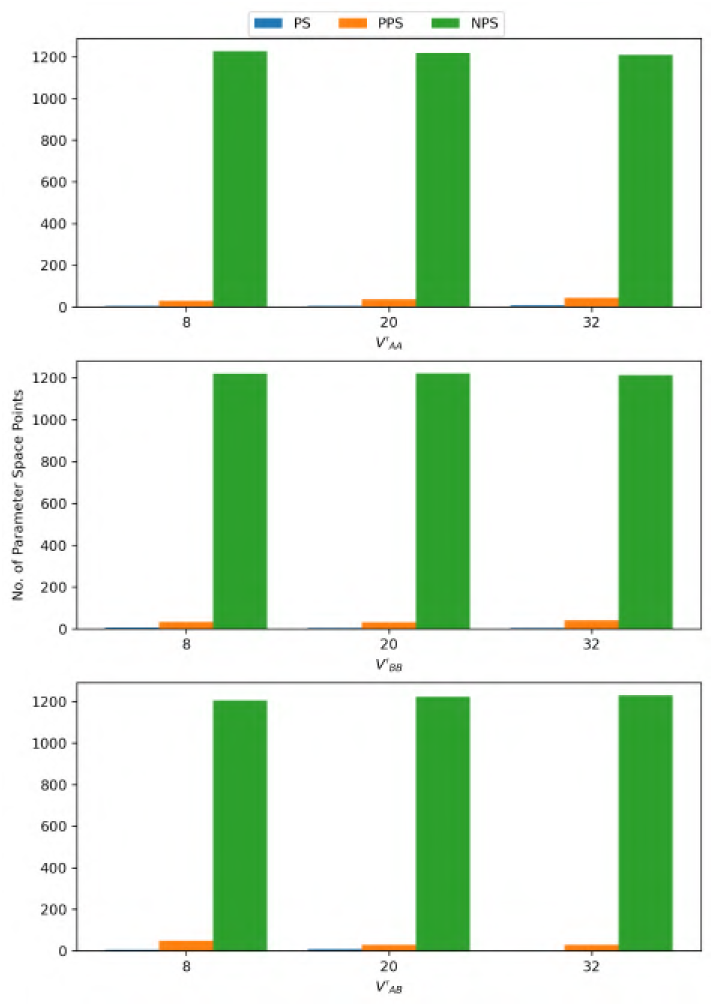
Histograms showing the populations of parameter space points in PS, PPS, and NPS states at different values of the interleaflet enthalpic interaction strength constants 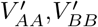, and 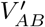 for the 3:1 DPPC-D34 system.

### VI. PHASE SEPARATION HISTOGRAMS FOR THE INTERLEAFLET ENTROPIC STRENGTH PARAMETER

This section shows the histograms showing the populations of PS, PPS, and NPS states at different values of the interleaflet entropic strength parameter 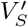 for the systems excluded from the main manuscript in section IV.C, namely the 1:1 DPPC-D23 system in fig.12, the 1:3 DPPC-D23 system in fig.13, the 3:1 DPPC-D23 system in fig.14, the 1:3 DPPC-D34 system in Fig.15, and the 3:1 DPPC-D34 system in Fig.16.

**FIG. 12:**
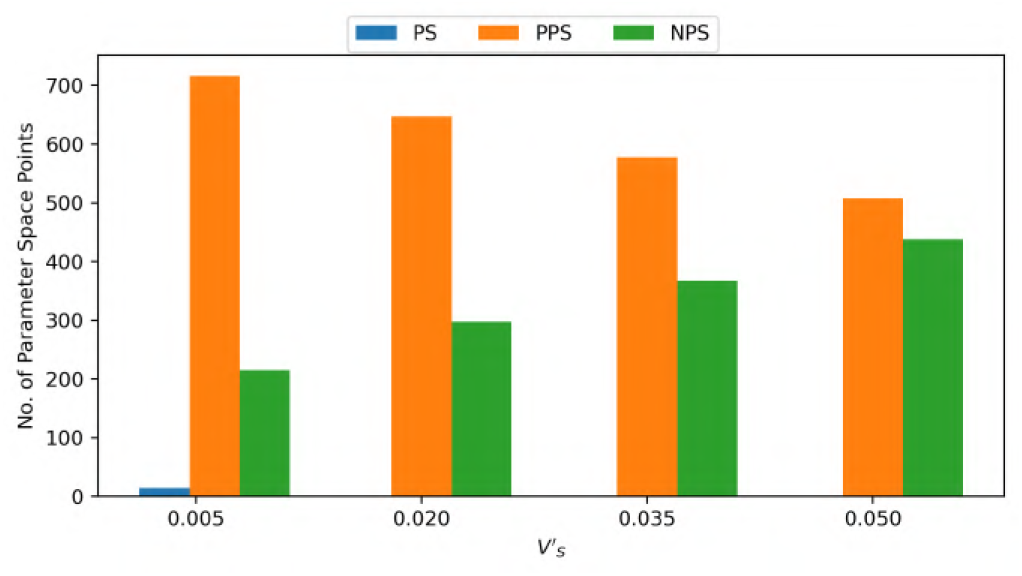
Histograms showing the populations of parameter space points in PS, PPS, and NPS states at different values of the interleaflet entropic strength contant 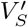 for the 1:1 DPPC-D23 system.

**FIG. 13:**
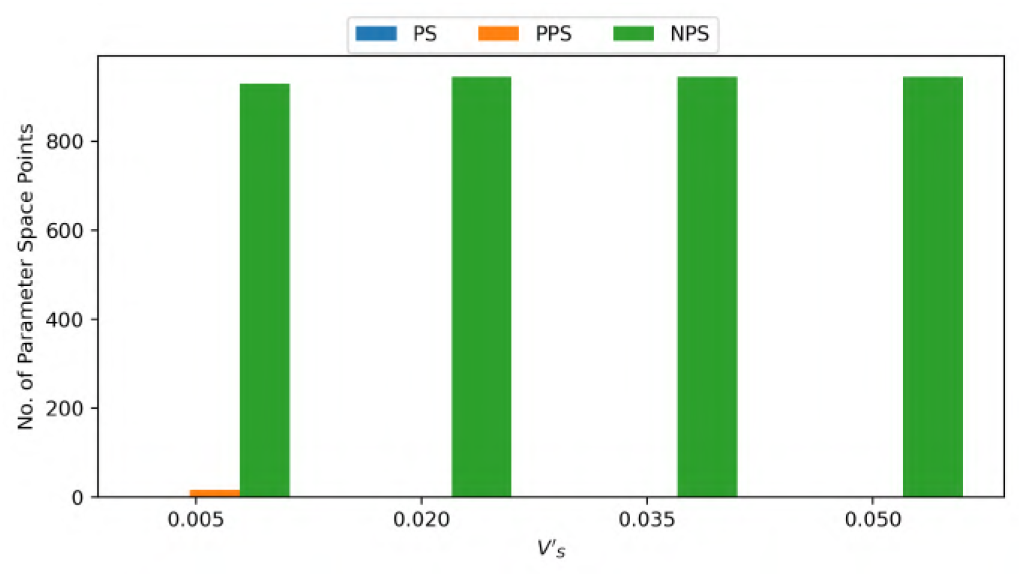
Histograms showing the populations of parameter space points in PS, PPS, and NPS states at different values of the interleaflet entropic strength contant 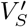 for the 1:3 DPPC-D23 system.

**FIG. 14:**
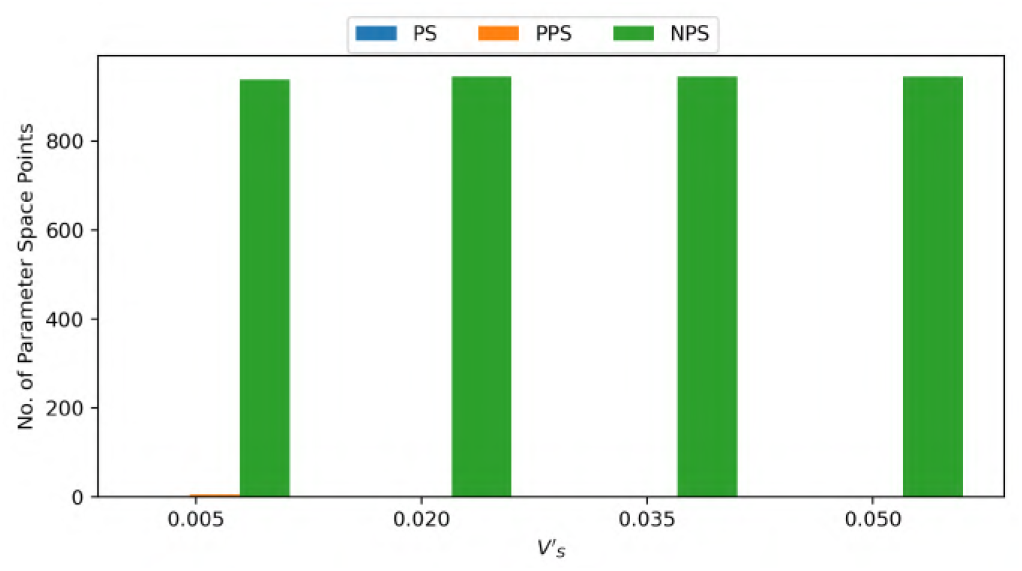
Histograms showing the populations of parameter space points in PS, PPS, and NPS states at different values of the interleaflet entropic strength contant 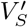 for the 3:1 DPPC-D23 system.

**FIG. 15:**
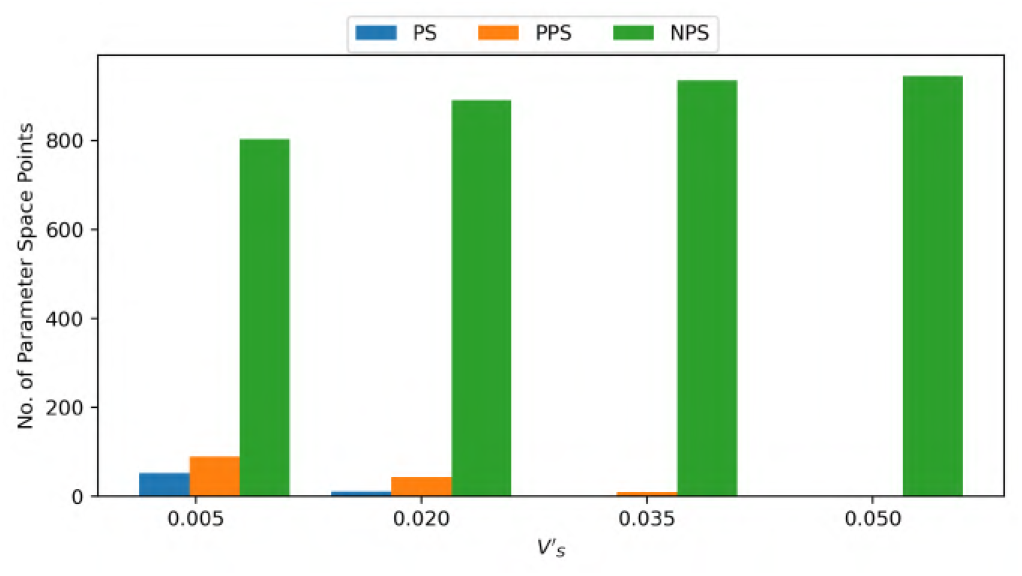
Histograms showing the populations of parameter space points in PS, PPS, and NPS states at different values of the interleaflet entropic strength contant 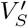 for the 1:3 DPPC-D34 system.

**FIG. 16:**
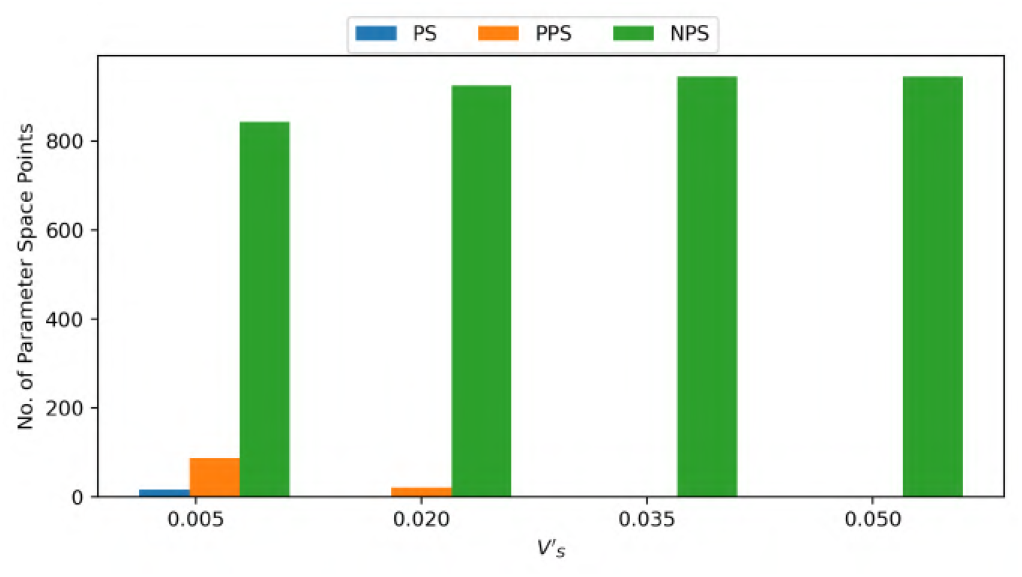
Histograms showing the populations of parameter space points in PS, PPS, and NPS states at different values of the interleaflet entropic strength contant 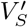 for the 3:1 DPPC-D34 system.

### VII. DOMAIN REGISTRATION HISTOGRAMS FOR THE LATERAL ENTHALPIC PARAMETERS

This section shows the histograms showing the populations of R, PR, UR, PAR, and AR states at different values of the lateral enthalpic parameters *V_AA_, V_BB_*, and *V_AB_* for the systems excluded from the main manuscript in section IV.D, namely the 1:1 DPPC-D23 system in fig.17, the 1:3 DPPC-D23 system in fig.18, the 3:1 DPPC-D23 system in fig.19, the 1:3 DPPC-D34 system in fig.20, and the 3:1 DPPC-D34 system in fig.21.

**FIG. 17:**
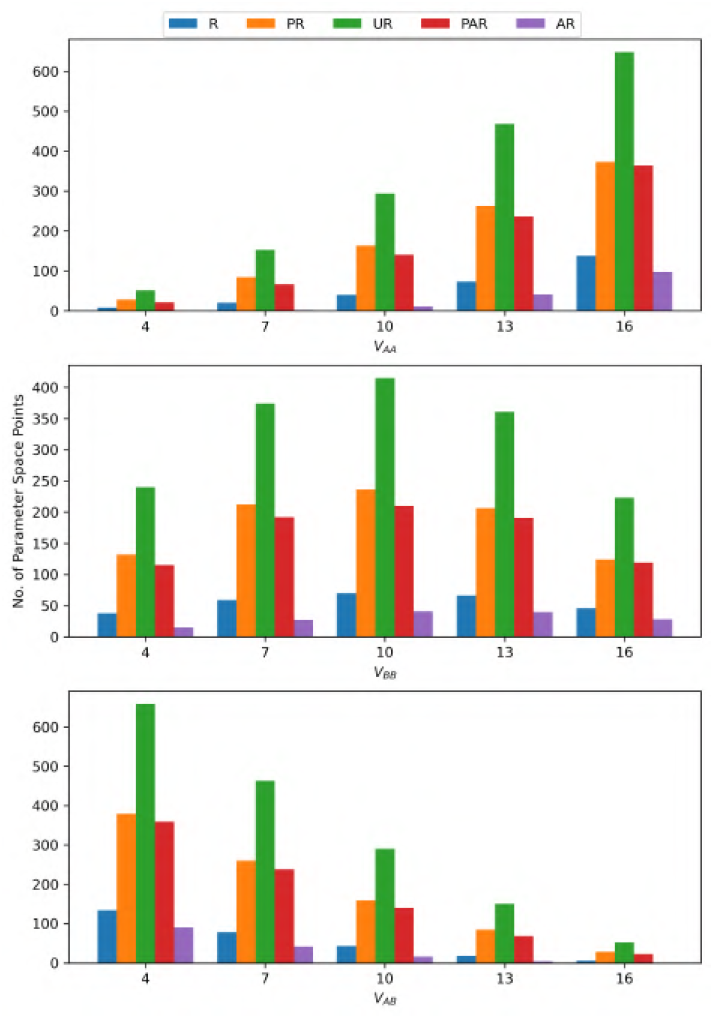
Histograms showing the populations of parameter space points in R, PR, UR, PAR, and AR states at different values of the lateral enthalpic interaction strength constants *V_AA_, V_BB_*, and *V_AB_* for the 1:1 DPPC-D23 system.

**FIG. 18:**
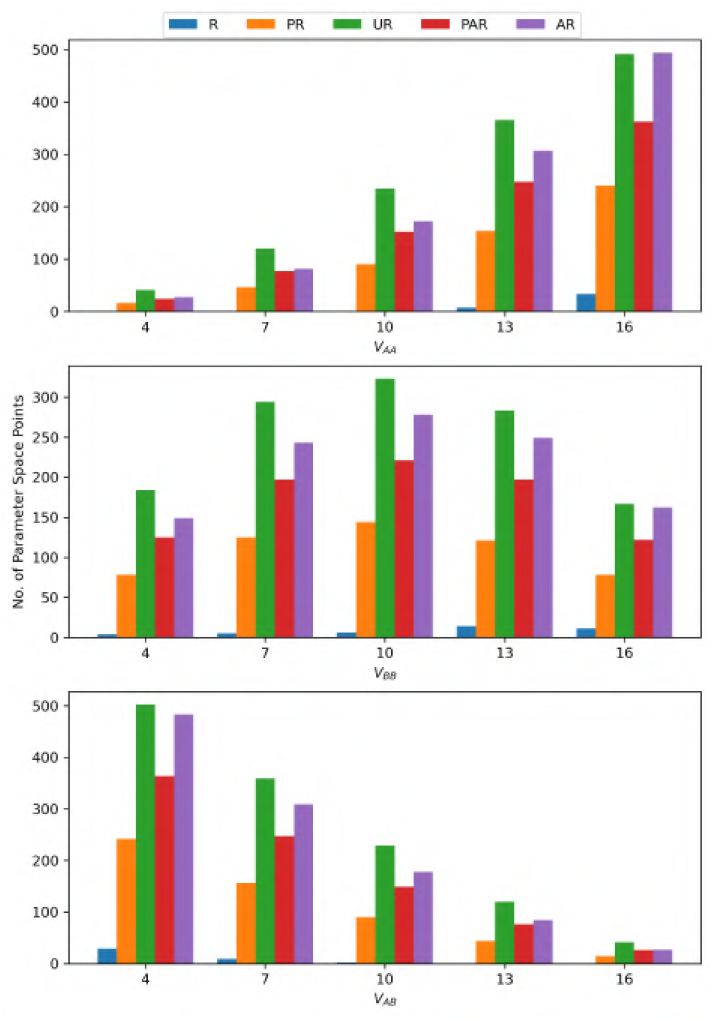
Histograms showing the populations of parameter space points in R, PR, UR, PAR, and AR states at different values of the lateral enthalpic interaction strength constants *V_AA_, V_BB_*, and *V_AB_* for the 1:3 DPPC-D23 system.

**FIG. 19:**
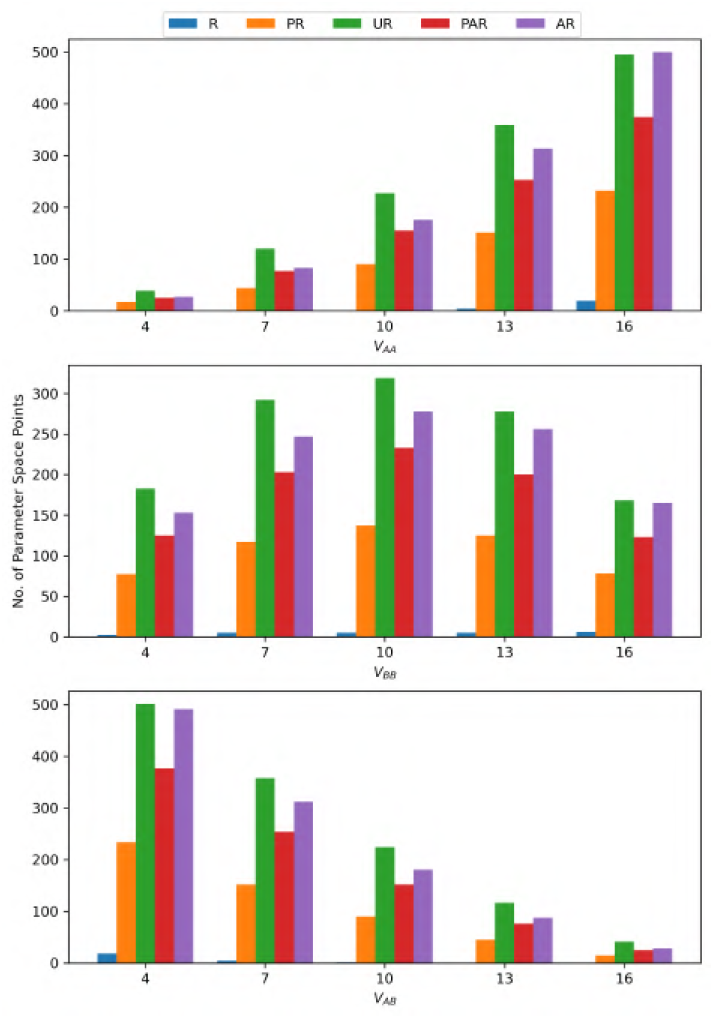
Histograms showing the populations of parameter space points in R, PR, UR, PAR, and AR states at different values of the lateral enthalpic interaction strength constants *V_AA_, V_BB_*, and *V_AB_* for the 3:1 DPPC-D23 system.

**FIG. 20:**
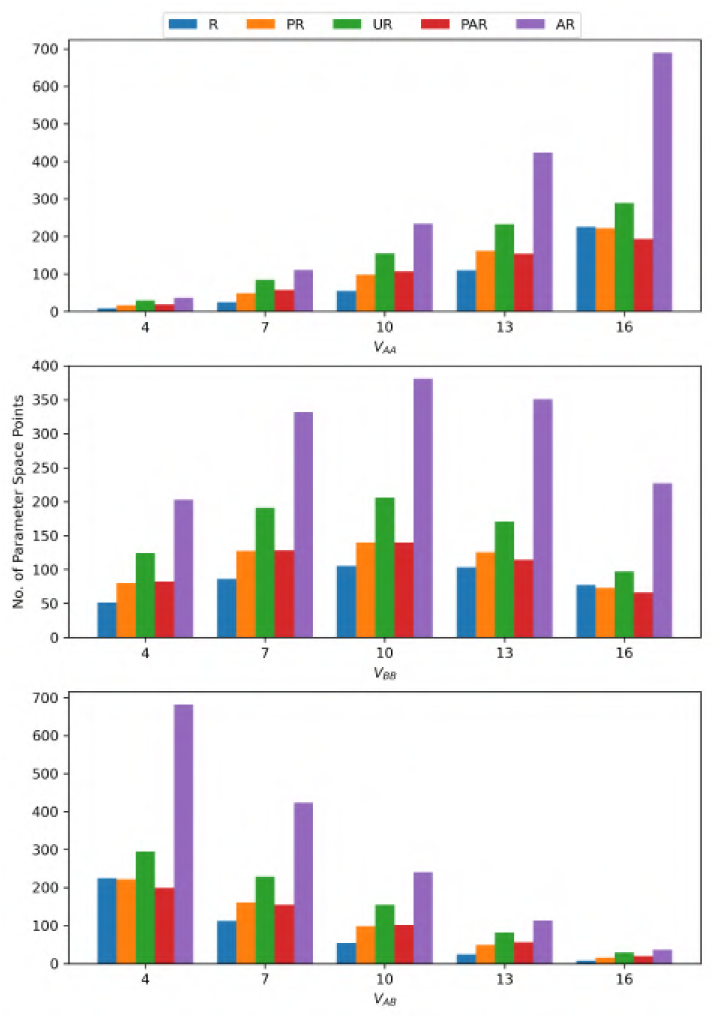
Histograms showing the populations of parameter space points in R, PR, UR, PAR, and AR states at different values of the lateral enthalpic interaction strength constants *V_AA_, V_BB_*, and *V_AB_* for the 1:3 DPPC-D34 system.

**FIG. 21:**
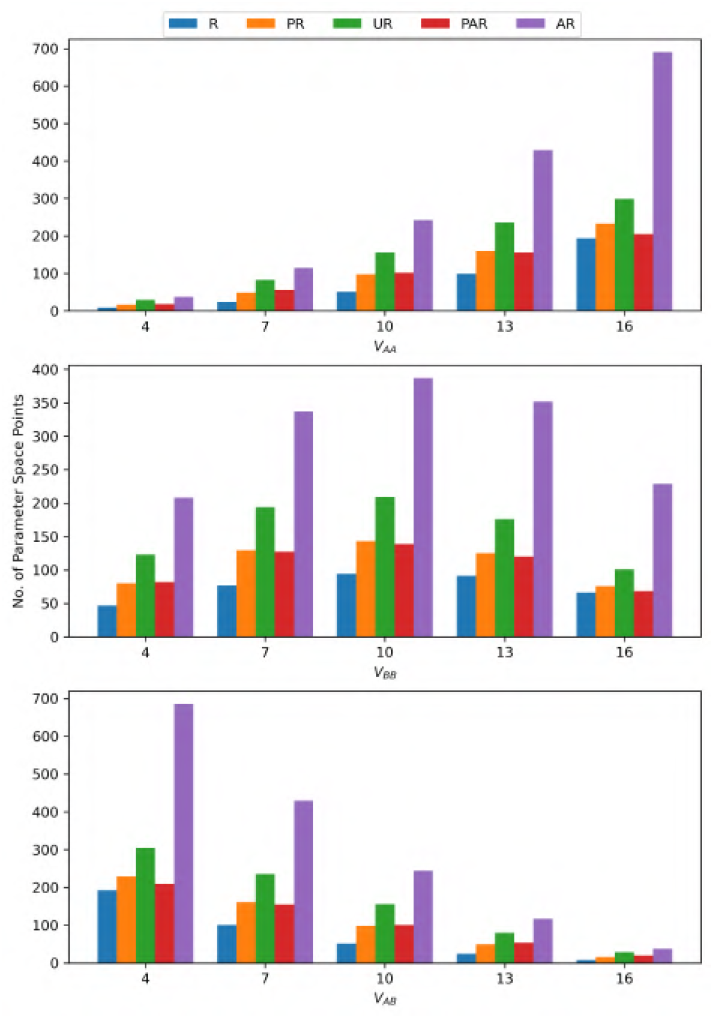
Histograms showing the populations of parameter space points in R, PR, UR, PAR, and AR states at different values of the lateral enthalpic interaction strength constants *V_AA_, V_BB_*, and *V_AB_* for the 3:1 DPPC-D34 system.

### VIII. DOMAIN REGISTRATION HISTOGRAMS FOR THE INTERLEAFLET ENTHALPIC PARAMETERS

This section shows the histograms showing the populations of R, PR, UR, PAR, and AR states at different values of the lateral enthalpic parameters 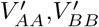, and 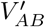 for the systems excluded from the main manuscript in section IV.E, namely the 1:3 DPPC-D23 system in fig.22, the 3:1 DPPC-D23 system in fig.23, the 1:3 DPPC-D34 system in fig.24, and the 3:1 DPPC-D34 system in fig.25.

**FIG. 22:**
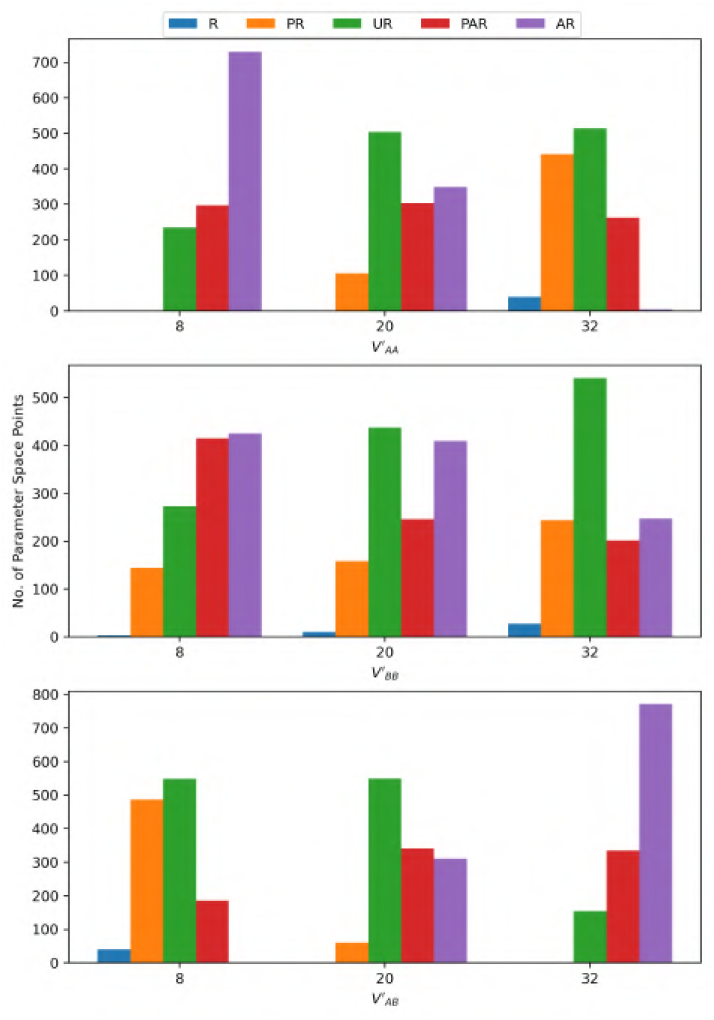
Histograms showing the populations of parameter space points in R, PR, UR, PAR, and AR states at different values of the interleaflet enthalpic interaction strength constants 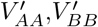, and 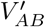 for the 1:3 DPPC-D23 system.

**FIG. 23:**
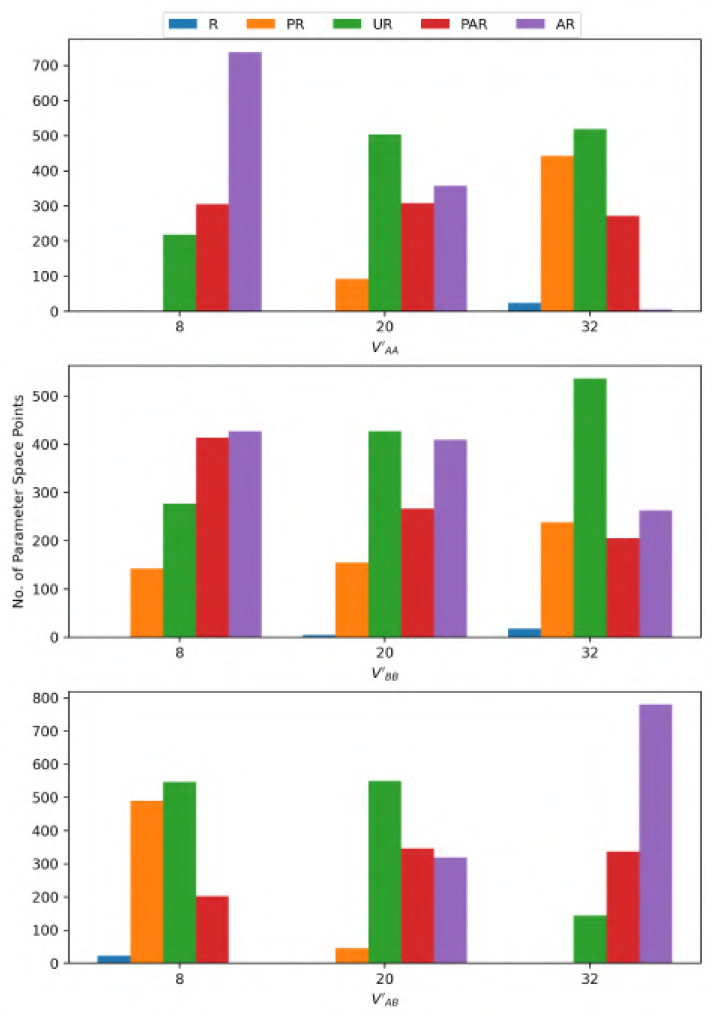
Histograms showing the populations of parameter space points in R, PR, UR, PAR, and AR states at different values of the interleaflet enthalpic interaction strength constants 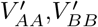, and 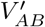 for the 3:1 DPPC-D23 system.

**FIG. 24:**
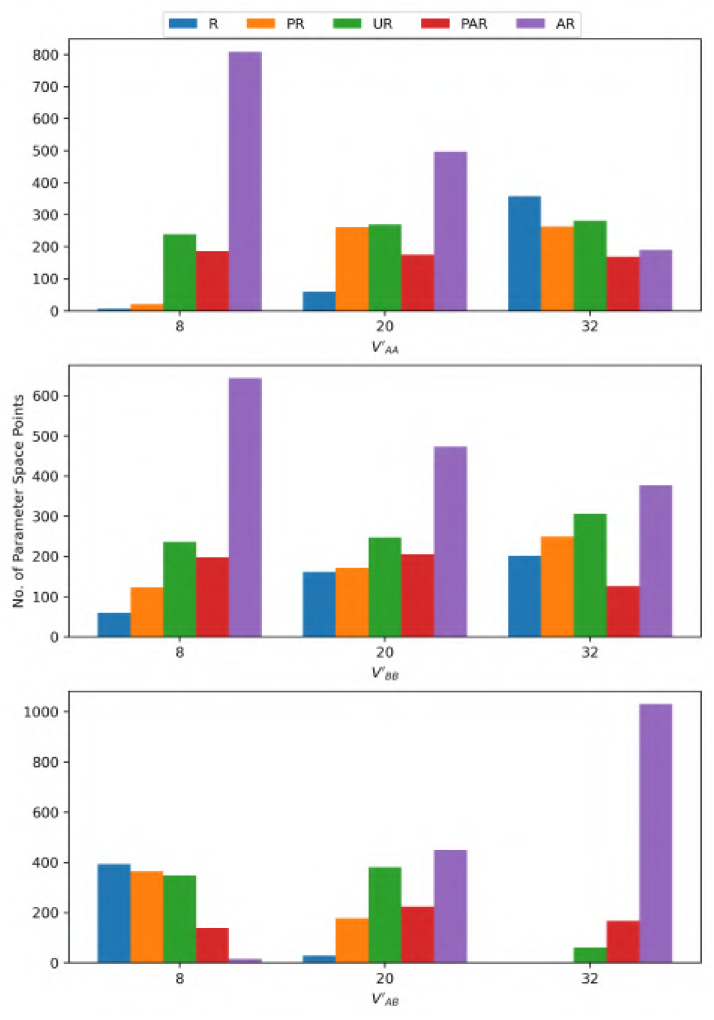
Histograms showing the populations of parameter space points in R, PR, UR, PAR, and AR states at different values of the interleaflet enthalpic interaction strength constants 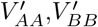, and 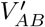 for the 1:3 DPPC-D34 system.

**FIG. 25:**
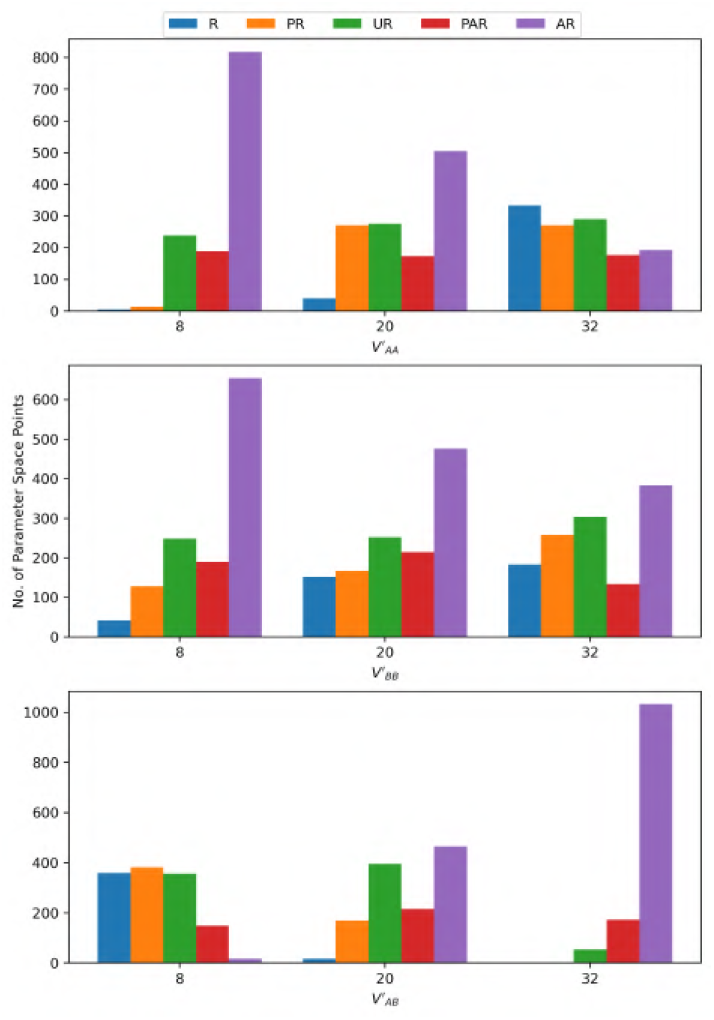
Histograms showing the populations of parameter space points in R, PR, UR, PAR, and AR states at different values of the interleaflet enthalpic interaction strength constants 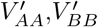, and 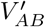 for the 3:1 DPPC-D34 system.

### IX. DOMAIN REGISTRATION HISTOGRAMS FOR THE INTERLEAFLET ENTROPIC STRENGTH PARAMETER

This section shows the histograms showing the populations of R, PR, UR, PAR, and AR states at different values of the interleaflet entropic strength parameter 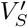 for the systems excluded from the main manuscript in section IV.F, namely the 1:1 DPPC-D23 system in fig.26, the 1:3 DPPC-D23 system in fig.27, the 3:1 DPPC-D23 system in fig.28, the 1:3 DPPC-D34 system in Fig.29, and the 3:1 DPPC-D34 system in Fig.30.

**FIG. 26:**
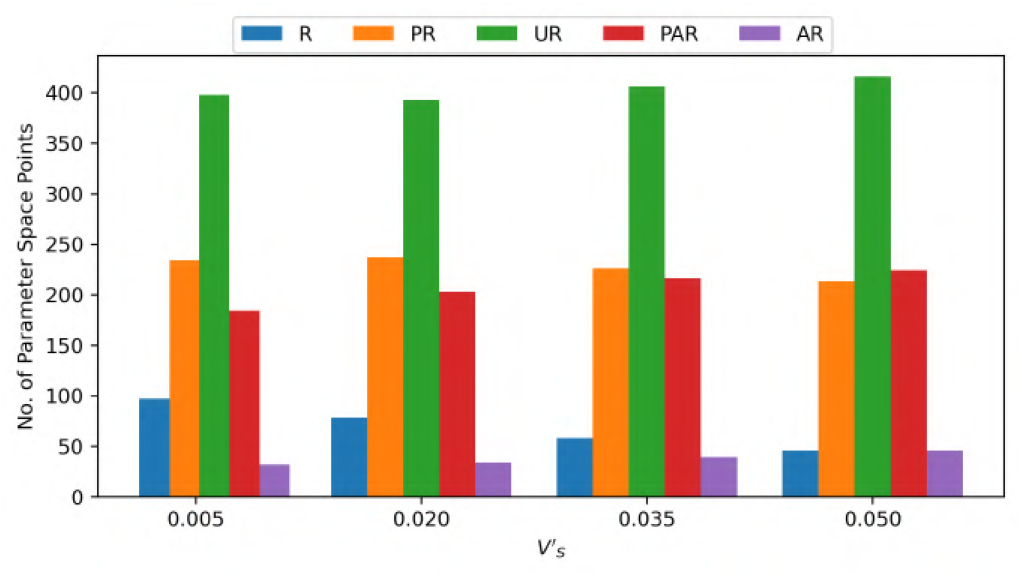
Histograms showing the populations of parameter space points in R, PR, UR, PAR, and AR states at different values of the interleaflet entropic strength contant 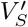 for the 1:1 DPPC-D23 system.

**FIG. 27:**
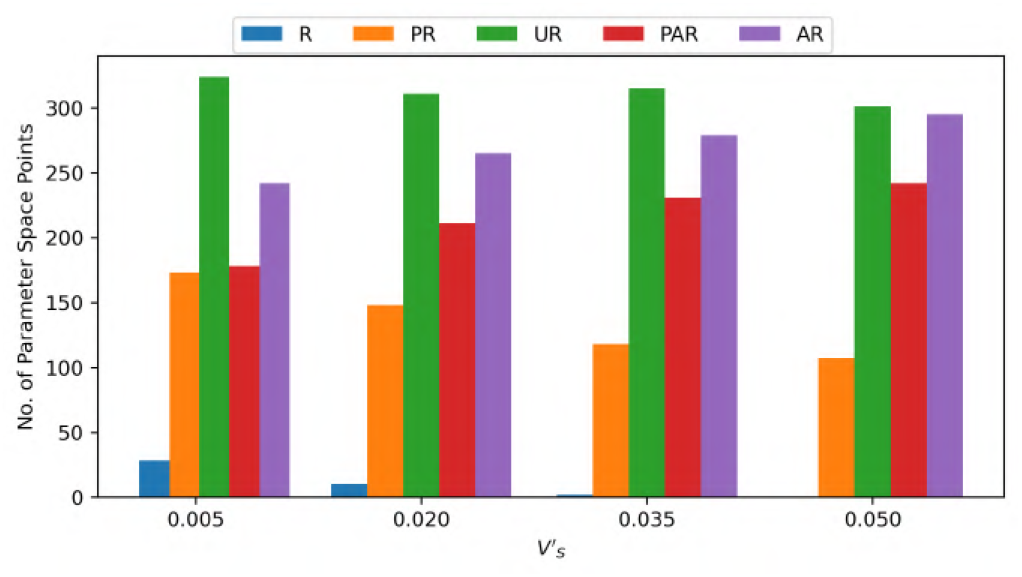
Histograms showing the populations of parameter space points in R, PR, UR, PAR, and AR states at different values of the interleaflet entropic strength contant 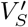 for the 1:3 DPPC-D23 system.

**FIG. 28:**
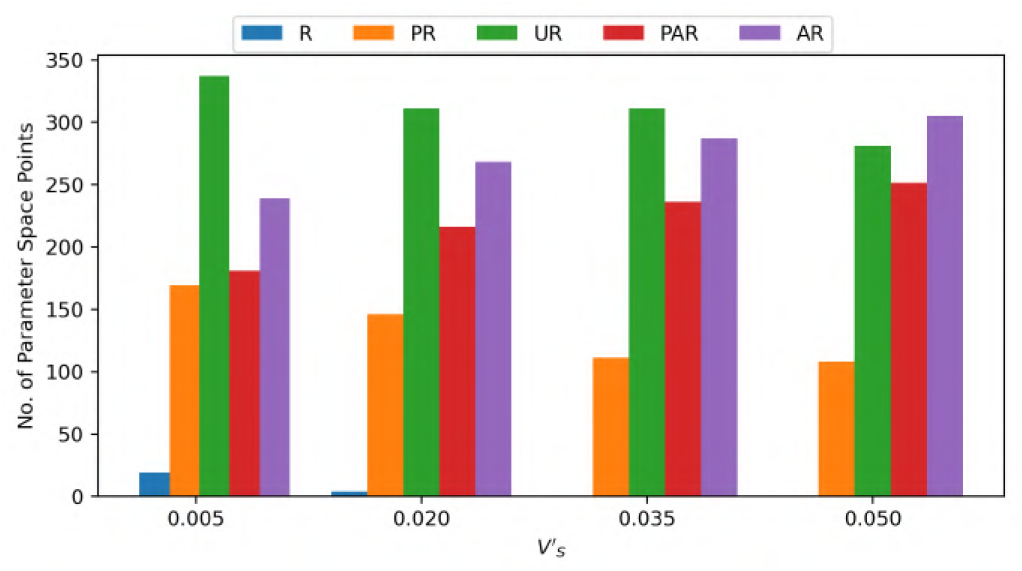
Histograms showing the populations of parameter space points in R, PR, UR, PAR, and AR states at different values of the interleaflet entropic strength contant 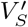 for the 3:1 DPPC-D23 system.

**FIG. 29:**
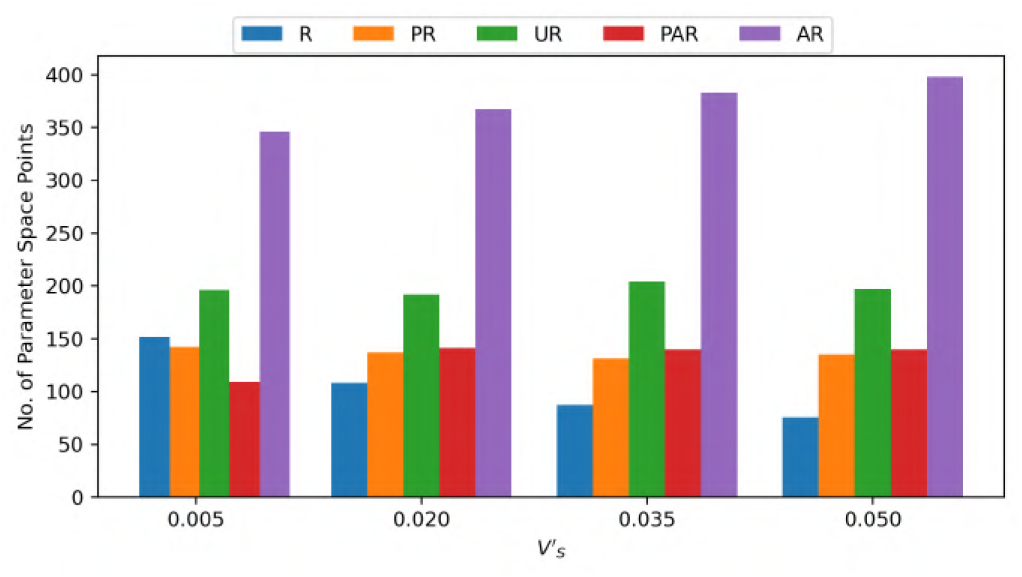
Histograms showing the populations of parameter space points in R, PR, UR, PAR, and AR states at different values of the interleaflet entropic strength contant 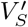 for the 1:3 DPPC-D34 system.

**FIG. 30:**
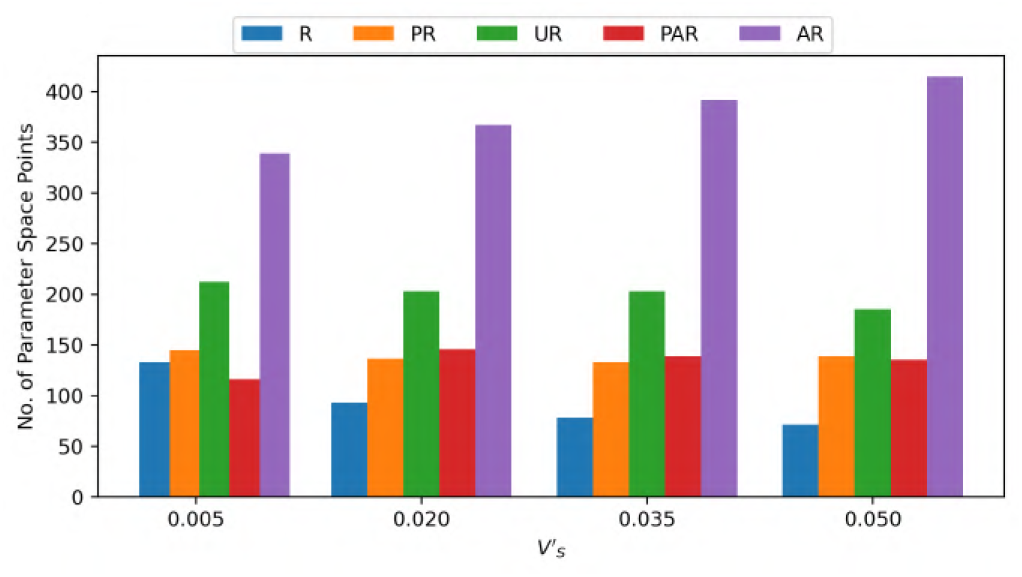
Histograms showing the populations of parameter space points in R, PR, UR, PAR, and AR states at different values of the interleaflet entropic strength contant 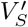 for the 3:1 DPPC-D34 system.

### X. ADDITIONAL POINTS IN PARAMETER SPACE DISPLAYING STABLE NANODOMAINS

Some additional points showing formation of stable nanodomains are shown in fig.31,fig.32, and fig.33. The blue coloured squares represent DPPC and the green squares represent D34 in the following images. In the differences map, yellow squares show mismatched/anti-registered sites and green squares represent registered sites.

**FIG. 31:**
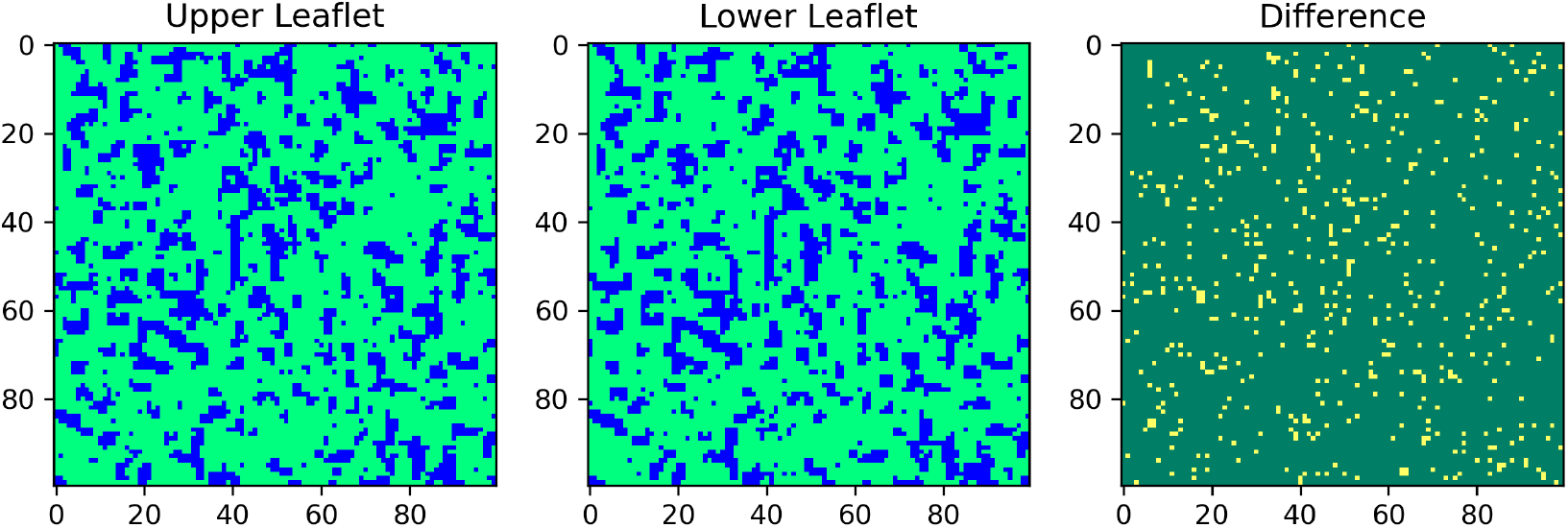
Configuration near the end of the 10^9^ step extended simulation showing the observed nanodomains for the 1:3 DPPC-D34 system at parameter space point 0-0-0-1-1-0-3-1-0.

**FIG. 32:**
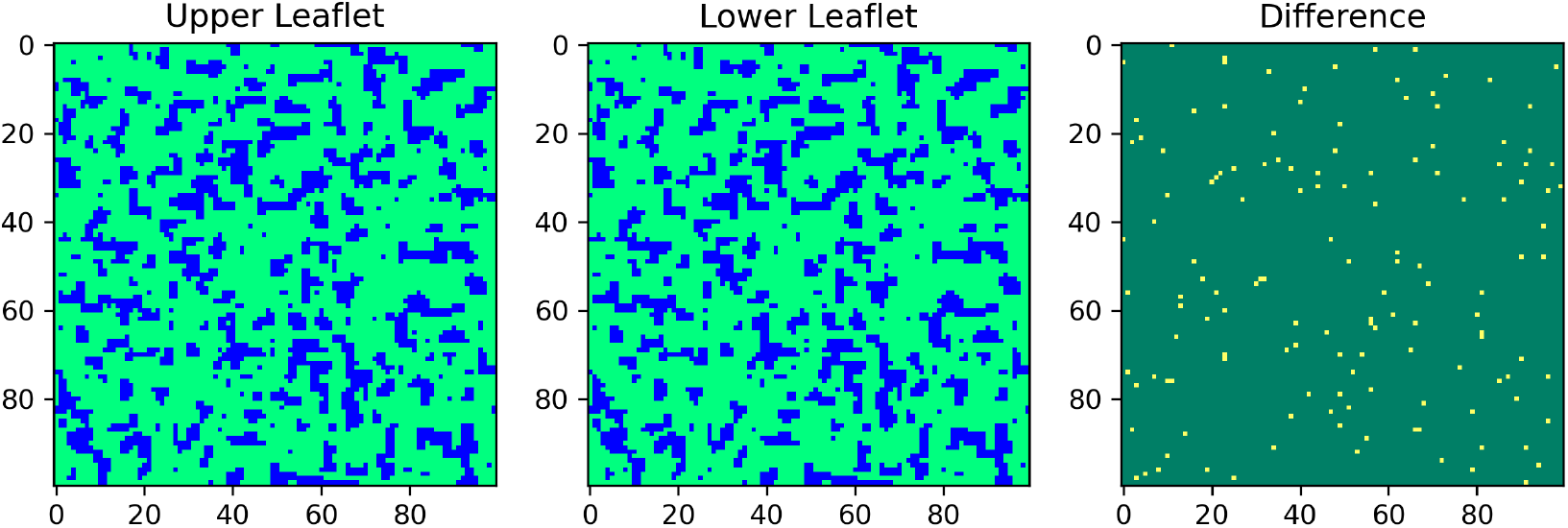
Configuration near the end of the 10^9^ step extended simulation showing the observed nanodomains for the 1:3 DPPC-D34 system at parameter space point 0-0-0-2-1-0-3-1-0.

**FIG. 33:**
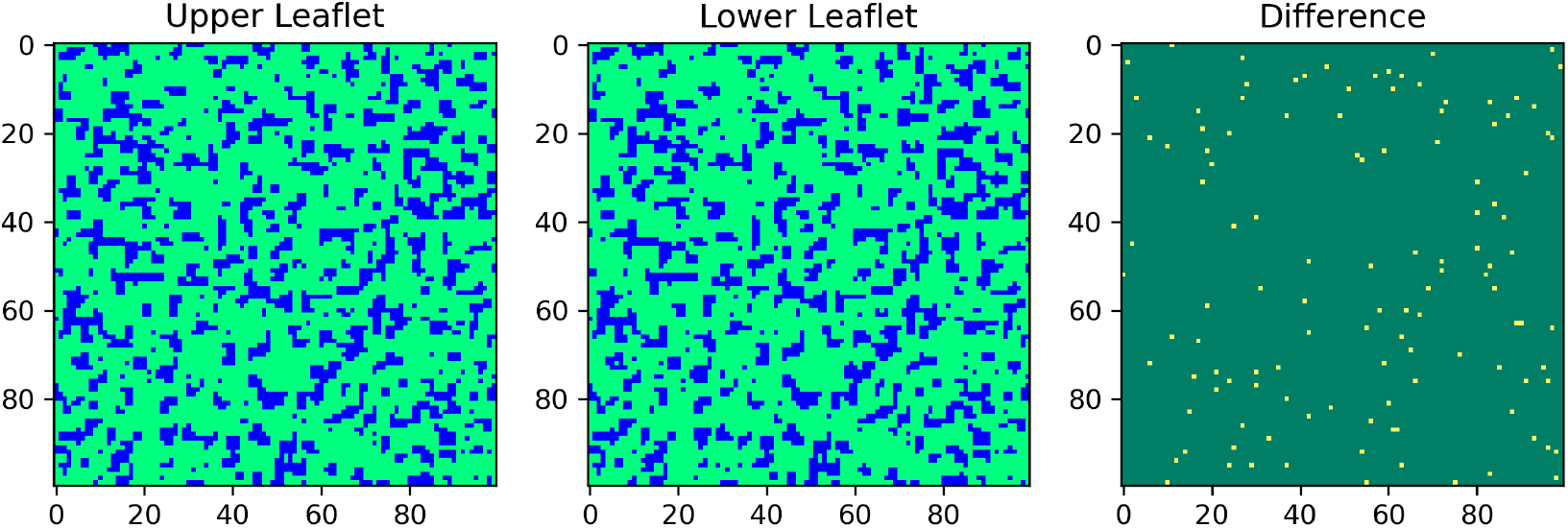
Configuration near the end of the 10^9^ step extended simulation showing the observed nanodomains for the 1:3 DPPC-D34 system at parameter space point 0-0-0-2-2-0-3-0-0.

### XI. PARAMETER SPACE PLOTS FOR PHASE SEPARATION

The data from the classification analysis tools was also used to plot parameter space plots with the seven varying parameters for a 2D ”phase diagram”-like representation for the phase separation data.

The following parameter space plots from Fig.34 to Fig.39, one for each of the 6 systems, have an outer pair of axes, the horizontal axis representing 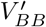 and the vertical one representing 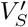. Nested within this pair of axes are 12 subplots and each of the 12 subplots have a pair of axes such that the horizontal one represents 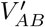 and the vertical one represents 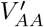. Within each of these subplots, there are 9 3D plots which have three axes representing the parameters *V_AA_, V_BB_*, and *V_AB_*. This effectively shows the variation of classification state with variation in the 7 changing parameters in a 2D plot.

### XII. PARAMETER SPACE PLOTS FOR DOMAIN REGISTRATION

Plots with the same structure as those above were made for the data from the domain registration analysis as well, leading to the following parameter space plots from Fig.40 to Fig.45.

---

**FIG. 34:**
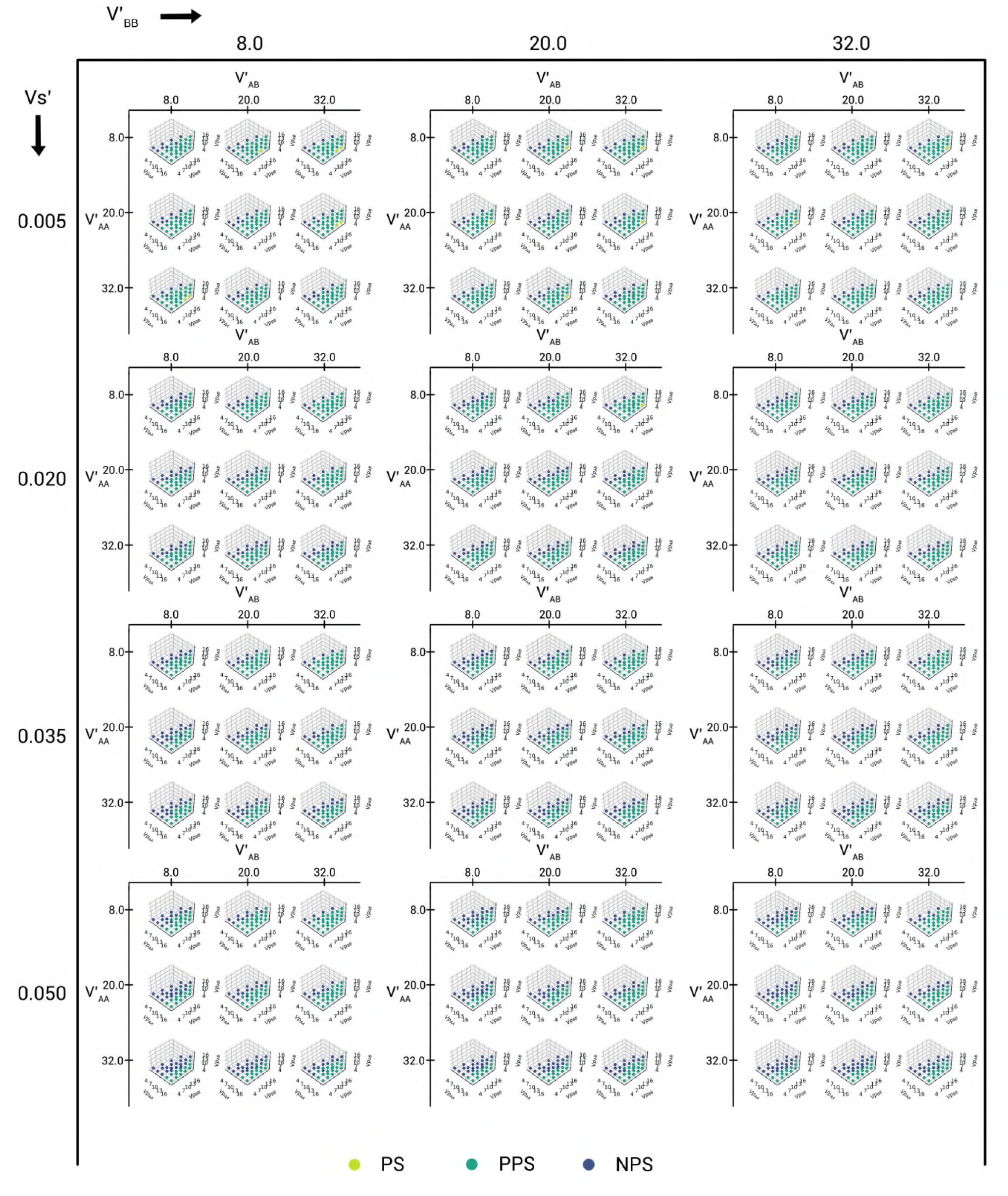
The plot of phase separation classification for all of the parameter space points for the 1:1 DPPC-D23 system, assigned using the domain size distribution analysis.

**FIG. 35:**
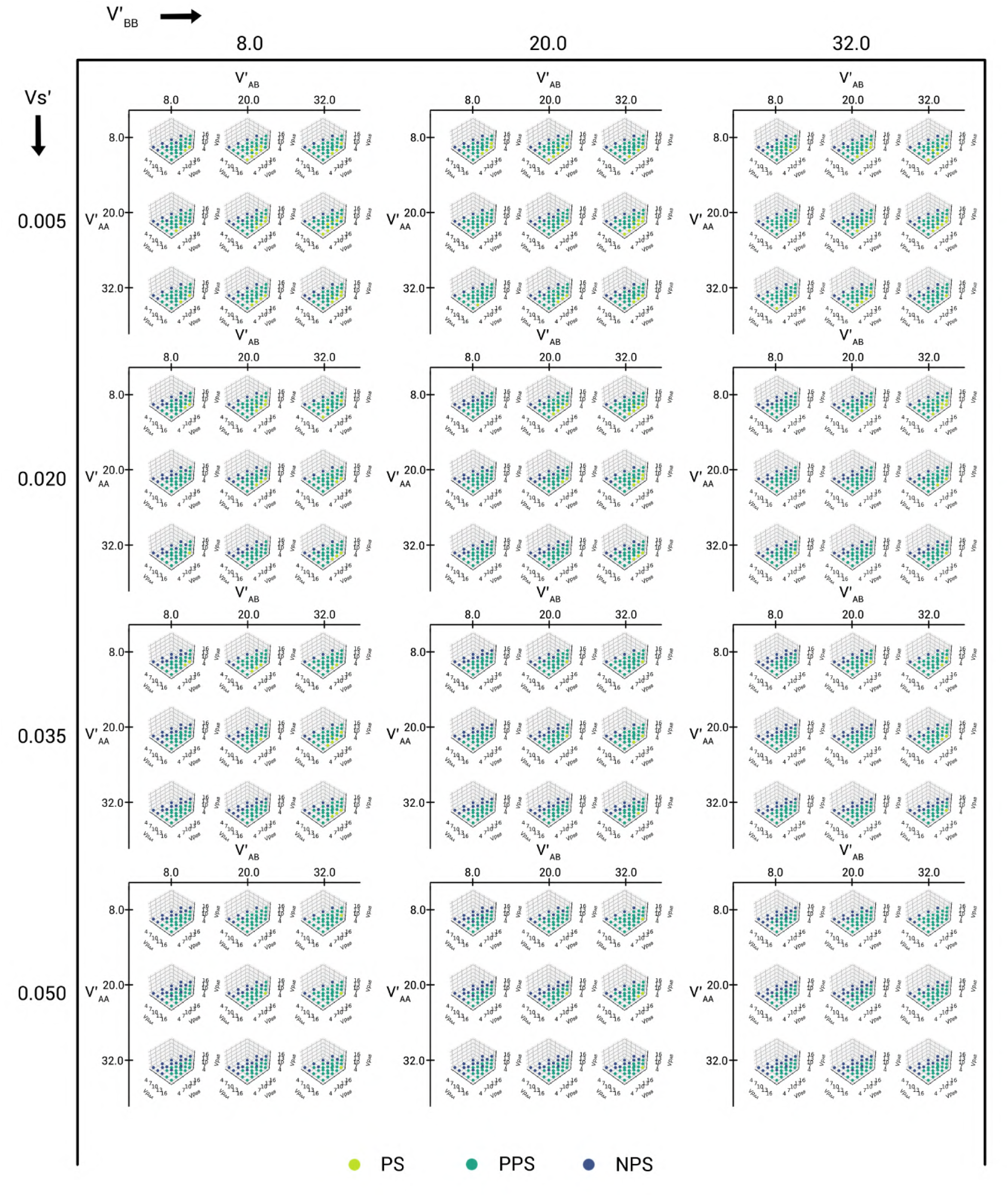
The plot of phase separation classification for all of the parameter space points for the 1:1 DPPC-D34 system, assigned using the domain size distribution analysis.

**FIG. 36:**
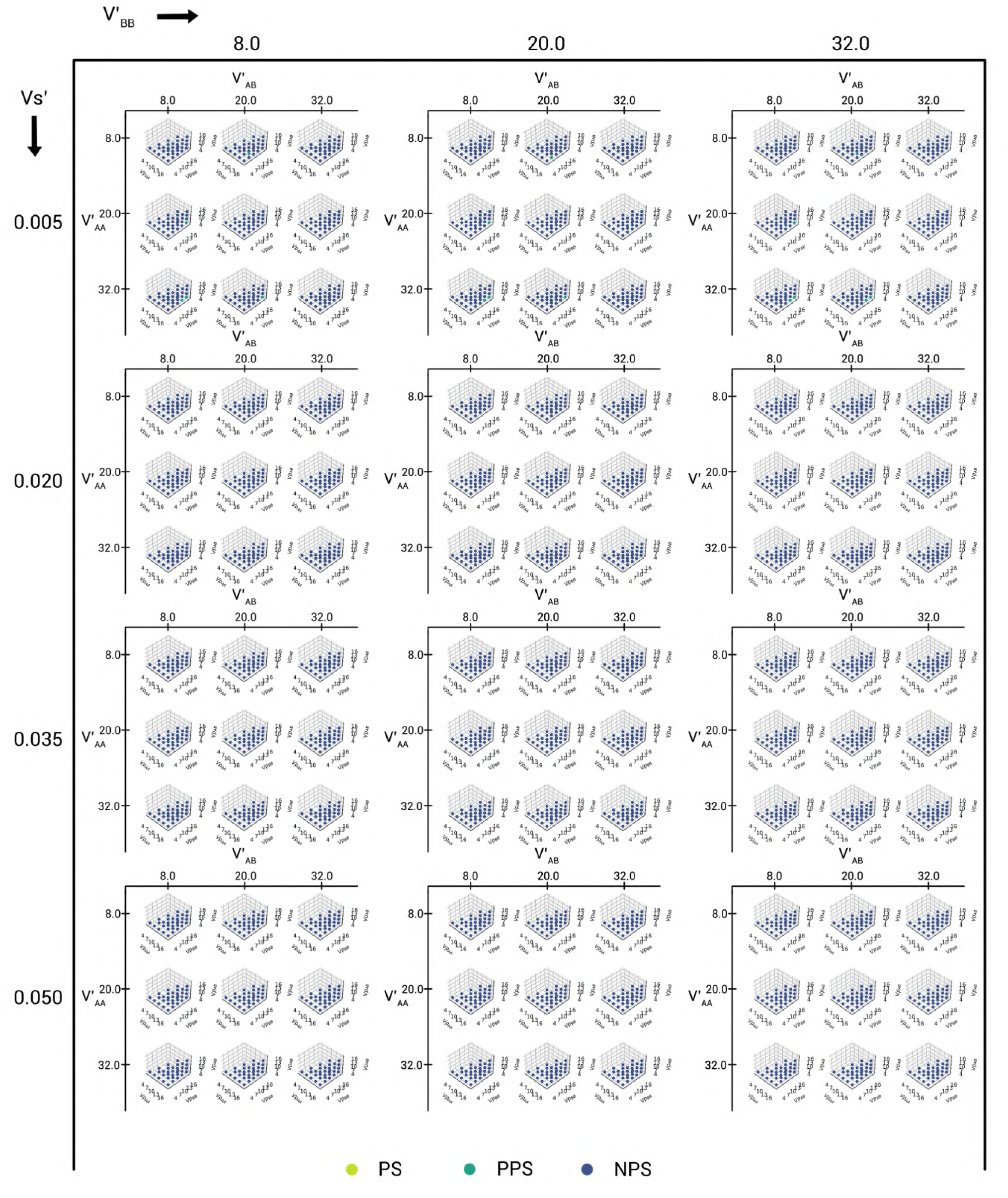
The plot of phase separation classification for all of the parameter space points for the 1:3 DPPC-D23 system, assigned using the domain size distribution analysis.

**FIG. 37:**
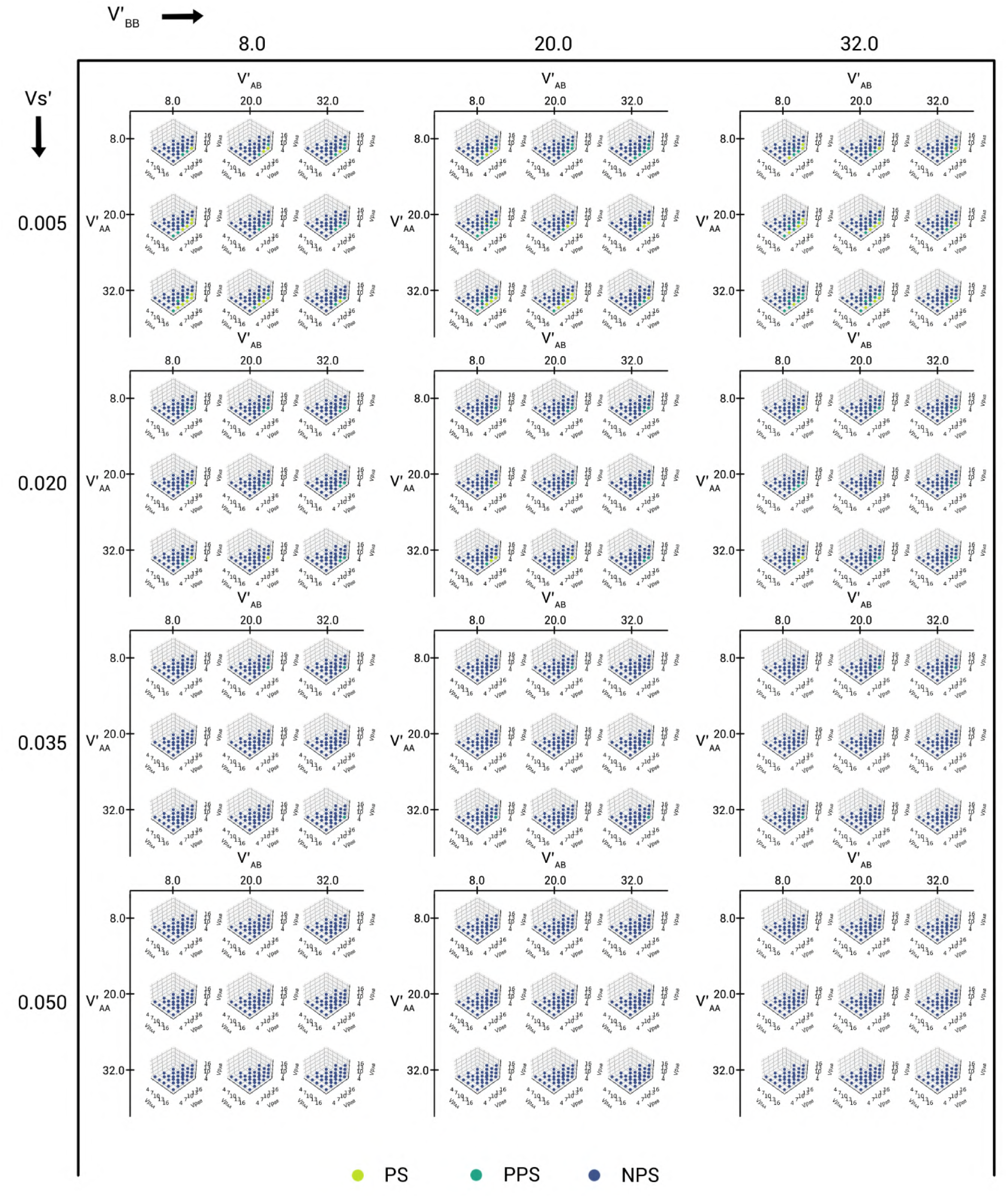
The plot of phase separation classification for all of the parameter space points for the 1:3 DPPC-D34 system, assigned using the domain size distribution analysis.

**FIG. 38:**
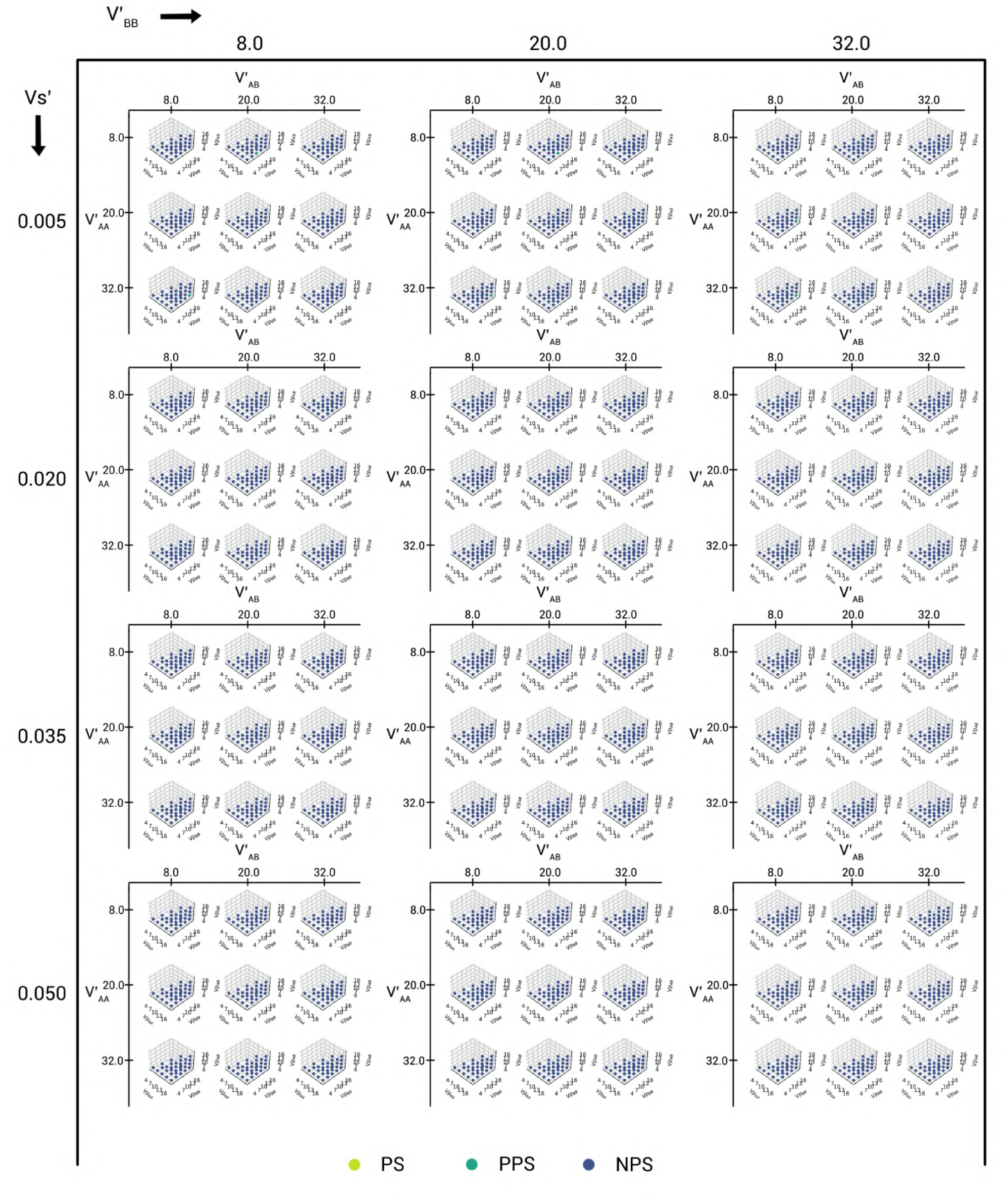
The plot of phase separation classification for all of the parameter space points for the 3:1 DPPC-D23 system, assigned using the domain size distribution analysis.

**FIG. 39:**
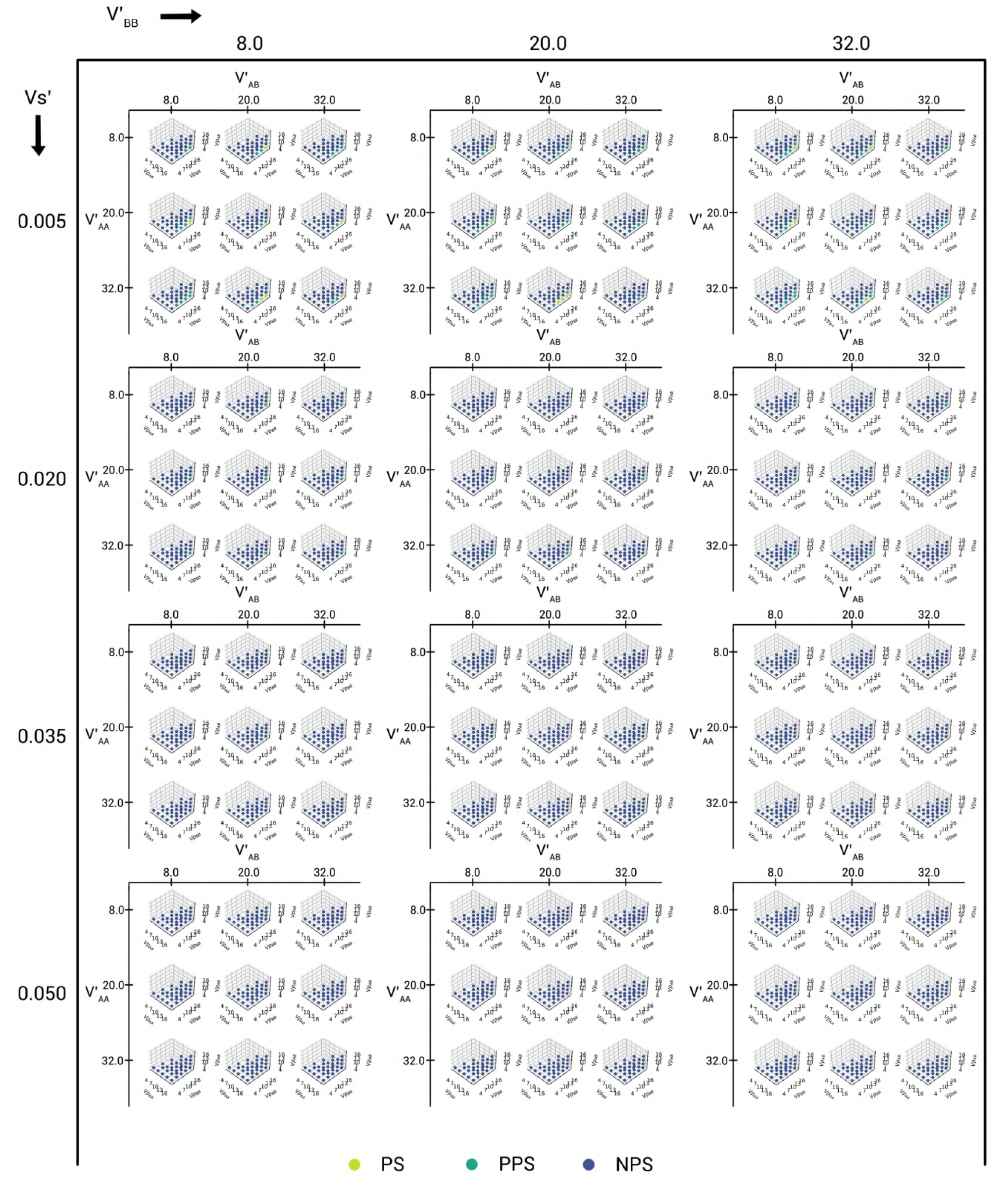
The plot of phase separation classification for all of the parameter space points for the 3:1 DPPC-D34 system, assigned using the domain size distribution analysis.

**FIG. 40:**
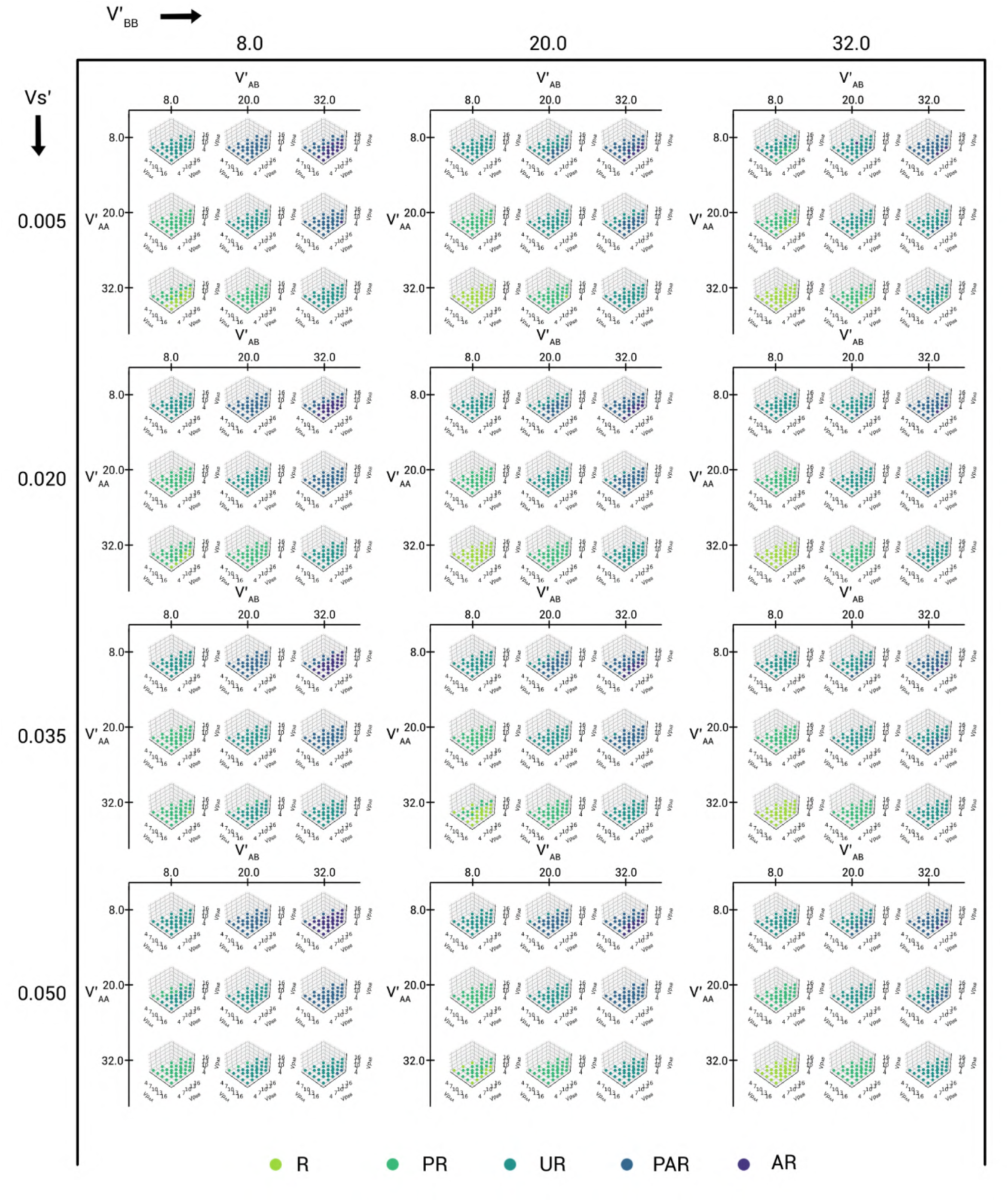
The plot of domain registration behaviour for all of the parameter space points for the 1:1 DPPC-D23 system, assigned using the KL Divergence analysis.

**FIG. 41:**
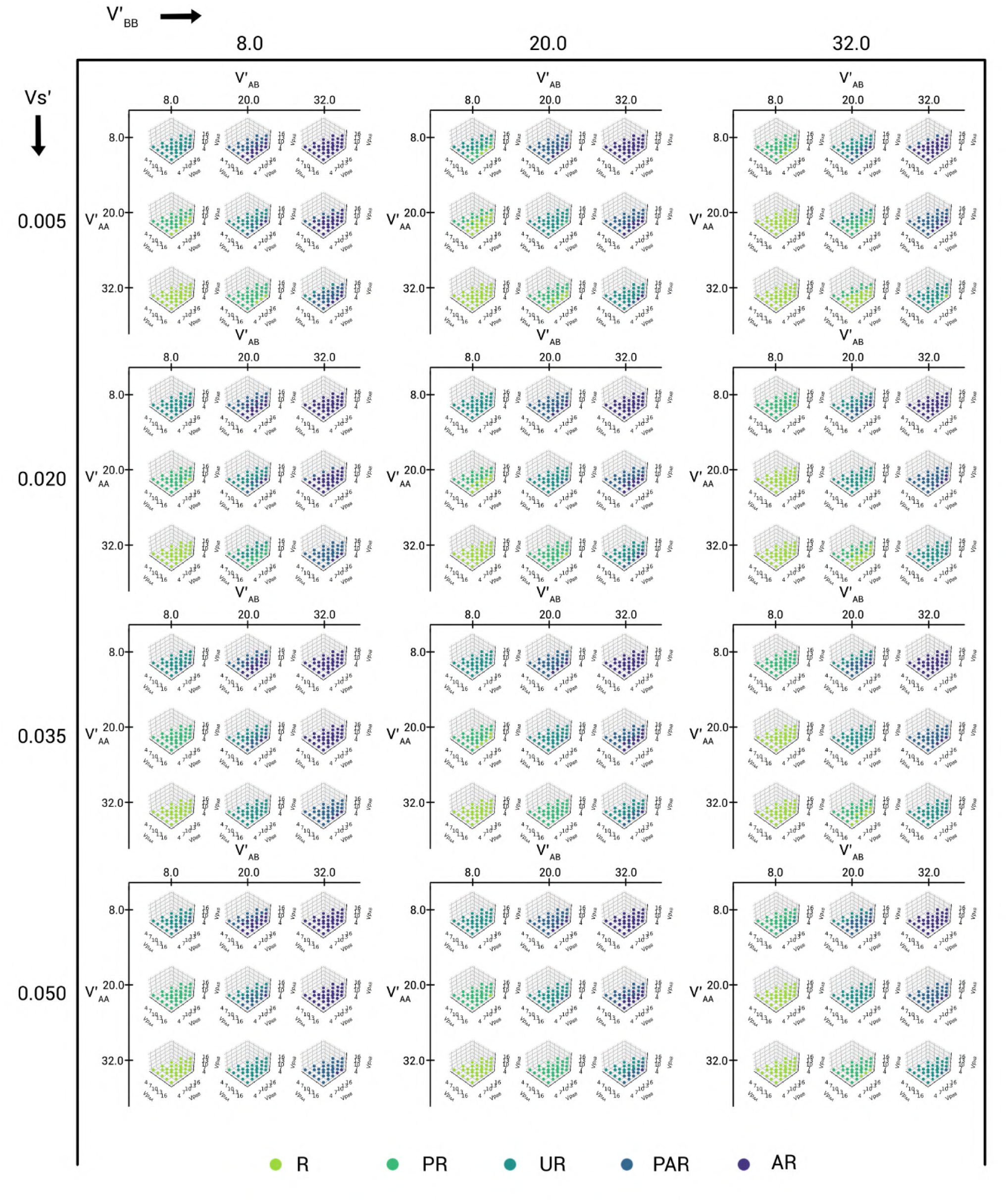
The plot of domain registration behaviour for all of the parameter space points for the 1:1 DPPC-D34 system, assigned using the KL Divergence analysis.

**FIG. 42:**
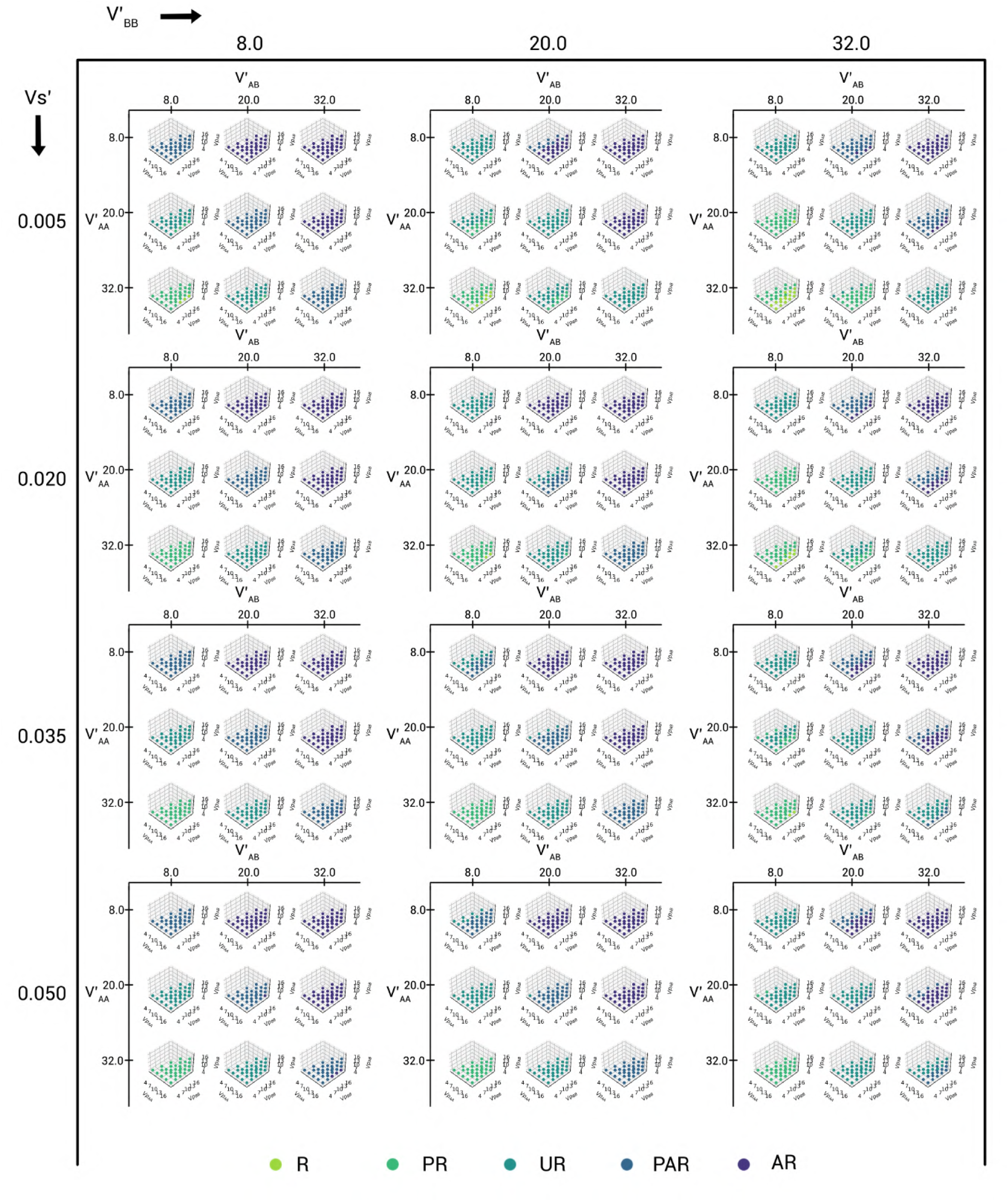
The plot of domain registration behaviour for all of the parameter space points for the 1:3 DPPC-D23 system, assigned using the KL Divergence analysis.

**FIG. 43:**
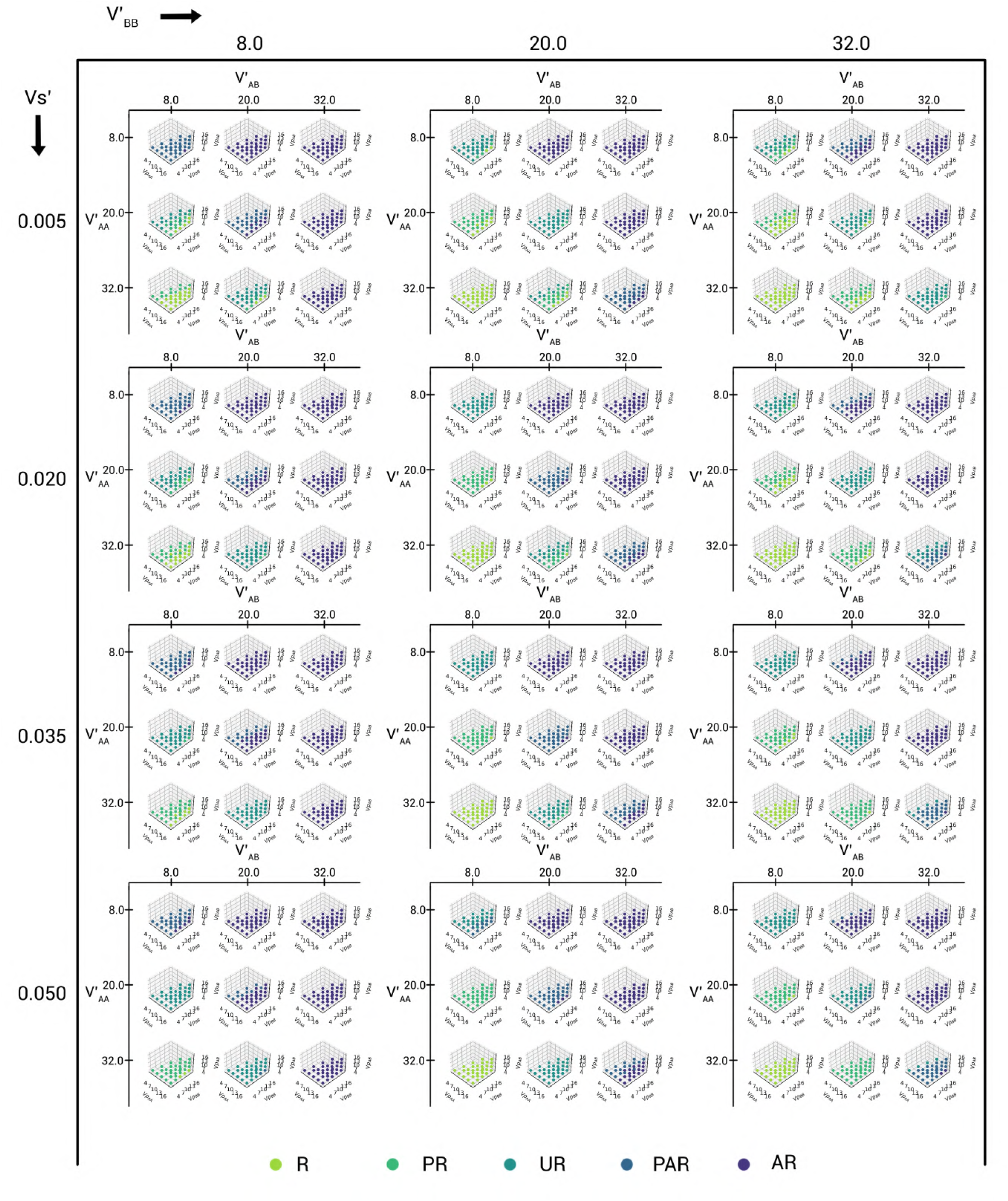
The plot of domain registration behaviour for all of the parameter space points for the 1:3 DPPC-D34 system, assigned using the KL Divergence analysis.

**FIG. 44:**
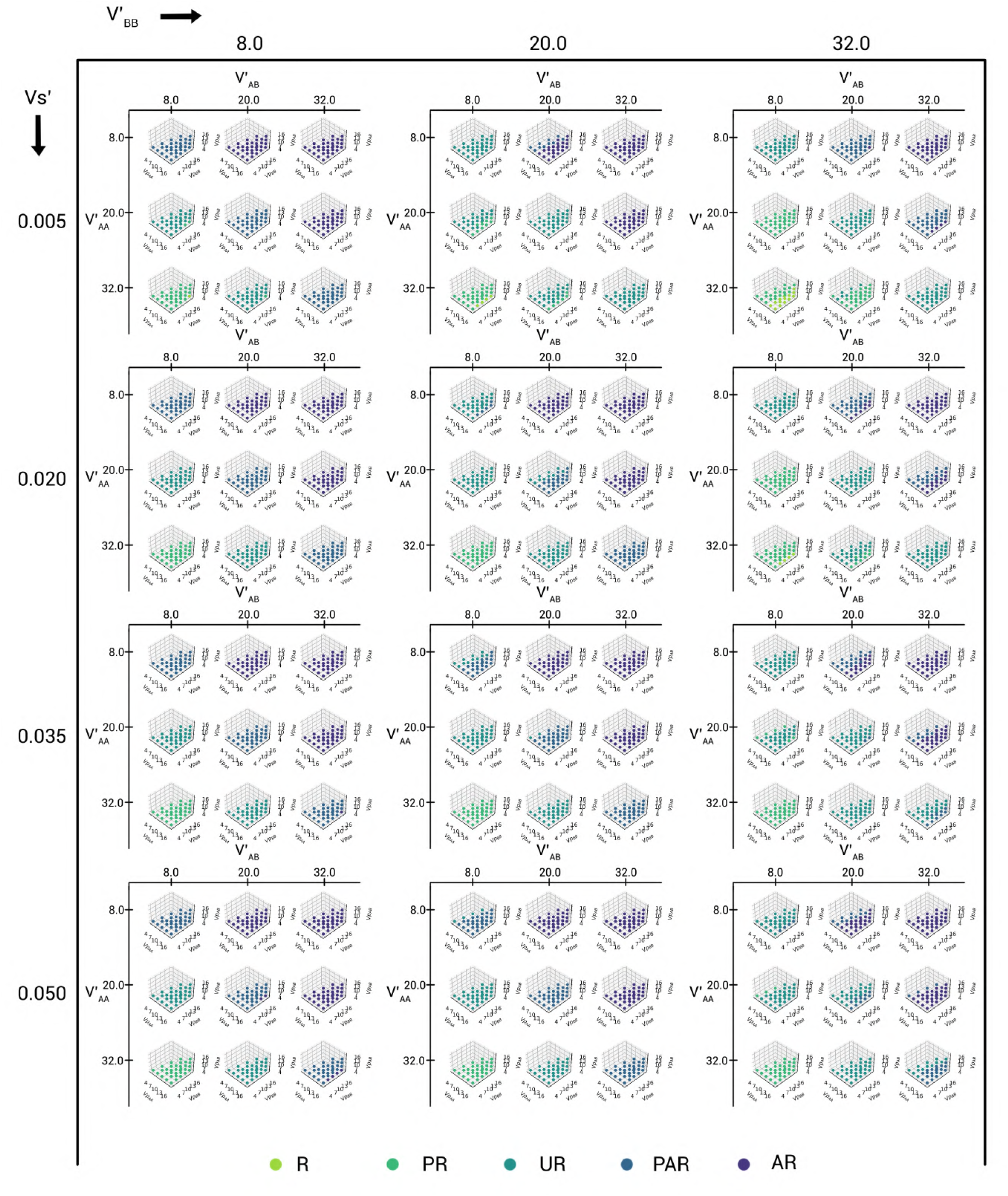
The plot of domain registration behaviour for all of the parameter space points for the 3:1 DPPC-D23 system, assigned using the KL Divergence analysis.

**FIG. 45:**
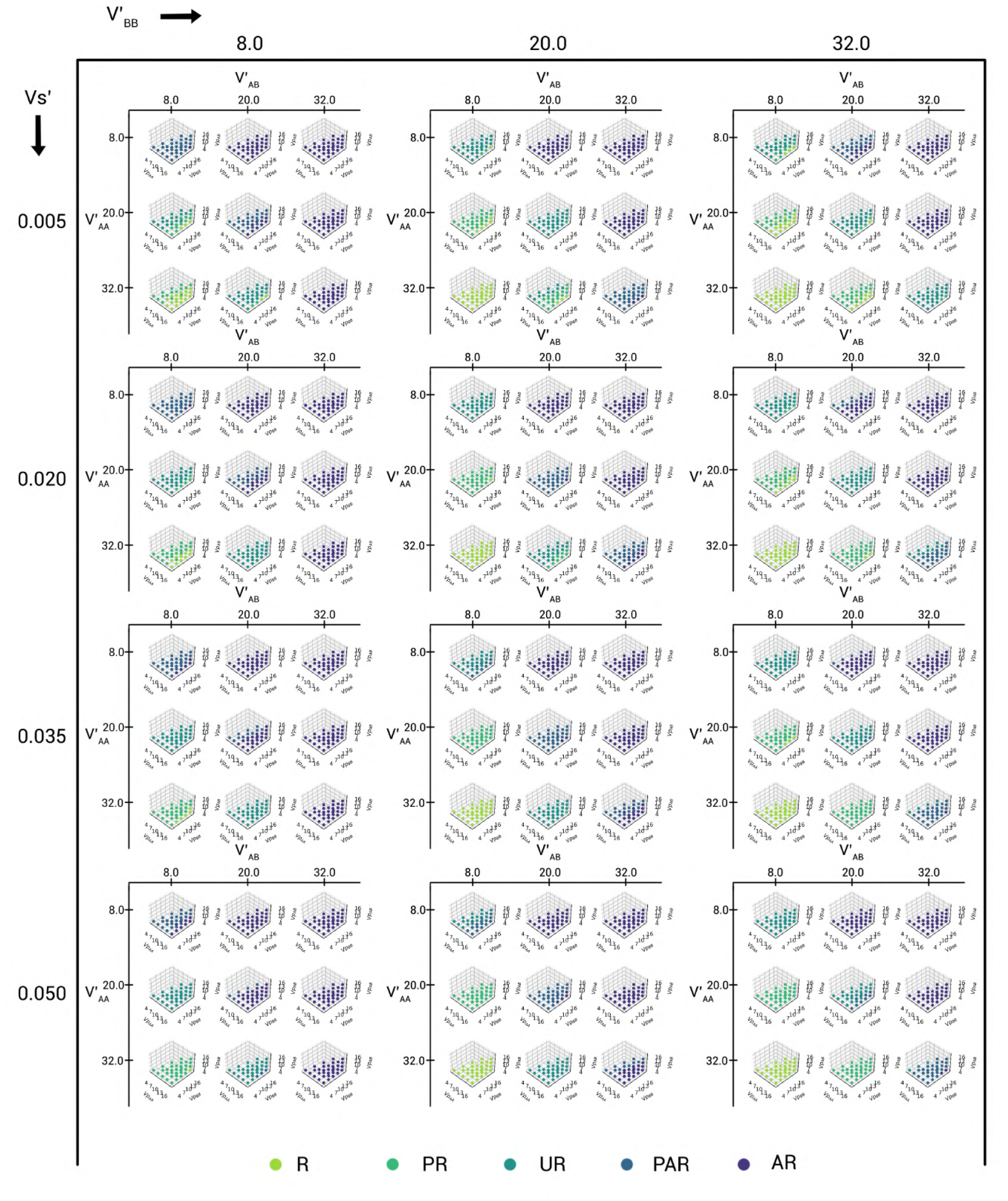
The plot of domain registration behaviour for all of the parameter space points for the 3:1 DPPC-D34 system, assigned using the KL Divergence analysis.

